# Ambiguity of protein abbreviation PDK1: Error categorization and propagation analysis

**DOI:** 10.64898/2025.12.01.691077

**Authors:** Blaž Kociper, Alja Nike Kastrin, Katarina Miš, Pablo M Garcia-Roves, Alexander V Chibalin, Arild C Rustan, Erich Gnaiger, Klemen Dolinar, Senja Pollak, Bojan Cestnik, Andrej Kastrin, Nada Lavrač, Boshko Koloski, Sergej Pirkmajer

## Abstract

Despite peer review, hundreds of biomedical articles contain errors due to identical abbreviations that refer to different proteins. This includes PDK1, denoting both pyruvate dehydrogenase kinase 1 and 3-phosphoinositide-dependent protein kinase 1. Our analysis shows that one-out-of-five articles with the term PDK1 on PubMed, published in 2019–2026, state incorrect antibodies, refer to incorrect sequences for PCR, gene silencing, or plasmids, merge the properties of the two proteins, or incorrectly cite the other protein. Confusion extends to websites of biotechnology providers, where PDK1 antibodies and recombinant proteins were misattributed. To mitigate PDK1 abbreviation misuse, we recommend using unique protein abbreviations, clear antibody and sequence identification, and implementation of more rigorous peer review processes, supported by our newly developed PDK1-TermTracker citation network visualization and analysis service.

## Main Text

Organizations, such as HUGO Gene Nomenclature Committee (HGNC), address vague designations of genes and proteins by assigning a unique abbreviation to each gene and corresponding protein (*1*). However, for historical reasons (*2*), many other abbreviations remain frequently in active use. As if it were not confusing enough to have several different abbreviations for one protein, the same abbreviation can be used for two or more different proteins. Consequently, from the abbreviation alone, one cannot infer which protein is referred to. As noted recently (*3*), hundreds of biomedical articles may contain significant errors due to protein name confusion, resulting in antibody mix-up^1^. The situation is similar for PDK1 (*4, 5*), which may refer to at least two different proteins: pyruvate dehydrogenase kinase 1 (UniProt: Q15118) and 3-phosphoinositide-dependent protein kinase 1 (UniProt: O15530).

PDK1 conceptual meaning A: Pyruvate dehydrogenase kinase 1 (HGNC symbol PDK1) is one of the four isoforms of pyruvate dehydrogenase kinases that inhibit pyruvate dehydrogenase (PDH) by phosphorylating it and thereby shifting glucose metabolism from oxidation to anaerobic glycolysis (**Figure 1**, **right**) (*4*). The discovery of pyruvate dehydrogenase kinase predates the discovery of 3-phosphoinositide-dependent protein kinase 1. The existence of a kinase that inhibits pyruvate dehydrogenase was described already in 1969 (*6*). At first, pyruvate dehydrogenase kinase was abbreviated as PDH kinase (*7*) and later as PDHK (*8*). The oldest reference we found to abbreviate pyruvate dehydrogenase kinase as PDK dates back to 1994 (*9*).

**Figure 1.**
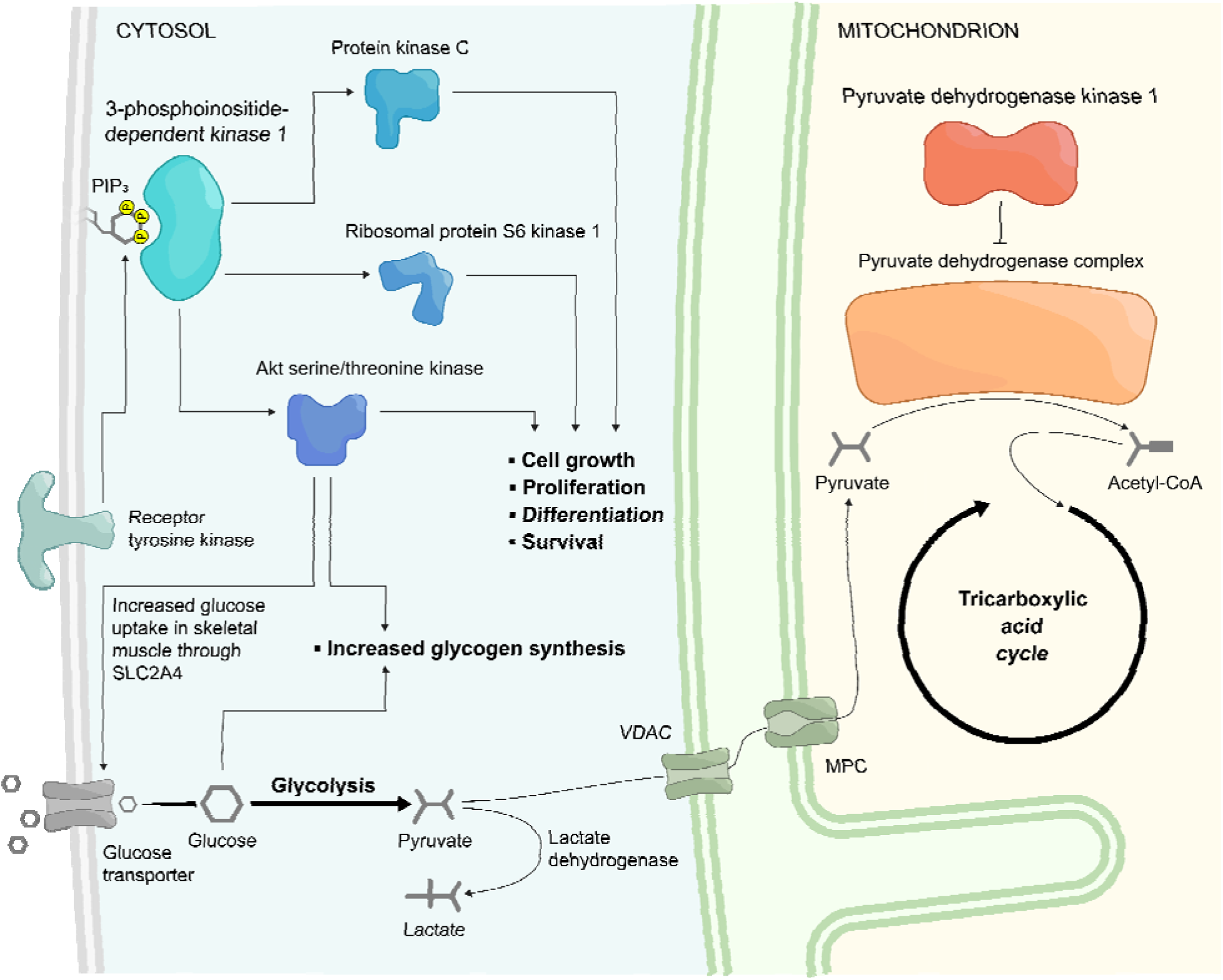
Scheme of pyruvate dehydrogenase kinase 1 (concept A, right) and 3- phosphoinositide-dependent protein kinase 1 (concept B, left). Pyruvate dehydrogenase kinase 1 inhibits pyruvate dehydrogenase complex through phosphorylation, thus redirecting glucose metabolism from the tricarboxylic acid cycle towards glycolysis and lactate formation (*4*). 3-phosphoinositide-dependent protein kinase 1 is activated upon ligand binding, such as insulin, to receptor tyrosine kinases. It plays an important role in regulating metabolic flux and promoting cell growth, proliferation, differentiation, and survival (*11*). VDAC = voltage-dependent anion channel, MPC = mitochondrial pyruvate carrier. The scheme and description simplify certain aspects for clarity.

PDK1 conceptual meaning B: 3-phosphoinositide-dependent protein kinase 1 (HGNC symbol PDPK1) is a kinase activated by phosphatidylinositol (3,4,5)-trisphosphate (PIP_3_). PIP_3_ is generated by phosphoinositide 3-kinases upon ligand binding to tyrosine kinase receptors. Once activated, 3-phosphoinositide-dependent protein kinase 1 is involved in the activation of AKT serine/threonine kinase, protein kinase C, and ribosomal protein S6 kinase B1 (*10*), which are part of an important signaling network involved in the regulation of energy metabolism, cell growth, proliferation, differentiation, and survival (**Figure 1**, **left**) (*11*). 3-phosphoinositide-dependent protein kinase 1 was first described in 1997 and was abbreviated in that very article as PDK1 (*5*).

The ambiguity of the term PDK1 has been noted before. Some authors observed that the term PDK1, referring to 3-phosphoinositide-dependent protein kinase 1, could lead to misidentification with pyruvate dehydrogenase kinase 1 (*12*). However, despite this early warning, both kinases are still persistently abbreviated as PDK1. We hypothesized that errors due to the ambiguous PDK1 abbreviation might have spread throughout scientific literature. Therefore, we conducted an in-depth review of PDK1-related articles and classified the types of most common errors. We also performed citation network analysis and network visualization, to track PDK1-induced error propagation in PubMed articles. To this end, we developed and openly released the PDK1- TermTracker service, supporting PDK1 error detection and propagation tracking in PubMed. In addition to helping visualize PDK1 error propagation in the existing set of 1,169 manually annotated articles, the Term Tracker actively searches for newly published PDK1-related articles on PubMed and suggests PDK1 error labeling of articles using a Large Language Model (LLM) classifier, while leaving the ultimate annotation decision to the expert. Finally, to evaluate potential ambiguities in protein target information from biotechnology providers and vendors, we experimentally assessed the specificity of selected PDK1 antibodies.

## Results

### Prevalence of errors due to the ambiguous terminology in PDK1 articles

To assess the prevalence of errors associated with the PDK1 abbreviation, we conducted a PubMed search for the keyword “PDK1” between January 1, 2019 and June 22, 2026. The search resulted in 1,248 records, which included the original articles (some of which were later corrected by the authors^2^), as well as some articles retracted by the authors^3^. Replacing original articles with their corrected versions, and the elimination of irrelevant articles^4^ resulted in a final set of 1,169 articles, which were manually reviewed and annotated. Out of these, 262 (22.4 %) contained PDK1-related errors. The entire list of articles used in the analysis is provided in **Supplement S1**.

The search, selection and manual annotation of these PubMed articles was performed by the biomedical expert (author Kociper). In addition, to assess the quality of manual annotations, an independent biomedical expert (author Dolinar) manually annotated a random sample of 200 articles, enabling us to assess the quality of expert annotations. Krippendorf alpha annotation agreement 0.806 was achieved, strengthening the credibility of the original expert assessment. For more detail see Section Methods and **Supplement S2**.

### Types of errors in PDK1 scientific articles

The identified articles exhibited several types of errors, which we have classified into four categories. Note that the references listed below include the articles from the 1,169 publications found via the PDK1 term search in the 2019–2026 period; for more details, see the Methods section.

Error type 1: Incorrect antibody (referred to as errors Ã_1_ or B_1_ ). Error Ã_1_ refers to incorrect use of PDK1 abbreviation, where PDK1 is erroneously presented as pyruvate dehydrogenase kinase 1 (concept A), but the antibody context corresponds to 3-phosphoinositide-dependent protein kinase 1 (concept B). On the other hand, error B_1_ refers to incorrect use of PDK1 abbreviation, where PDK1 is erroneously presented as 3-phosphoinositide-dependent protein kinase 1 (concept B), but the antibody context corresponds to pyruvate dehydrogenase kinase 1 (concept A). Such antibodies may be reported for use in immunoblotting or immunohistochemistry (**Supplement S2**). The issue is further aggravated by the fact that some articles do not provide catalogue numbers of antibodies, making verification impossible (*13-15*), which reduces the transparency and reproducibility of such reports.

Error type 2: Incorrect sequence (referred to as errors Ã_2_ or B̃_2_). This applies to articles where pyruvate dehydrogenase kinase 1 (concept A) is discussed, but the stated nucleotide sequence corresponds to 3-phosphoinositide-dependent protein kinase 1 (referred to as error Ã_2_), and *vice versa* (referred to as error B _2_). These sequences were stated to be used in PCR reactions, gene silencing (e.g., siRNA/shRNA), or plasmid construction (**Supplement S2**). As with antibodies, some studies do not disclose the sequences used (*13, 16, 17*), thereby reducing the transparency and reproducibility of these articles.

Error type 3: Merging of the two proteins (referred to as error M). This error happens when an article repeatedly fails to distinguish between pyruvate dehydrogenase kinase 1 (concept A) and 3-phosphoinositide-dependent protein kinase 1 (concept B) and describes them as if both were the same protein. A common indicator of this error is the inclusion of at least two citations for each of the kinases, both of which are used in the context of the same protein. This creates a false impression that a single protein performs the function of both pyruvate dehydrogenase kinase 1 and 3-phosphoinositide-dependent protein kinase 1 (**Supplement S2**).

Error type 4: Citation related to the other protein (minor single citation errors Ã_C_ or B_C_ ). In this type of error, an article focused on pyruvate dehydrogenase kinase 1 (concept A) incorrectly cites a single time a study discussing 3-phosphoinositide-dependent protein kinase 1, as indicated by clearly explained abbreviation or unique characteristic (error Ã_C_), or *vice versa* (error B _C_). Such misattributions often appear in discussions of disease involvement, metabolic functions, or signaling pathways (**Supplement S2**).

To assess the trend of growing numbers of main error types (error type 1, 2 and 3) in the 2019– 2026 period, we plotted the articles through a timeline (**Figure 2**) (**Supplement S3**).

**Figure 2.**
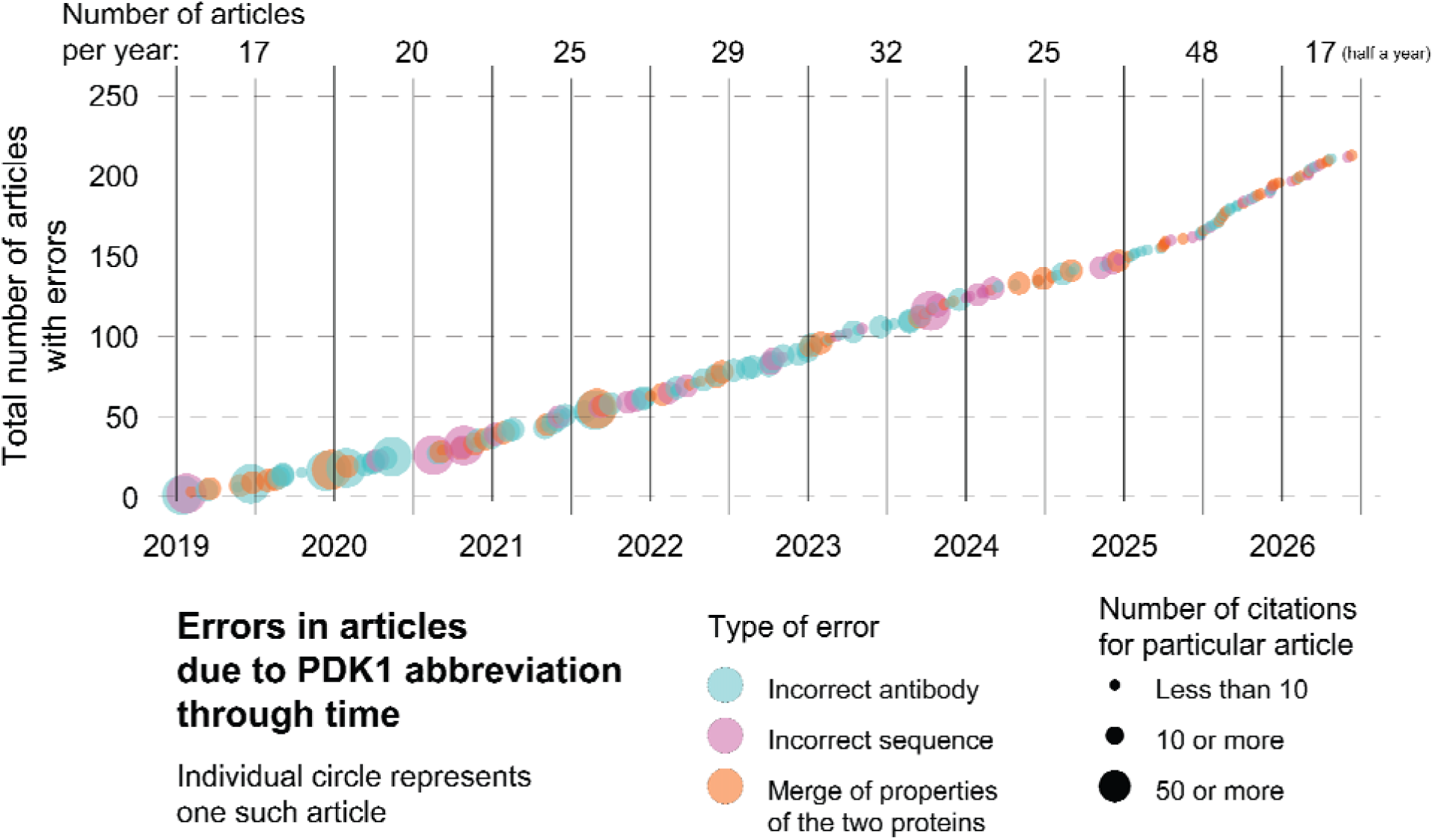
Graph of articles with errors due to interchangeable use of PDK1 from 2019 to mid-2026. Graph shows individual articles with errors represented as circles through January 1, 2019 to June 22, 2026. Circle colors mark the type of error; blue for use of wrong antibody for detection of the wanted protein, pink for stating incorrect sequence for PCR, siRNA, plasmid construction or other, and orange for merging the two proteins together, where discussed protein has characteristics of both. Three different sizes of circles denote the number of times article has been cited with ranges up to 10, up to 50, and beyond. At the very top of the graph stands the number of articles published that year with such errors. As we covered first half of 2026 number 17 stands for the number of articles during these 6 months.

### The PDK1-TermTracker service

To support systematic inspection of how the PDK1 misinterpretation errors are distributed across the literature, we developed the PDK1-TermTracker, an interactive citation network exploration service built around the corpus of manually labeled articles described above^5^. Rather than attempting to automatically detect terminology errors—a task that, as the error categories above illustrate, often requires inspection of antibodies, nucleotide sequences, plasmids, or methods-section detail beyond what titles and abstracts can reveal—the PDK1-TermTracker is designed as an exploratory, expert-support service: it reveals the citation structure and finds category-level patterns, which can guide the domain expert towards articles and citation paths worthy further manual inspection.

The underlying citation network and its construction from the Web of Science (WoS) bibliometric metadata are described in detail in the Methods section. Briefly, each article is represented as a node, and a directed edge from article *u* to article *v* indicates that article *u* cites article *v*. Nodes are annotated with the manually assigned PDK1 category (A, B, Ã , B , or M)^6^ together with bibliographic and citation-count attributes.

To examine whether articles sharing a PDK1 meaning or error category also share citation relationships, node positions in the visualization in **Figure 3**^7^ were derived directly from the topology of this citation network. The figure shows the resulting network for our manually labeled corpus. Articles correctly discussing pyruvate dehydrogenase kinase 1 (A) and 3-phosphoinositide-dependent protein kinase 1 (B) occupy largely distinct regions of this layout, indicating that articles sharing a PDK1 meaning tend to cite, or be cited by, similar sets of other articles. On the other hand, articles in the merged error category (M) are positioned mostly between these two regions, rather than within either one, suggesting that merging errors arise at—and may propagate along—citation paths that bridge the two otherwise separate bodies of literature.

**Figure 3.**
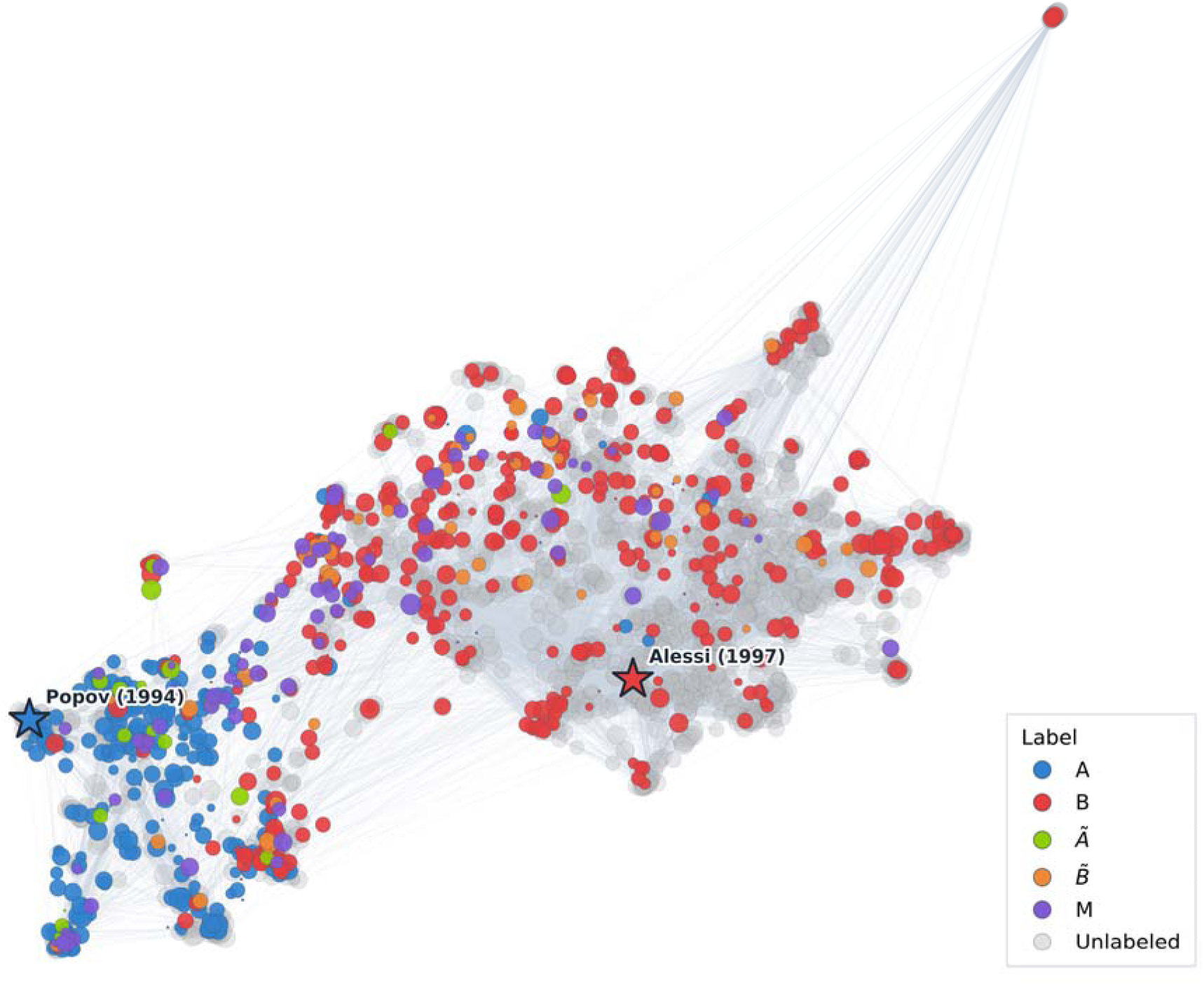
Citation-network landscape of PDK1 ambiguity in the 2019–2026 biomedical literature. A two-dimensional UMAP projection of the citation network separates publications discussing pyruvate dehydrogenase kinase 1 (blue, concept A) from those discussing 3-phosphoinositide-dependent protein kinase 1 (red, concept B). Blue star represents article which first described pyruvate dehydrogenase kinase 1 (*9*), with red star representing article first describing 3-phosphoinositide-dependent protein kinase 1 (*5*). Mixed-use articles (violet, error M) predominantly occupy the transitional region between the two domains, suggesting pathways through which terminological confusion propagates across literature. Green ) and orange ) nodes indicate manually validated reference articles, respectively. Grey nodes represent unlabeled articles. Single node represents one article, node size reflects citation count, and edges indicate citation links between publications.

### PDK1 ambiguities on websites of biotechnology providers and vendors

Multiple biotechnology companies reference both pyruvate dehydrogenase kinase 1 and 3- phosphoinositide-dependent protein kinase 1 for the same antibody or recombinant protein. The errors appear in various forms, including discrepancies between the protein listed on the company website and the protein listed in the datasheet, database links directing to both kinases, listing both proteins as synonyms, and mentioning both on the product page of a single product (for details see **Supplement S4**). For example, anti-PDK1 antibody [EPR19571] (ab202468) by Abcam had target specified as pyruvate dehydrogenase kinase 1, but in the datasheet under Supplementary information the description reads: “*The PDK1 protein also known as 3- phosphoinositide-dependent protein kinase-1 is a kinase enzyme coded by the PDPK1 gene.*” (access date: December 31, 2024). Information in the datasheet has since been corrected referring now to pyruvate dehydrogenase kinase 1 (access date: September 29, 2025). Unfortunately, but as expected for such ambiguous descriptions, various studies used the same antibody for the detection of either pyruvate dehydrogenase kinase 1 (*18-20*) or 3- phosphoinositide-dependent protein kinase 1 (*21-23*).

### Experimental validation of antibodies for detection of pyruvate dehydrogenase kinase 1 or 3-phosphoinositide-dependent protein kinase 1

Given PDK1 ambiguities on websites of biotechnology providers and vendors, we decided to experimentally validate several antibodies. Experiments were conducted by silencing either pyruvate dehydrogenase kinase 1 (A) or 3-phosphoinositide-dependent protein kinase 1 (B) in cultured primary human myotubes (**Figure 4A**, **B**). PDHK1 antibody #3820 offered by vendor Cell Signaling Technology, whose designated target is A (pyruvate dehydrogenase kinase 1), produced a decreased signal in cells with silenced pyruvate dehydrogenase kinase 1 (**Figure 4C**, **D**). PDK1 antibody #3062 offered by the same vendor Cell Signaling Technology for the designated target 3-phosphoinositide-dependent protein kinase 1 (concept B) was used for detection of pyruvate dehydrogenase kinase 1 (concept A) (*24*). Our experimental analysis showed a decreased signal in cells with silenced 3-phosphoinositide-dependent protein kinase 1, but not in those with silenced pyruvate dehydrogenase kinase 1 (**Figure 4C**, **E**).

**Figure 4.**
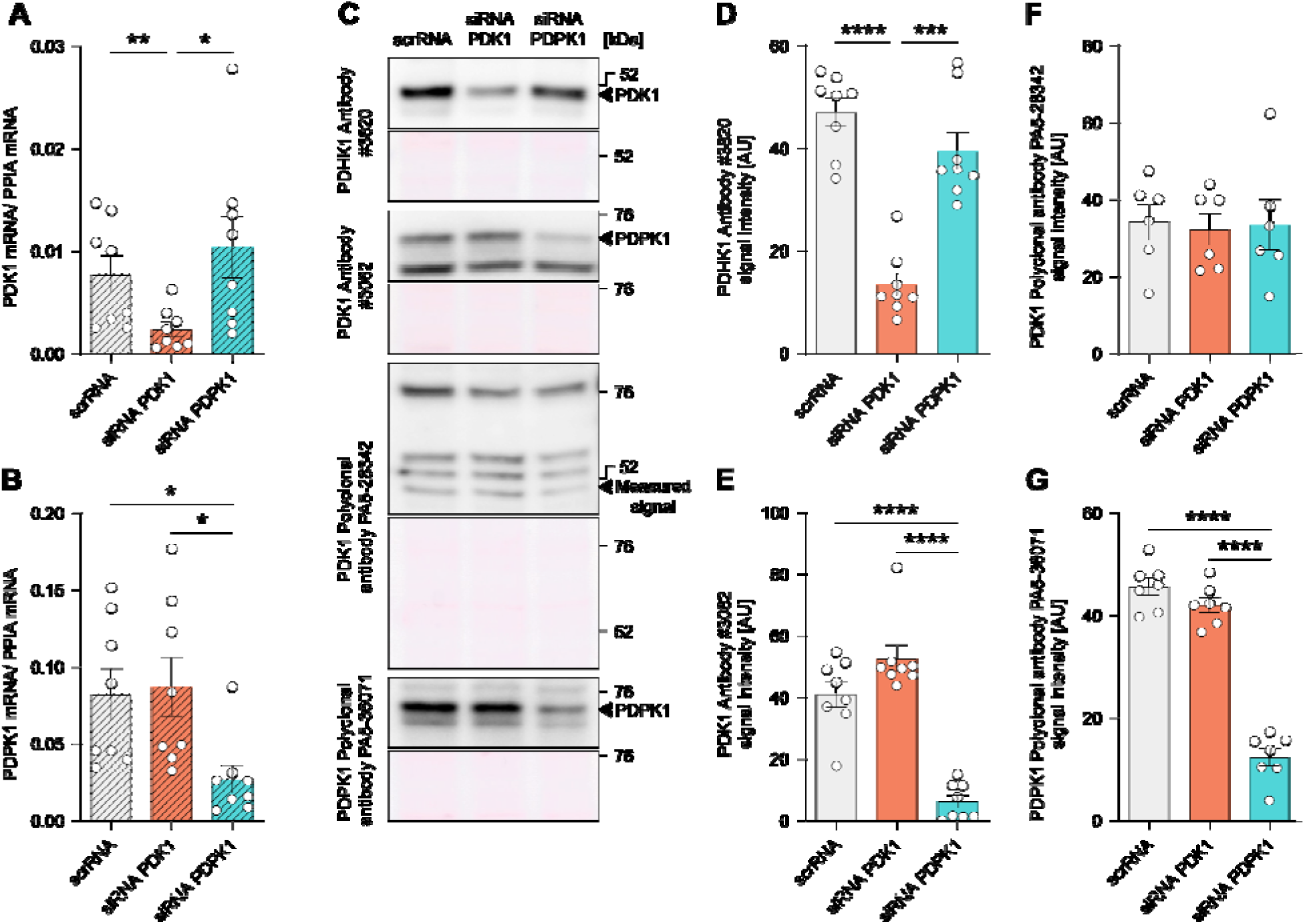
Effect of silencing of pyruvate dehydrogenase kinase 1 or 3-phosphoinositide-dependent protein kinase 1 on their immunodetection by different primary antibodies. (**A**) Pyruvate dehydrogenase kinase 1 (PDK1) mRNA and (**B**) 3-phosphoinositide-dependent protein kinase 1 (PDPK1) mRNA (endogenous control: peptidylprolyl isomerase A (PPIA) mRNA). n = 8. (**C**) Western blot results for different primary antibodies with corresponding Ponceau S staining as a loading control. (**D**) Cell Signaling #3820 antibody. n = 8. (**E**) Cell Signaling #3062 antibody. n = 8. (**F**) Thermo Fisher Scientific PA5-28342 antibody. n = 6. (**G**) Thermo Fishe Scientific PA5-36071 antibody. n = 7. Note that on the y-axis, the names of the antibodies are written as specified by the vendor. On the x-axis, silenced mRNAs are indicated with the following abbreviations: PDK1 = pyruvate dehydrogenase kinase 1, PDPK1 = 3- phosphoinositide-dependent protein kinase 1. Data are presented as means ± standard error. *P < 0.05, **P < 0.01, ***P < 0.001, ****P < 0.0001, with comparisons made between all treatments. AU = arbitrary units. Data are from two independent experiments (n = 4 per experiment), resulting in two blots per antibody. For immunoblotting results (**D**, **E**, **F**, **G**): Brown-Forsythe and Welch ANOVA tests with Dunnett T3 test for multiple comparison; for qPCR results (**A**, **B**): RM one-way ANOVA, with Geisser-Greenhouse correction with Tukey test for multiple comparisons.

PDK1 polyclonal antibody PA5-28342 by vendor Thermo Fisher Scientific, with 3- phosphoinositide-dependent protein kinase 1 (concept B) and pyruvate dehydrogenase kinase 1 (concept A) both stated as designated target (access date: September 29, 2025), did not yield visible bands responsive to silencing of 3-phosphoinositide-dependent protein kinase 1 and pyruvate dehydrogenase kinase 1, meaning that it does not detect either kinase in our cell cultures (**Figure 4C**, **F**). Another tested antibody by the same vendor was PDPK1 polyclonal antibody PA5- 36071, whose designated target is 3-phosphoinositide-dependent protein kinase 1 (concept B). However, its product webpage (access date: September 29, 2025) also listed HGNC gene ID number 8809, which corresponds to pyruvate dehydrogenase kinase 1 (concept A). The antibody produced decreased signal in cells with silenced 3-phosphoinositide-dependent protein kinase 1, but not in cells with silenced pyruvate dehydrogenase kinase 1 (**Figure 4C**, **G**).

## Discussion

This study demonstrates that unclear and non-unique abbreviations such as PDK1 lead to selecting incorrect antibodies for protein detection, using incorrect sequences for mRNA measurements or gene silencing, mistaking one protein for another, or attributing the properties of one protein to another. In fact, 22.4 % of articles containing PDK1 as a keyword in PubMed from 2019 to mid-2026 include at least one of these errors. Each of these errors can lead to incorrect interpretations and conclusions that not only enable the spread of false information but also provide a basis for the generation of mistaken hypotheses for new studies, thus indirectly wasting valuable time and resources. Note that some of the authors of the present article have made similar PDK1-related errors (*25*), for example, by citing a paper in which PDK1 is abbreviated as pyruvate dehydrogenase kinase 1, even though the cited article describes characteristics of 3- phosphoinositide-dependent protein kinase 1, such as AKT activation (*26*).

To reduce the errors described in future studies, we recommend the following mitigation strategies:

- Every research paper should use the nomenclature from recognized databases (**Table 1**), which is already suggested by major publishers (*27, 28*). Furthermore, each gene or protein under investigation should be accompanied by database links to avoid any ambiguity. Some research areas, like enzymology (*29*) or mitochondrial physiology (*30*) have, to combat lack of clarity, introduced more precise classification and standardization.
- Researchers should ensure that all used or studied sequences (nucleotide or amino acid) are provided with either a catalog number or the full sequence, while antibodies should always be referenced by their catalog numbers. To improve transparency, this information should be included in the main article rather than in the supplement whenever possible (**Table 1**).
- Reviewers should verify whether the stated sequences and antibodies are valid and properly referenced. In the fields where the use of vague or potentially ambiguous abbreviations is common, such as PDK1, additional attention should be paid to ensure references correspond correctly to the protein of interest (**Table 1**). Importantly, all this cannot be expected from reviewers if they are assigned too little time to prepare the report, given the time pressure of very short review deadlines set by some journals.
- Biotechnology providers and vendors should provide clear and unambiguous descriptions of their products, again using unique abbreviations, full names, and correct links to databases. Wikipedia provides helpful warnings to avoid confusion between the two proteins. In the article for pyruvate dehydrogenase kinase 1, it states: ”*Not to be confused with the master kinase PDK1, 3-phosphoinositide-dependent protein kinase (PDPK1)*” (*31*). Similar warning is found on the 3- phosphoinositide-dependent protein kinase 1 page.

**Table 1.**
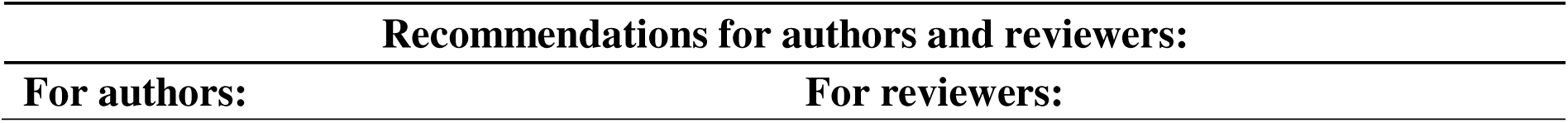
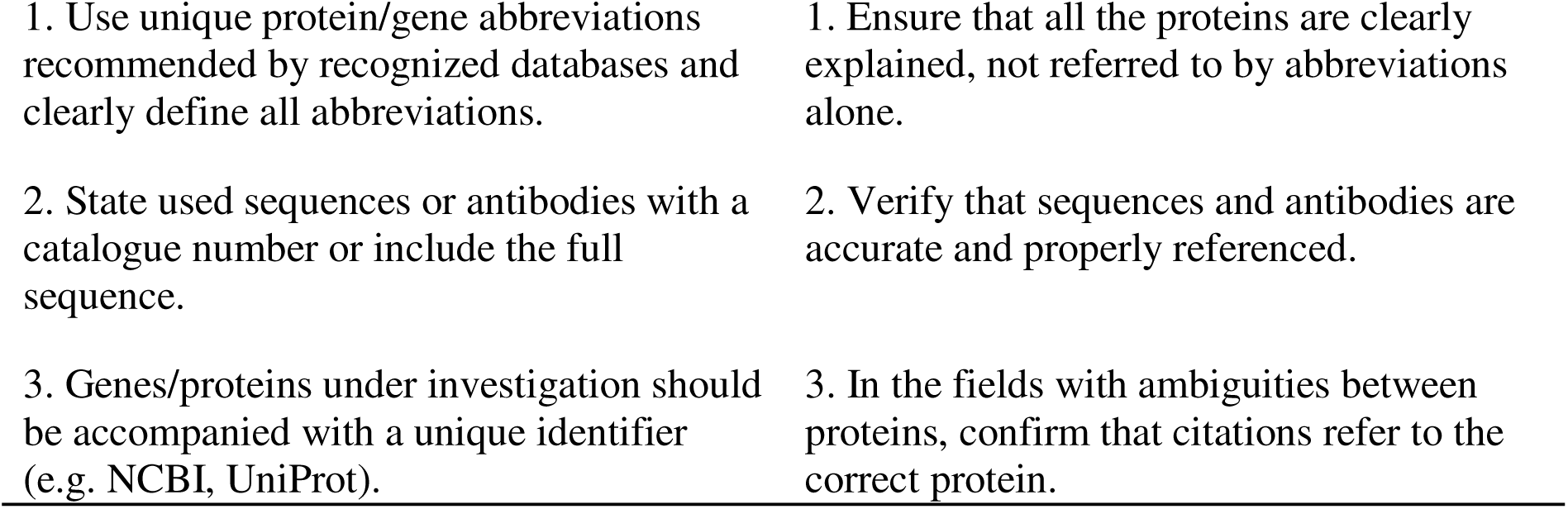
Minimal recommendations for authors and reviewers to reduce ambiguity related to abbreviations. Table outlines three key recommendations for authors and three for reviewers to help prevent some of the errors that use of abbreviation introduces.

|  |  |
| --- | --- |
| 1. Use unique protein/gene abbreviations recommended by recognized databases and clearly define all abbreviations. | 1. Ensure that all the proteins are clearly explained, not referred to by abbreviations alone. |
| 2. State used sequences or antibodies with a catalogue number or include the full sequence. | 2. Verify that sequences and antibodies are accurate and properly referenced. |
| 3. Genes/proteins under investigation should be accompanied with a unique identifier (e.g. NCBI, UniProt). | 3. In the fields with ambiguities between proteins, confirm that citations refer to the correct protein. |

In addition to our recommendations above, the PDK1-TermTracker introduced in this study can be used by the authors and reviewers to curate and verify the correctness of the cited works. With the pressures in modern research, where scientists and reviewers are expected to accomplish more in less time, we contribute a service for PDK1 error-checking, source verification, and support careful manuscript review and prevent errors from slipping through (*32*) and propagate over time.

The PDK1-TermTracker relies on a living pre-annotated database of both correctly and erroneously interpreted PDK1 abbreviations in previously published articles. In addition to this publicly available computational service, all the annotated papers (labeled by error type categories) are made available, including detailed descriptions of identified PDK1 misinterpretations.

There is ample room for further work, as the ambiguity surrounding PDK1 is not an isolated phenomenon. Some articles made similar errors with pyruvate dehydrogenase kinase 2 (HGNC symbol: PDK2, UniProt: Q15119) and a hypothetical kinase proposed to phosphorylate AKT, which has alternatively been referred to as PDK2, by analogy to PDK1 (3-phosphoinositide-dependent protein kinase 1) (*33-35*). Even proteins with vaguely similar names, such as pyruvate kinase, resulted in articles using wrong antibodies (*36, 37*). The confusion of abbreviations extends beyond PDK and arises with various other proteins. JNK commonly denotes JUN N-terminal kinases; however, it is occasionally incorrectly used to refer to Janus kinases, which resulted in articles with the same types of errors as with PDK1 (*38-42*). MHC can refer to either myosin heavy chain or major histocompatibility complex (*43, 44*). Complexity increases further, as MYH7 gene encodes myosin heavy chain 7 (UniProt: P12883), which is typically referred to as myosin heavy chain slow/β (MyHC-slow/β) found in type I muscle fibers. Similarly, MYH1 gene encodes myosin heavy chain 1 (UniProt: P12882), which is commonly referred to as myosin heavy chain IIx (MyHC-IIx), where the “II” denotes association with type II muscle fibers (*45*). Additionally the major histocompatibility complex is divided into three classes: MHC class I, II, and III, adding further overlap in terminology (*44*). Unsurprisingly, some articles confused myosin heavy chain isoforms (*46-48*)^8^ or major histocompatibility complexes and myosin heavy chains (*49, 50*).

## Acknowledgments

Use of AI-assisted technologies

We acknowledge the use of AI-assisted technologies, specifically ChatGPT using the GPT-5.5 and GPT-5.6 models with enterprise data protection, to help in the development of Python scripts for bibliographic data management. All AI-assisted outputs, scripts, and retrieved information were critically reviewed and verified by the authors, who take full responsibility for the final content.

## Funding

Slovenian Research And Innovation Agency program P3-0043

Slovenian Research And Innovation Agency grant J7-3153

Slovenian Research And Innovation Agency grant J7-60125

Slovenian Research And Innovation Agency grant BI-NO/25-27-004

Slovenian Research And Innovation Agency program P2-0103

Slovenian Research And Innovation Agency program P3-0154

Slovenian Research And Innovation Agency program PR-12394

## Author contributions

Conceptualization: BlK, ANK, BoK, SrP

Data curation: BlK, ANK, AK, BoK

Formal analysis: BlK, ANK, KD, SeP, AK, BoK

Investigation: BlK, ANK, KM, AVC, BoK, SrP

Methodology: BlK, ANK, KM, AK, NL, BoK Visualization: BlK, ANK, BoK

Funding acquisition: NL, SrP

Project administration: SrP

Resources: KM, SrP Supervision: SrP

Writing – original draft: BlK

Writing – review & editing: BlK, ANK, KM, PMGR, AVC, ACR, EG, KD, SeP, BC, AK, NL, BoK, SrP

Initials key: BlK, Blaž Kociper; ANK, Alja Nike Kastrin; KM, Katarina Miš; PMGR, Pablo M Garcia-Roves; AVC, Alexander V Chibalin; ACR, Arild C Rustan; EG, Erich Gnaiger; KD, Klemen Dolinar; SeP, Senja Pollak; BC, Bojan Cestnik; AK, Andrej Kastrin; NL, Nada Lavrač; BoK, Boshko Koloski; SrP, Sergej Pirkmajer.

## Data, code, and materials availability

All data are available in the main text or the supplementary materials.

## Materials and Methods

### Methods related to citation network analysis

#### Citation network

Although PubMed was used for the primary search of PDK1 articles, it lacks the comprehensive citation data necessary to construct a citation network. Therefore, the WoS database was utilized, as it provides extensive coverage of biomedical literature and, owing to its precise and complete metadata, represents one of the most frequently utilized sources for advanced network analyses. Pubmed identifiers (PMIDs) of the articles manually curated and labeled by domain experts were used to construct a corresponding PMID-based query, which was then submitted to WoS. The network was constructed from bibliometric metadata retrieved from WoS. PMIDs were used to link each WoS record to the corresponding manually assigned label from the curated PDK1 corpus. The analysis was organized around a directed citation network in which nodes represent scientific articles and directed edges represent citation relations: a directed edge from article *u* to article *v* indicates that article *u* cites article *v*. Each node was assigned a manual annotation and a set of attributes derived from the WoS record: PMID, first author, publication date, article title, digital object identifier (DOI), total (global) citation count as reported by WoS, and internal citation count, defined as the in–degree of the node within the retrieved citation network, i.e. the number of citations received from other articles within the same corpus.

To examine whether the citation network exhibits structure beyond what is captured by publication year and citation count—in particular, whether articles associated with different PDK1 meanings or error categories tend to cluster according to their citation relationships—node positions were additionally derived directly from the topology of the citation network.

The directed citation graph was first converted to an undirected adjacency representation: for every pair of articles where at least one cited the other, the corresponding matrix entry was set to one, and zero otherwise. This symmetrization was adopted because the analytical interest lies in shared citation relationships between articles rather than in citation direction specifically, and a symmetric representation allows similarity between articles to be assessed regardless of which article cited which. Each row of the resulting adjacency matrix was then L2-normalized, so that the subsequent embedding step9 reflects the relative composition of each article’s citation neighborhood—that is, the set of other articles with which it shares citation relationships—rather than being dominated by differences in the absolute number of citation links.

The normalized adjacency matrix was projected into two dimensions using Uniform Mani-fold Approximation and Projection (UMAP), a non-linear dimensionality-reduction technique that seeks to preserve local neighborhood structure when mapping high-dimensional data into a lower-dimensional space. Cosine distance was used as the similarity metric, reflecting the orientation rather than the magnitude of each article’s normalized citation-neighborhood vector. The embedding was computed with 15 nearest neighbors and a minimum distance parameter of 0.1, and a fixed random seed was used throughout to ensure that the resulting layout is reproducible across runs. The resulting two-dimensional coordinates were stored as node attributes and subsequently used for visualization.

#### The PDK1-TermTracker application

The citation network and its associated node attributes were integrated into the PDK1-TermTracker, designed to support manual inspection of the labeled corpus and exploration of citation relationships between articles. The application was developed using Observable Framework, a framework for building reactive, data-driven web applications10.

The application provides two complementary views of the network. In the first, node position is set directly from bibliographic metadata—publication year on the horizontal axis and (log-transformed) global citation count from WoS on the vertical axis—so that the user can immediately relate a node’s position to when it was published and how widely it has been cited, independent of any network or text analysis. In the second view, described below, node position instead reflects the topology of the citation network itself.

In both views, node color encodes the manually assigned PDK1 category, and the interaction model is shared throughout: the user can filter the displayed network by category label, and select individual nodes to inspect their associated metadata, including title, publication year, DOI, PMID, and both global and internal citation counts. Selecting a node also highlights its direct citation neighbors, distinguishing articles that it cites from articles that cite it. In addition to node-level inspection, the application provides aggregate, category-level summaries: for a given selection of category labels, the interface reports the number of articles and internal citation links within the selection, together with the number and categorical composition of external articles citing, or cited by, the selected subset, alongside publication-year and citation-count distributions and rankings of the most internally and globally cited articles within the selection. This allows the user to assess, for instance, whether articles in one error category are predominantly cited by articles of a particular other category, which may be informative when investigating how a terminological error originated or propagated through the literature.

The second view replaces the year/citation-count layout with node positions derived from the topology of the citation network, as described above, allowing the same category-level and node-level inspection tools to be used to examine whether articles sharing a PDK1 meaning or error category also share citation relationships.

The technical implementation of the application comprises: (i) Observable Framework (http://observablehq.com/framework/) for reactive application development; (ii) Sigma.js and Graphol-ogy for rendering and interacting with the citation network; (iii) D3.js for auxiliary statistical visualizations, including category distributions and citation-count summaries; and (iv) a Python preprocessing pipeline using NetworkX for citation network construction, and scikit-learn and umap-learn for computing the topology-based two-dimensional embedding described in the Citation network section. The processed network, together with all node attributes (including PMID, category label, publication year, citation counts, citation edges, and visualization coordinates), is serialized to a single JSON file, which is loaded directly by the application at runtime.

### Methods related to in vitro experiments

#### Materials and reagents

Advanced MEM (#12492013), fetal bovine serum (FBS), Gluta-Max (#35050061), MEM vitamins (#11120052), (#A5256701 and #10500064), Gentamicin (#15710049), Fungizone (250 µg/mL amphotericin B) (#15290018), High-Capacity cDNA Reverse Transcription Kit (#4368814), MicroAmp optical 96-well reaction plates (#N8010560), MicroAmp optical adhesive sheets (#4311971), TaqMan Universal Master Mix (#4440038), TaqMan gene expression assays for human pyruvate dehydrogenase kinase 1 (Hs01561847_m1), 3- phosphoinositide-dependent protein kinase 1 (Hs00928927_m1) and peptidylprolyl isomerase A (PPIA) also known as cyclophilin A (Hs99999904_m1) were from Thermo Fisher Scientific (Waltham, MA, USA). The 4–12% Criterion™ XT Bis-Tris polyacrylamide gels (#3450123), XT MES electrophoresis buffer (#1610789), and goat anti-rabbit IgG-horseradish peroxidase conjugate were from Bio-Rad (#1706515) (Hercules, CA, USA). The Amersham ECL Full-Range Rainbow Molecular Weight Marker was from GE Healthcare Life Sciences Cytiva (#RPN800E) (Marlborough, MA, USA). The polyvinylidene difluoride (PVDF) membrane (#IPVH85R) and Immobilon Crescendo Western HRP Substrate Millipore (#WBLUR0500) were from Merck Millipore (Burlington, MA, USA). The primary antibodies used are listed in Table 1. HP Total RNA Isolation Kit was from Omega Bio-tek (#R6812-00S) (Norcross, GA, USA). All other reagents were from Sigma-Aldrich unless otherwise stated.

#### Primary human skeletal muscle cells

Skeletal muscle tissue for preparation of human skeletal muscle cell cultures was obtained with informed written consent, signed by the participants or, in the case of minors, their legal representatives, and with approval by the Republic of Slovenia National Medical Ethics Committee (*m. semitendinosus*, permit numbers 71/05/12 and 0120-698/2017/4). All experiments were conducted in accordance with the Declaration of Helsinki and Good Laboratory Practice regulations.

Human skeletal muscle cell cultures were derived from satellite cells isolated from the *semitendinosus* muscle samples, which were discarded as surgical waste during routine anterior cruciate ligament reconstruction as described (*51*). The muscle tissue was cleaned of connective and adipose tissue, cut into small pieces and trypsinized at 37 °C for 45 min to release the muscle satellite cells. The isolated cells were grown in 100 mm Petri dishes in growth medium (Advanced MEM supplemented with 10 % (v/v) FBS, 0.3 % (v/v) Fungizone, 0.15 % (v/v) Gentamicin, Gluta- Max, and vitamins) at 37 °C and in humidified air with 5 % (v/v) CO_2_. Before reaching confluence, muscle cells were separated into the CD56^+^ and CD56^-^ fractions using MACS CD56 microbeads (Miltenyi Biotec (Bergisch Gladbach, Germany)) as described (*51*). Both fractions were expanded in growth medium until they became subconfluent. At this stage, cells were trypsinized and frozen in growth medium supplemented with 10 % (v/v) DMSO, then were stored in liquid nitrogen until further use. Only the myogenic (CD56^+^) fraction of cells was used for experiments described in this study.

#### Gene silencing

Prior to the experiment, human skeletal muscle myoblasts were seeded on 6-well Matrigel (#356231, Corning) coated plates. Gene silencing was performed on day 2 after seeding. Cells were switched to differentiation medium without antibiotics and antimycotics, 6 h before transfection. Cells were transfected with 10 nM of siRNAs either against pyruvate dehydrogenase kinase 1 (#L-005019-00- 0005 ON-TARGETplus–SMARTpool, Dharmacon Horizon Discovery, target sequences: GAUCA- GAAACCGACACAAU, GCCAGAAUGUUCAGUACUU, GCAUAAAUCCAAACUGCAA,

CAAAGGAAGUCCAUCUCAU) or 3-phosphoinositide-dependent protein kinase 1 (#L003017- 00-0005 ON-TARGETplus–SMARTpool, Dharmacon Horizon Discovery, target sequences: UAUAUUAUGUGGAUCCUGU, GACCAGAGGCCAAGAAUUU, GCAGCAACAUAGAGCAGUA, GAAGCAGGCUGGCGGAAAC). Transfection with scrambled RNA (ON-TARGETplus Non-targeting Control Pool, Dharmacon Horizon Discovery) was used as a negative control. After 24 h, the medium was re-moved and replaced with fresh growth medium which contained antibiotics and antimycotics. After 48 h, cells were harvested for quantitative real-time PCR and after 72 h for immunoblotting.

Quantitative real-time PCR (qPCR). 48 h after transfection myoblasts were washed with phosphate-buffered saline (PBS: 137 mM NaCl, 2.7 mM KCl, 10 mM Na_2_HPO_4_, 1.8 mM KH_2_PO_4_, pH 7.4) and total RNA was isolated using the E.Z.N.A. HP Total RNA Kit. Reverse transcription was performed using the High-Capacity cDNA Reverse Transcription Kit. qPCR was performed with the 7500 Real-Time PCR System and QuantStudio3 (Thermo Fisher Scientific) using TaqMan gene expression assays from Thermo Fisher Scientific (**Table S1**). Results are expressed as gene expression ratio: (1 + E_reference_)^Ct, reference^/(1 + E_target_)^Ct, target^ , where E is the PCR efficiency and Ct is the threshold cycle. PCR efficiency was estimated using LinRegPCR software (*52*). PPIA was used as a reference gene. For raw data see S6 PCR raw data.

Immunoblotting. 72 h after transfection myoblasts were washed 3 times with cold phosphate-buffered saline (PBS: 137 mM NaCl, 2.7 mM KCl, 10 mM Na_2_HPO_4_, 1.8 mM KH_2_PO_4_, pH 7.4). Cells were lysed in Laemmli buffer (62.5 mM Tris-HCl (pH 6.8), 2 % (w/v) sodium dodecyl sulphate (SDS), 10 % (w/v) glycerol, 5 % (v/v) 2-mercaptoethanol, 0.002 % (w/v) bromophenol blue). After sonication, the lysates were heated at 56 °C for 20 min. Samples were loaded onto a precast 4–12 % Bis-Tris polyacrylamide gel, resolved, and subsequently transferred to a polyvinylidene difluoride (PDVF) membrane using the Criterion™ system cell and blotter (from Bio-Rad). The proteins on the membranes were stained with Ponceau S (0.1 % (w/v) in 5 % (v/v) acetic acid) to evaluate the loading of the samples and the efficiency of the transfer. After blocking with 7.5 % (w/v) low-fat dry milk in Tris-buffered saline with Tween (TBST: 20 mM Tris, 150 mM NaCl, 0.02 % (v/v) Tween 20, pH 7.5) for 1 h at room temperature, the membranes were incubated overnight at 4 °C with primary antibodies (**Table S1**). The membranes were then incubated with the horseradish peroxidase-conjugated secondary antibodies in 5 % (w/v) low-fat dry milk in TBST for 1 h at room temperature. The immunoreactive bands were detected by the enhanced chemiluminescence (ECL) method using Immobilon Crescendo Western HRP substrate from Millipore (Billerica, Massachusetts, USA). The signal was recorded with Fusion FX (Vilber (Paris, France)). All the blots are provided in section S7 Western blot raw data.

**Table S1.** Primary and secondary antibodies used for immunoblotting. Abbreviations: PDK1 = pyruvate dehydrogenase kinase 1, PDPK1 = 3-phosphoinositide-dependent protein kinase 1, CST = Cell Signaling Technology (Danvers, MA, USA), TFS = Thermo Fisher Scientific (Waltham, MA, USA), produced by Invitrogen, O/N = overnight. *As stated in the datasheet.

| Primary antibody |  |  |  |  |  | Secondary antibody |  |  |  |  |
| --- | --- | --- | --- | --- | --- | --- | --- | --- | --- | --- |
| Target | kDa | Supplier | Cat. | Ab | Dilution | Time | Supplier | Cat. No. | Dilution | Time |
| protein* |  |  | No. | Host |  |  |  |  |  |  |
| PDK1 | 47 | CST | 3820 | rabbit | 1:1000 | O/N | Bio-Rad | 1706515 | 1:5000 | 1 h |
| PDPK1 | 58 - 68 | CST | 3062 | rabbit | 1:1000 | O/N | Bio-Rad | 1706515 | 1:20,000 | 1 h |
| PDK1 | 49 - 51 | TFS | PA5-28342 | rabbit | 1:500 | O/N | Bio-Rad | 1706515 | 1:20,000 | 1 h |
| PDPK1 | 63 | TFS | PA5-36071 | rabbit | 1:500 | O/N | Bio-Rad | 1706515 | 1:20,000 | 1 h |

## S2 Article list and search

Search on PubMed with the key phrase “PDK1” yielded 1248 hits, with 1169 hits analyzed. Articles containing errors are listed below, grouped by a type of error.

### Search protocol

1. Find for which protein PDK1 stands

1. If article abbreviates both pyruvate dehydrogenase kinase 1 and 3- phosphoinositide-dependent protein kinase 1 to PDK1 = **M**
2. If article does not explain abbreviation (does not state which protein PDK1 stands for) protein is deduced from the context (e.g. PDK1 phosphorylates AKT is a characteristic of 3-phosphoinositide-dependent protein kinase 1, while PDK1 phosphorylates PDH is a characteristic pyruvate dehydrogenase kinase 1)
2. Check to which protein do citations relate:

1. If there are 2 or more citations related to the other protein (e.g. article abbreviates pyruvate dehydrogenase kinase 1 to PDK1, but there are 2 citations to 3- phosphoinositide-dependent protein kinase 1) = **M**
2. _2.2._ If only 1 citation was incorrect and there were no errors related to antibodies/sequences = **Ã_C_ or** B̃**_C_**
3. Check for antibodies and sequences and others:

_3.1._ Antibodies are checked by looking at the stated target as per biotechnology provider and if they relate to the other protein, inhibitor related to the other protein (excluding ambiguously labelled antibodies in **Table S2**) = **Ã_1_ or** B̃**_1_**
3.2. Sequences are checked by searching them on BLAST alignment tool and if they relate to the other protein, or 3D structure, amino acid sequence related to the other protein = **Ã_2_ or** B̃**_2_**
4. If none of the mistakes are found, article is classified as: **A or B**

### Checklist

1. Check what article abbreviates PDK1 to (it is helpful to use CTRL + F and search “pyruvate” and “inositol”), check both.
2. Check if articles used methods using antibodies (often found in the supplement), check the antibodies under PDK1 and compare to biotechnology provider stated target,
3. Check if article used methods requiring sequences (often found in the supplement), (sometimes found in the figures as a picture of the sequence, requires manual copying of letters), check the sequences under PDK1 on BLAST
4. Check for additional identifiers of a protein, such as PDK1 inhibitors, PDK1 3D structures
5. Check the citations, those relating to PDK1 abbreviations (example: *“PDK1 is a protein **[6, 7, 24]”***, here check all three), also check citations that include PDK1 in the title, or mention pyruvate dehydrogenase kinase or 3-phosphoinositide-dependent protein kinase 1 (CTRL + F is useful here again, you can additionally search for “HIF” and “AKT” as they often correlate with either of the proteins)

### Additional explanations

**A** is an article correctly related to pyruvate dehydrogenase kinase 1

**B** is an article correctly related to 3-phosphoinositide-dependent protein kinase 1

**Ã_1_** is an article related to pyruvate dehydrogenase kinase 1 but uses 3-phosphoinositide-dependent protein kinase 1 antibody, or inhibitor, or activator…

B̃**_1_** is an article related to 3-phosphoinositide-dependent protein kinase 1 but uses pyruvate

dehydrogenase kinase 1 antibody, or inhibitor, or activator…

**Ã_2_** is an article related to pyruvate dehydrogenase kinase 1, but uses 3-phosphoinositide-dependent protein kinase 1 nucleic sequence, amino acid sequence, or 3D structure, …

B̃**_2_** is an article related to 3-phosphoinositide-dependent protein kinase 1, but uses pyruvate

dehydrogenase kinase 1 nucleic sequence, amino acid sequence, or 3D structure, …

**Ã_C_** is an article related to pyruvate dehydrogenase kinase 1, but there is **ONE** citation related to 3-phosphoinositide-dependent protein kinase 1

B̃**_C_** is an article related to 3-phosphoinositide-dependent protein kinase 1, but there is **ONE** citation related to pyruvate dehydrogenase kinase 1

U Unrelated topic

E Erratum R Retraction

K Autosomal dominant polycystic kidney diseases related article

PREPRINT Preprint

Note that second experts article sample did not include excluded hits (that is U, E R, K and PREPRINT categories. Furthermore, second expert had to categorize into 5 groups; A, B, Ã (which includes both Ã_1_ and Ã_2_), B̃ (which includes both B̃_1_ and B̃_2_), Ã_C_ and B̃_C_.

Comparison between the experts can be found in S5 Expert comparison.

### Priority ladder

**M > Ã or** B̃, **Ã or** B̃ **> Ã_C_ or** B̃**_C_**

If article satisfies conditions for **M**, it is determined to be **M,** no matter the other mistakes

If article satisfies conditions for **Ã or** B̃ (and not conditions for M), it is determined to be **Ã or** B, no matter the other mistakes

**Table S2.** Ambiguously antibodies labeled by biotechnology vendors. If any of these antibodies were claimed to be used for detection of either of the two proteins, they were not considered a mistake.

| Biotechnology provider | Antibody | Other names |
| --- | --- | --- |
| Abcam | 4A11 | Ab110025 |
| Abcam | ab90444 |  |
| Abcam | EPR19571 | ab202468 |
| Abcam | SP270 | ab227682 |
| Rockland Immunochemicals | 600-401-411 |  |
| Thermo Fisher Scientific | PA5-18441 |  |
| Thermo Fisher Scientific | A302-130A-T |  |
| Thermo Fisher Scientific | PA5-82971 |  |
| Thermo Fisher Scientific | PA5-79797 |  |
| Thermo Fisher Scientific | PA5-79798 |  |
| Thermo Fisher Scientific | PA1-16945 |  |
| Thermo Fisher Scientific | PA5-115732 |  |
| Thermo Fisher Scientific | PA5-106712 |  |
| Thermo Fisher Scientific | PA5-28342 |  |
| Thermo Fisher Scientific | 3B7 | H00005163-M01 |
| Thermo Fisher Scientific | 4A11 | MA5-15797 |
| Thermo Fisher Scientific | 3E1 | H00005163-M02 |
| Thermo Fisher Scientific | 9A7X4 | MA5-55304 |
| Thermo Fisher Scientific | 5T6S8 | MA5-35736 |
| Thermo Fisher Scientific | 8J7H4 | MA5-55303 |
| Thermo Fisher Scientific | JA67-30 | MA5-32702 |
| Thermo Fisher Scientific | 18262-1-AP |  |
| Thermo Fisher Scientific | PA5-120471 |  |
| Thermo Fisher Scientific | PA5-36071 |  |
| Novus Biologicals | BSA Free NBP2-44132 |  |
| Proteintech | 10026-1-AP |  |

## S4 Ambiguities by biotechnology providers and vendors

### Abcam

#### Anti-PDK1 antibody [4A11] (ab110025)

Date of website access: 31.12.2024

Website link: https://www.abcam.com/en-us/products/primary-antibodies/pdk1-antibody-4a11-ab110025

Antibody website provided target information as that of pyruvate dehydrogenase kinase 1 (Figure S1), yet in the datasheet information was about 3-phosphoinositide-dependent protein kinase 1 (Figure S2). Note that information in the datasheet has since been changed referring to pyruvate dehydrogenase kinase 1.

**Figure S1.**
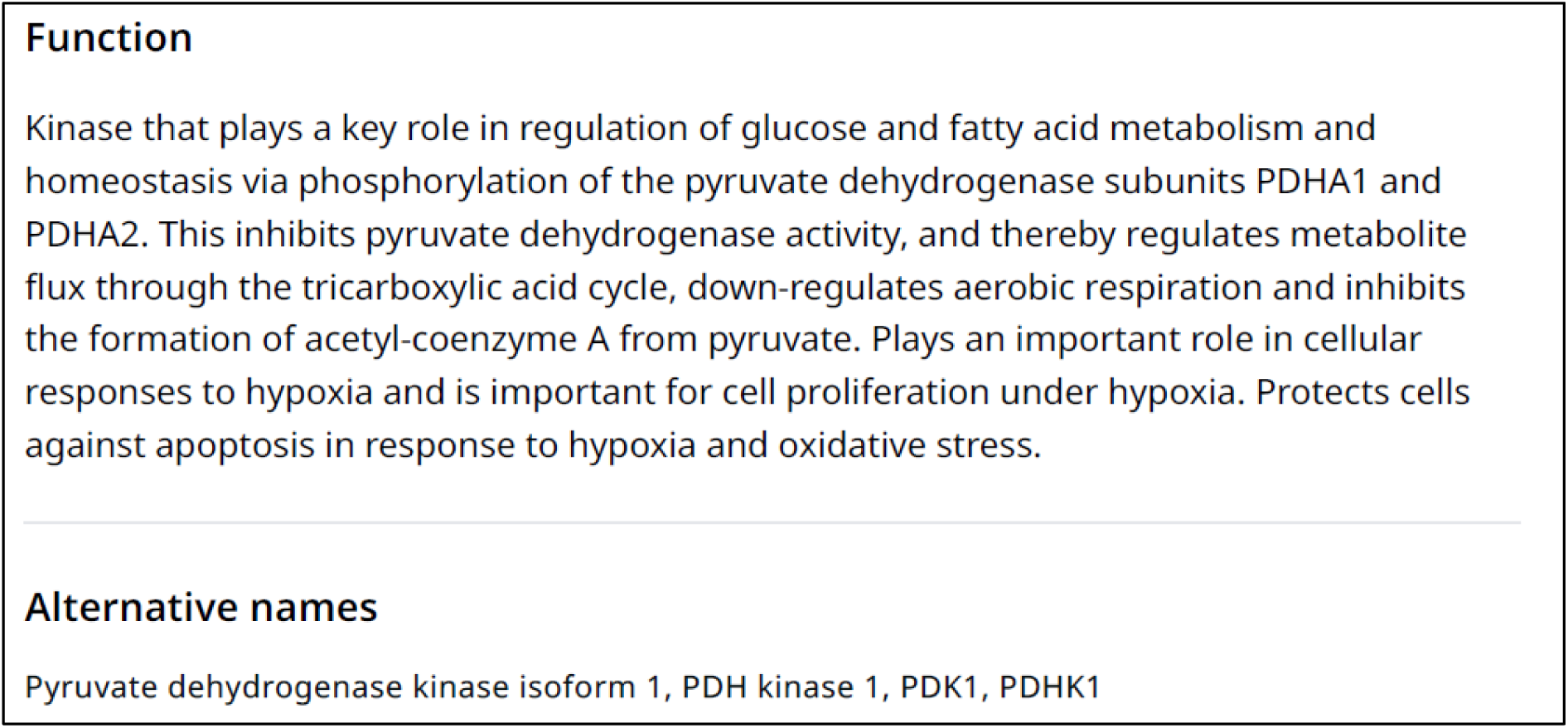
Screenshot of the webpage. Information on antibody website regarding pyruvate dehydrogenase kinase 1.

**Figure S2.**
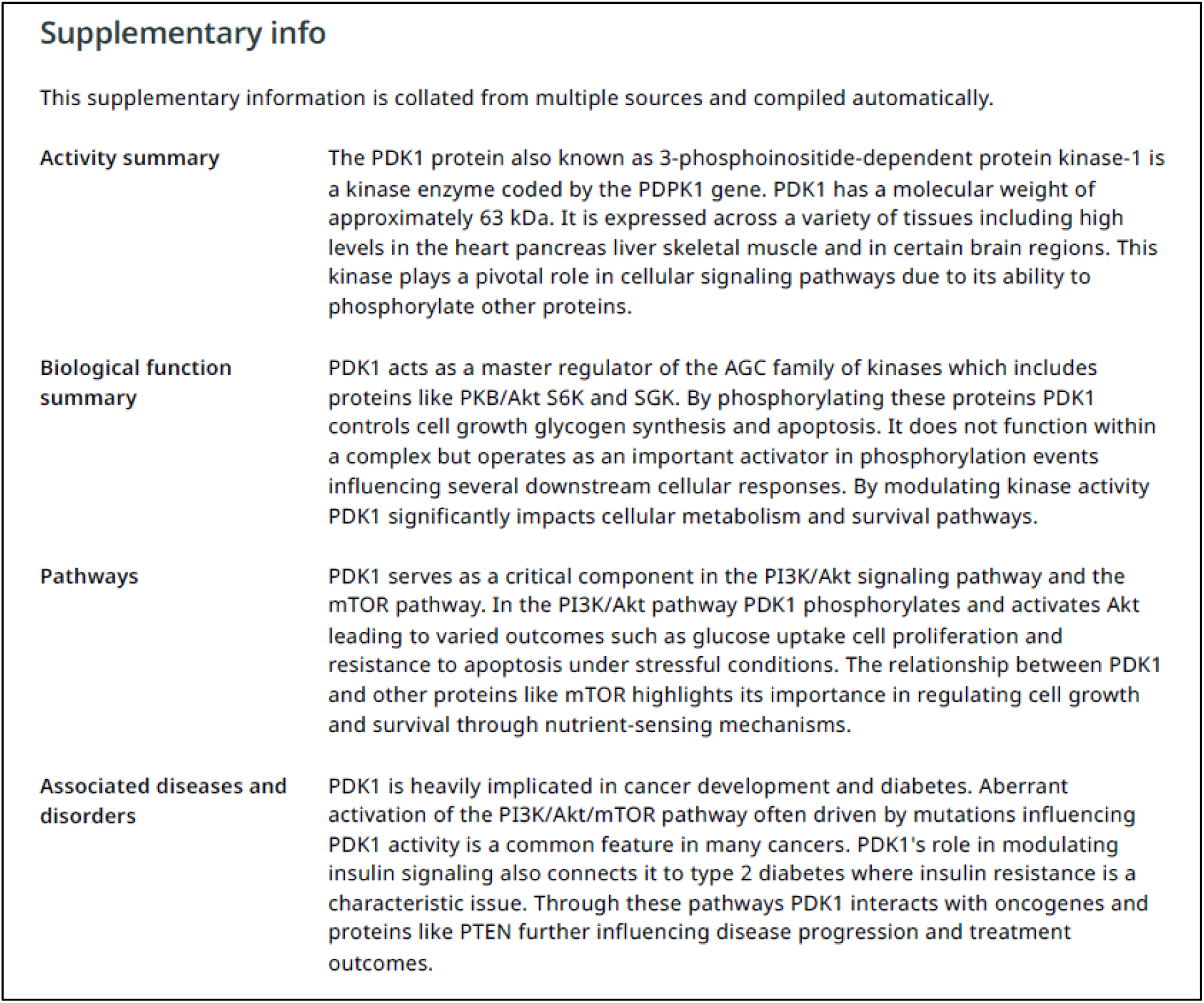
Screenshot of datasheet. Information in antibody datasheet regarding 3-phosphoinositide dependent kinase 1.

#### Anti-PDK1 antibody (ab90444)

Date of website access: 3.1.2025

Website link: https://www.abcam.com/en-us/products/primary-antibodies/pdk1-antibody-ab90444#

Antibody website provided target information as that of pyruvate dehydrogenase kinase 1 (**Figure S3**), yet in the Supplementary info part of the webpage information was about 3-phosphoinositide-dependent protein kinase 1 (**Figure S4**). Note that Supplementary info has since been removed from the webpage.

**Figure S3.**
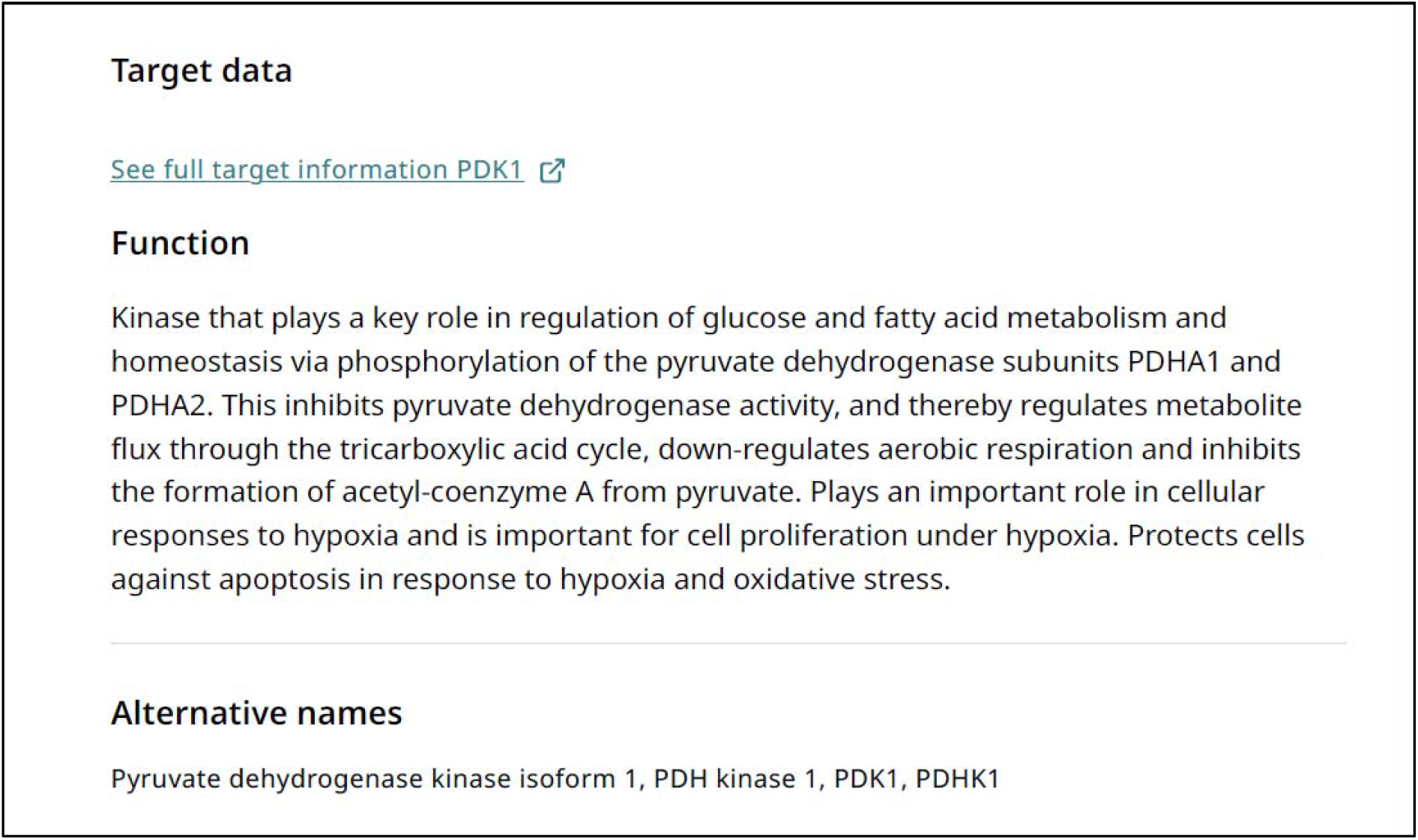
Screenshot of the webpage. Information on antibody website regarding pyruvate dehydrogenase kinase 1.

**Figure S4.**
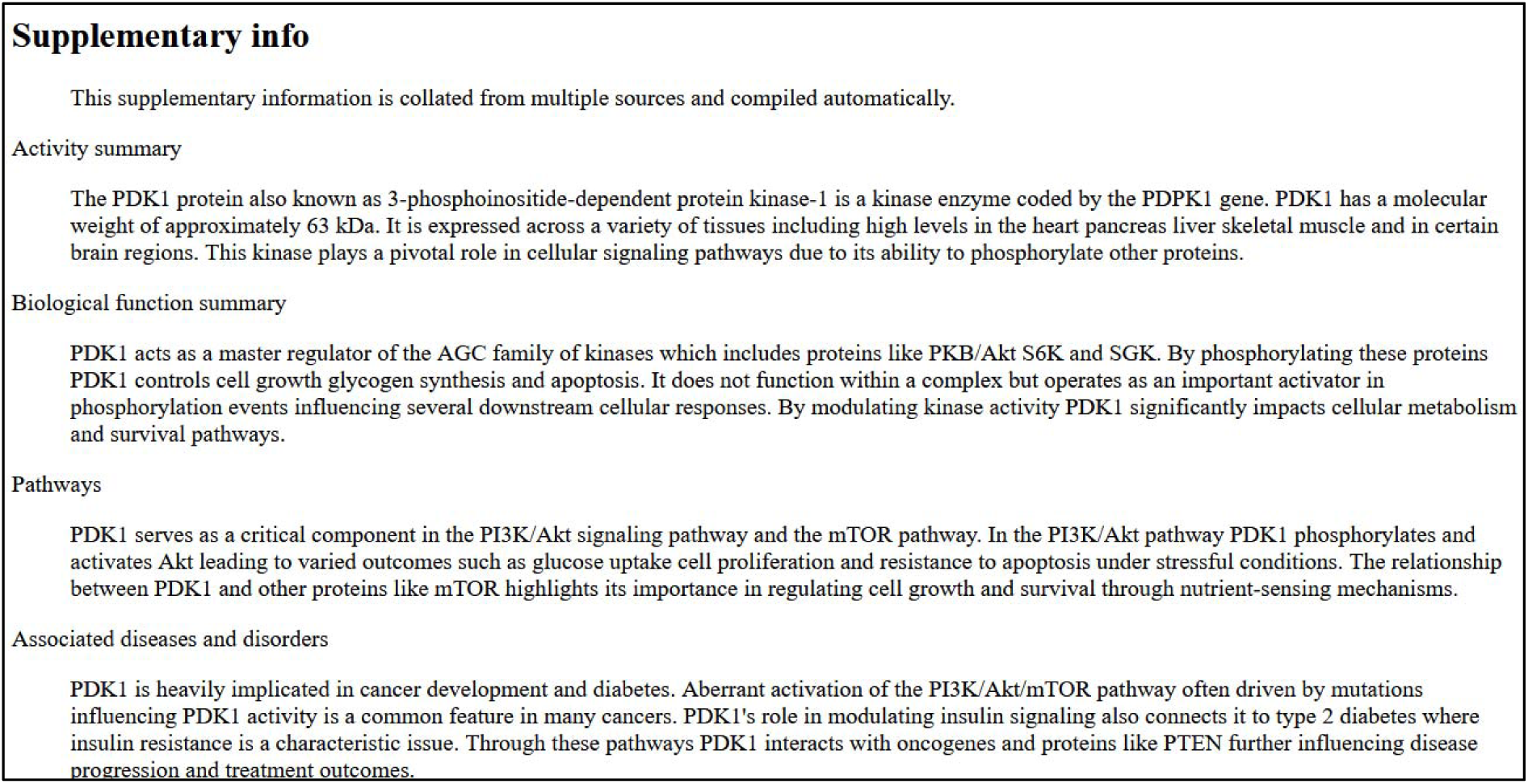
Screenshot of saved webpage. Information under Supplementary info regarding 3-phosphoinositide dependent kinase 1. Webpage style looks different, due to screenshot being taken on a saved website from 3.1.2025.

#### Anti-PDK1 antibody [EPR19571] [ab202468]

Date of website access: 3.1.2025

Website link: https://www.abcam.com/en-us/products/primary-antibodies/pdk1-antibody-epr19571-ab202468

Antibody website provided target information as that of pyruvate dehydrogenase kinase 1 (**Figure S5**), yet in the datasheet information was about 3-phosphoinositide-dependent protein kinase 1 (**Figure S6**). Note that information in the datasheet has since been changed referring to pyruvate dehydrogenase kinase 1.

**Figure S5.**
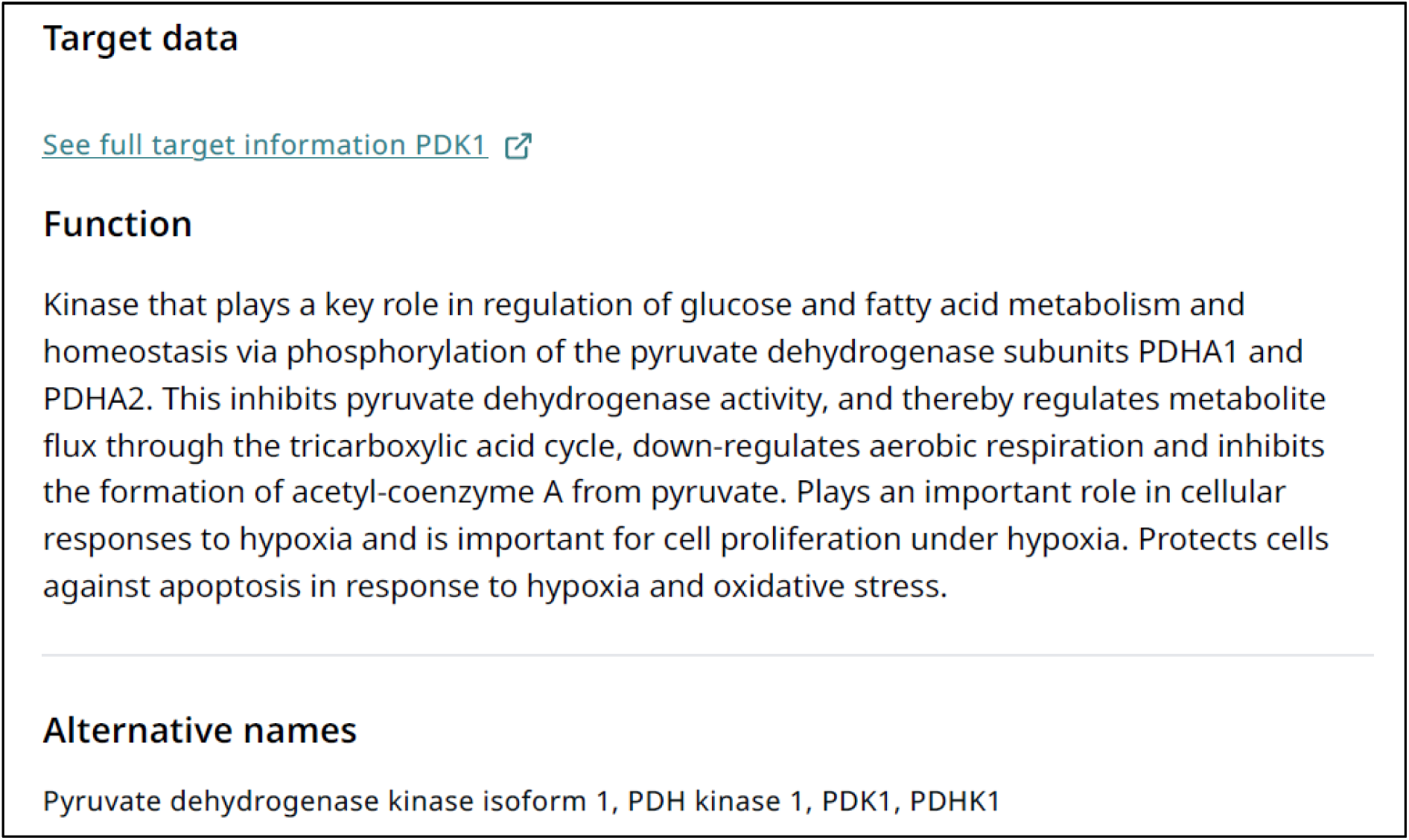
Screenshot of the webpage. Information on antibody website regarding pyruvate dehydrogenase kinase 1.

**Figure S6.**
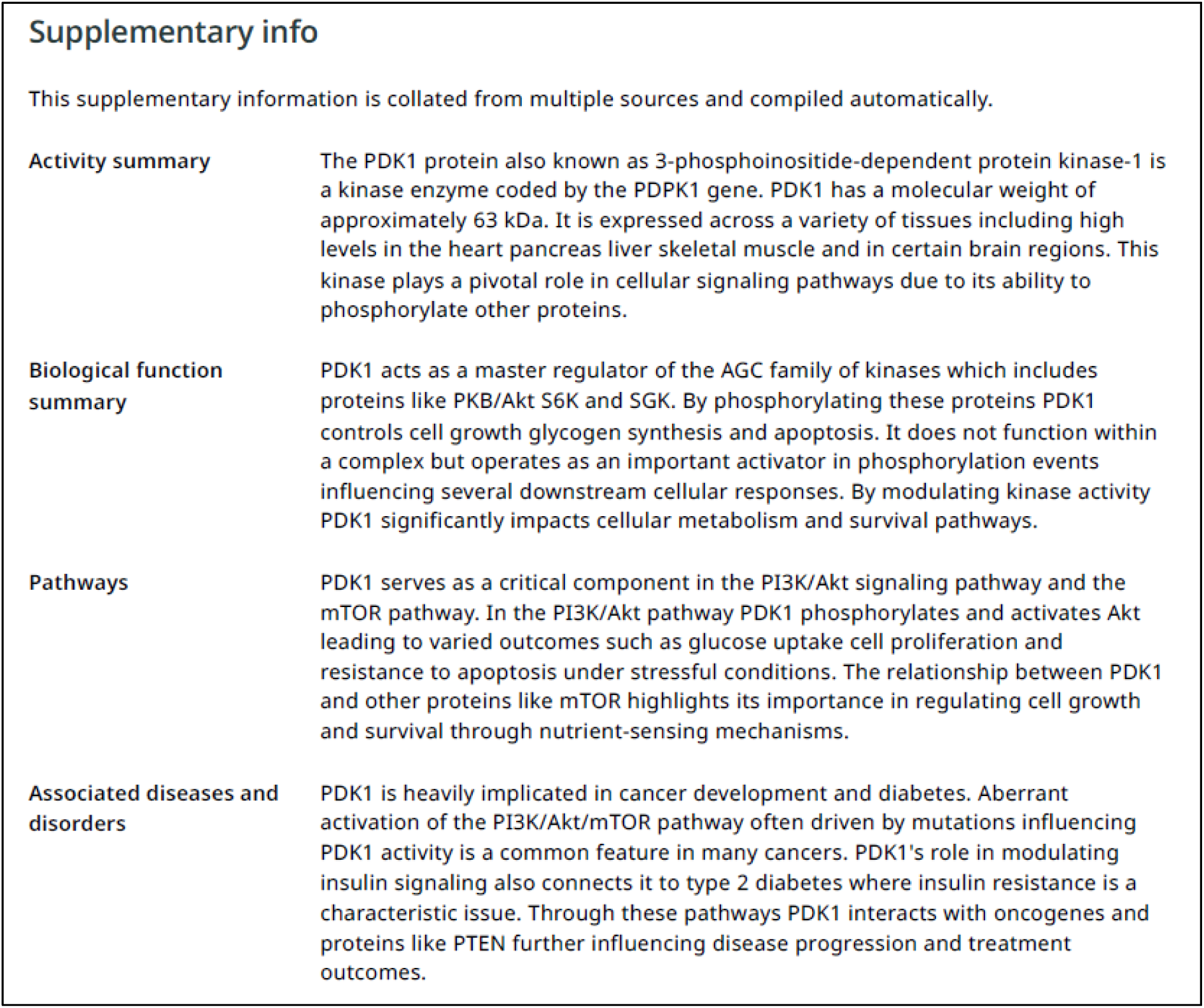
Screenshot of datasheet. Information in antibody datasheet regarding 3-phosphoinositide dependent kinase 1.

#### Anti-PDK1 antibody [SP270] - C-terminal (ab227682)

Date of website access: 3.1.2025

Website link: https://www.abcam.com/en-us/products/primary-antibodies/pdk1-antibody-sp270-c-terminal-ab227682

Antibody website provided target information as that of pyruvate dehydrogenase kinase 1 (**Figure S7**), yet in the datasheet information was about 3-phosphoinositide-dependent protein kinase 1 (**Figure S8**). Note that information in the datasheet has since been changed referring to pyruvate dehydrogenase kinase 1.

**Figure S7.**
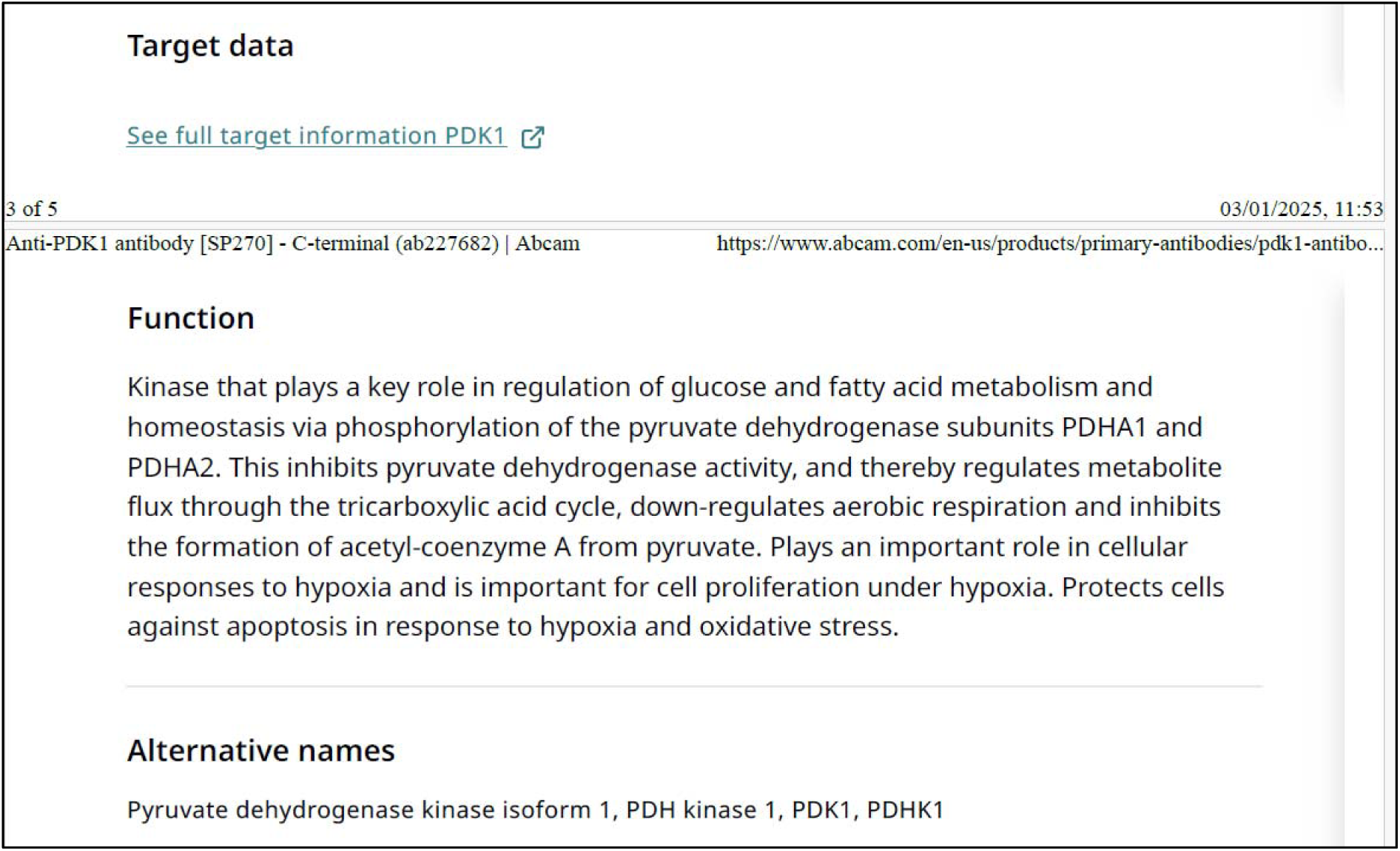
Screenshot of the webpage. Information on antibody website regarding pyruvate dehydrogenase kinase 1.

**Figure S8.**
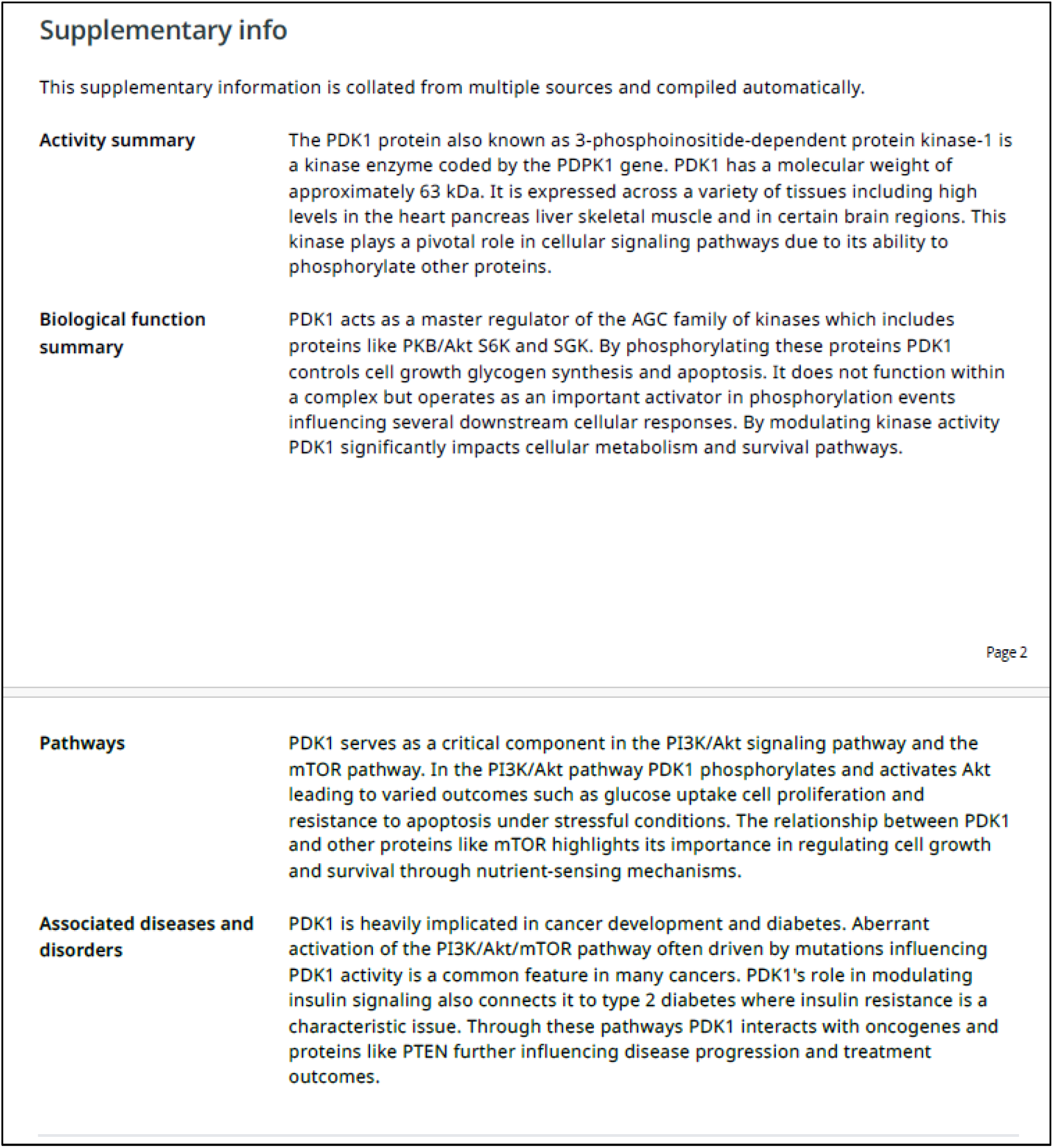
Screenshot of datasheet. Information in antibody datasheet regarding 3-phosphoinositide dependent kinase 1.

### Rockland Immunochemicals

#### PDK1 Polyclonal Antibody [#600-401-411]

Date of website access: 21.7.2025

Website link: https://www.rockland.com/categories/primary-antibodies/pdk1-antibody-600-401-411/

Antibody website provides target information as that of 3-phosphoinositide-dependent protein kinase 1 (**Figure S9**), yet one of the database links (Q15118 UniProtKB) links to pyruvate dehydrogenase kinase 1 (**Figure S10**), with other two linking to 3-phosphoinositide-dependent protein kinase 1.

**Figure S9.**
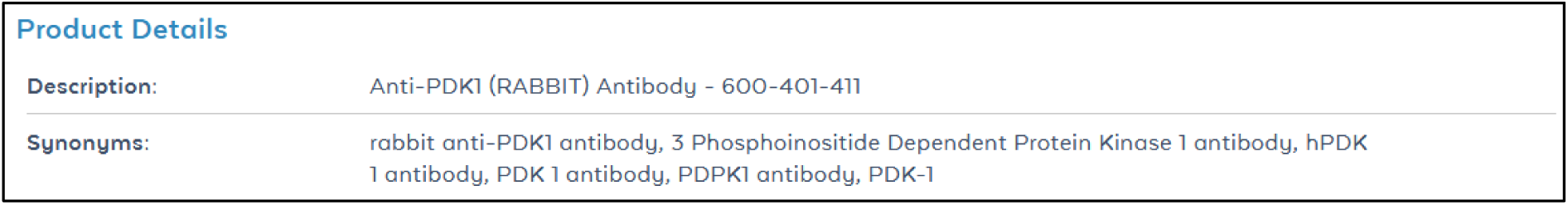
Screenshot of the webpage. Information on antibody website regarding 3-phosphoinositide-dependent protein kinase 1.

**Figure S10.**
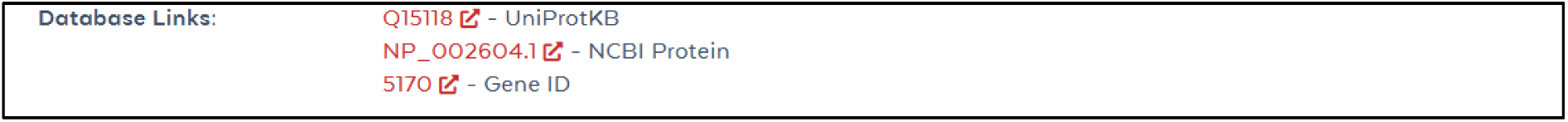
Screenshot of the webpage. First of the database links – Q15118 for UniProtKB – links to pyruvate dehydrogenase kinase 1.

### Sigma Aldrich

#### Anti-PDK1 antibody produced in rabbit

Date of website access: 21.7.2025

Website link: https://www.sigmaaldrich.com/SI/en/substance/antipdk1antibodyproducedinrabbit1234598765

Antibody website provides target information where you can find both 3-phosphoinositide-dependent protein kinase 1 and pyruvate dehydrogenase kinase 1 under synonyms (**Figure S11**).

**Figure S11.**
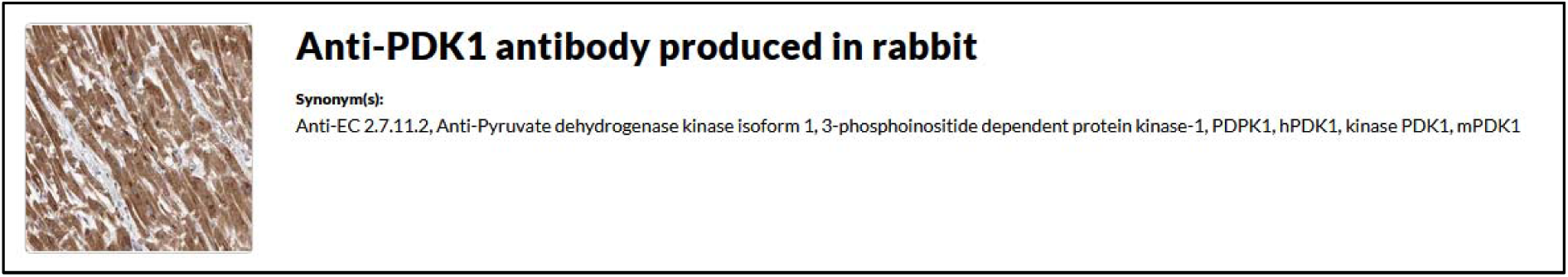
Screenshot of the webpage. Both pyruvate dehydrogenase kinase 1 and 3-phosphoinositide-dependent protein kinase 1 are listed as synonyms.

### Thermo Fisher Scientific

#### PDK1 Polyclonal Antibody [#PA5-18441]

Date of website access: 21.7.2025

Website link: https://www.thermofisher.com/antibody/product/PDK1-Antibody-Polyclonal/PA5-18441

Antibody website provides target information where both 3-phosphoinositide-dependent protein kinase 1 an pyruvate dehydrogenase kinase 1 are referenced as targets (**Figure S12 and S13**). Provided immunogen sequence DFKDKSAEDAK corresponds to pyruvate dehydrogenase kinase 1 (**Figure S14**).

**Figure S12.**
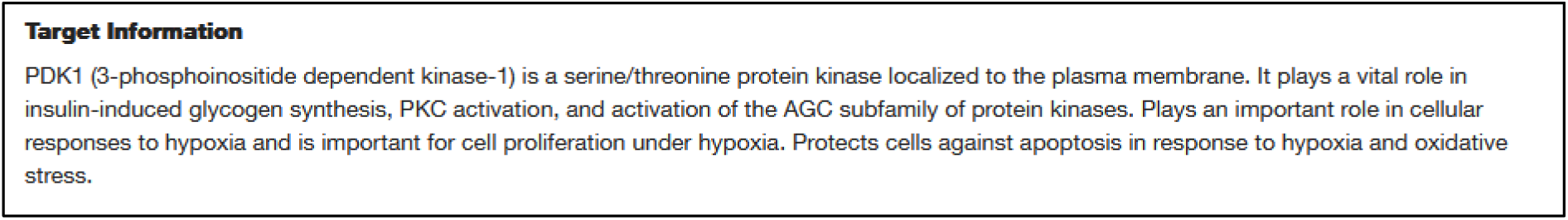
Screenshot of the webpage. Information on antibody website regarding 3-phosphoinositide-dependent protein kinase 1.

**Figure S13.**
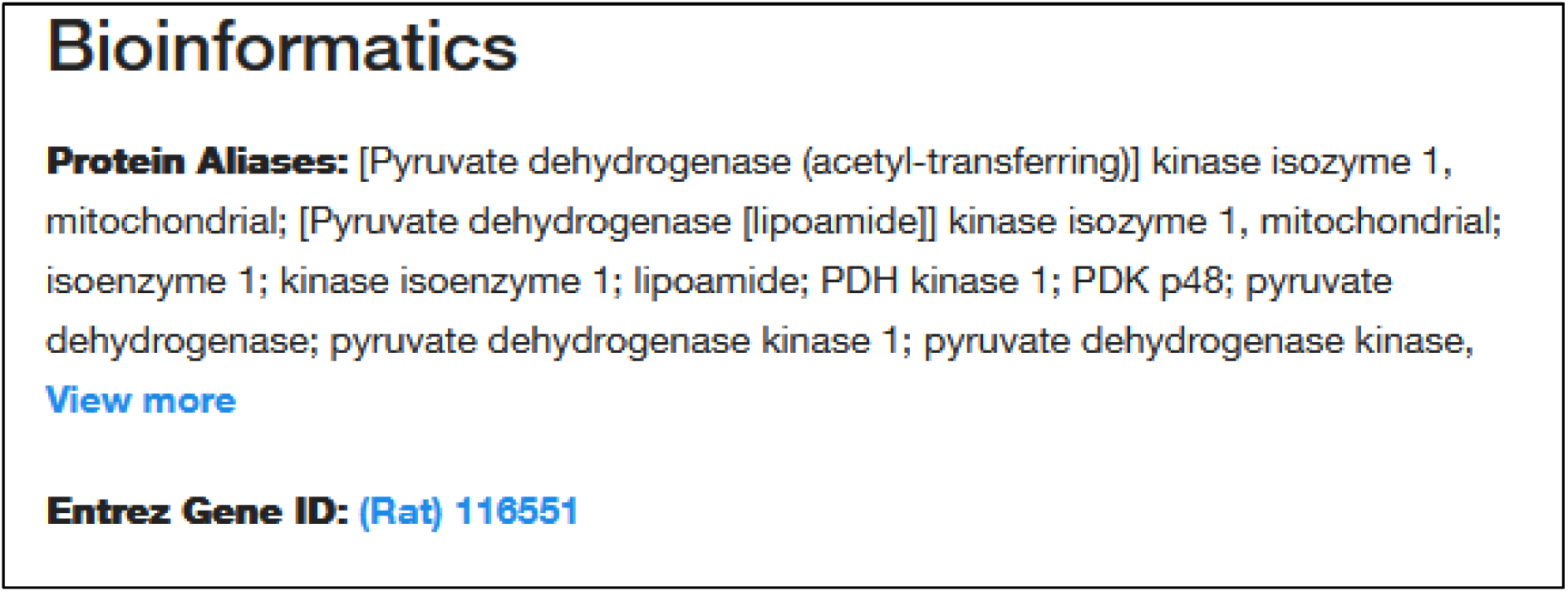
Screenshot of the webpage. Information on antibody website regarding pyruvate dehydrogenase kinase 1.

**Figure S14.**
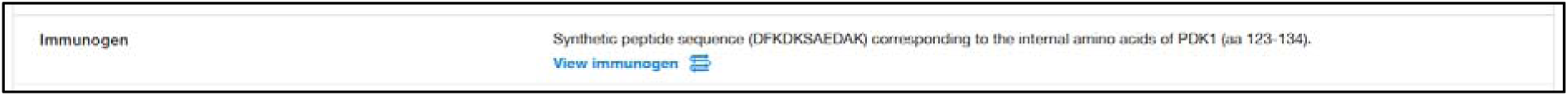
Screenshot of the webpage. Stated immunogen sequence.

#### PDK1 Polyclonal Antibody [#A302-130A-T]

Date of website access: 22.10.2024

Website link: https://www.thermofisher.com/antibody/product/PDK1-Antibody-Polyclonal/A302-130A-T Antibody website provided target information where both 3-phosphoinositide-dependent protein kinase 1 an pyruvate dehydrogenase kinase 1 were referenced as targets (**Figure S15 and S16**). Information on the webpage has since been corrected.

**Figure S15.**
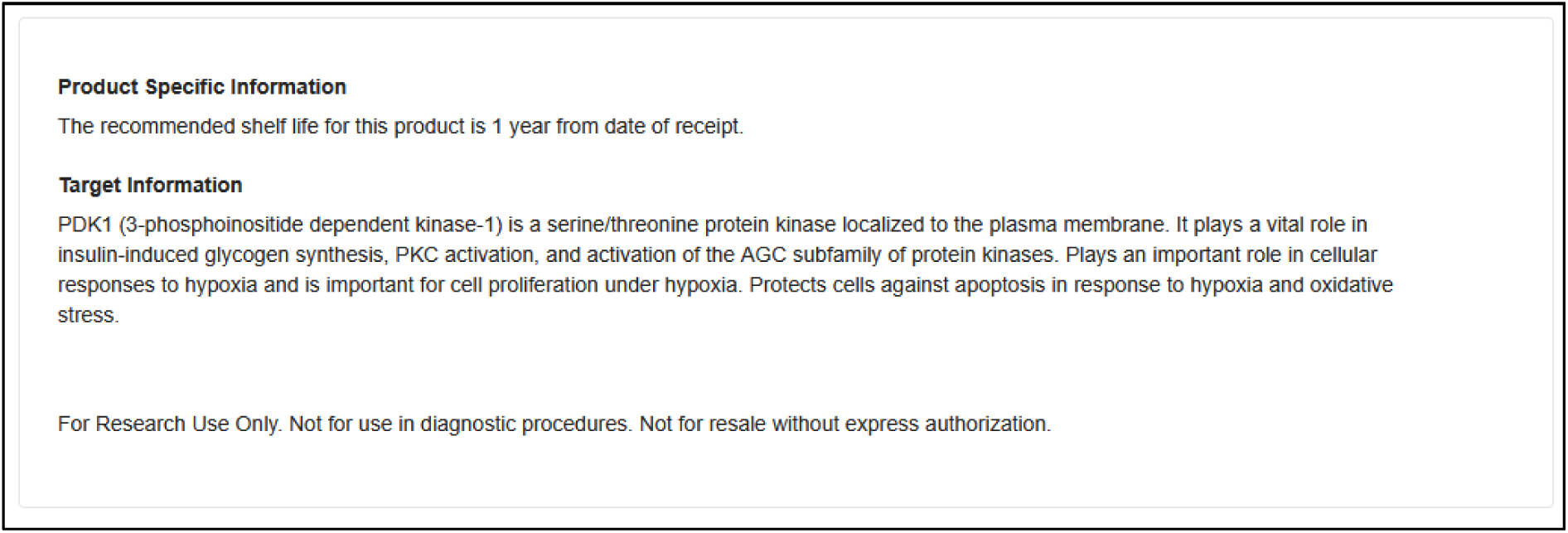
Screenshot of the webpage. Information on antibody website regarding 3-phosphoinositide-dependent protein kinase 1.

**Figure S16.**
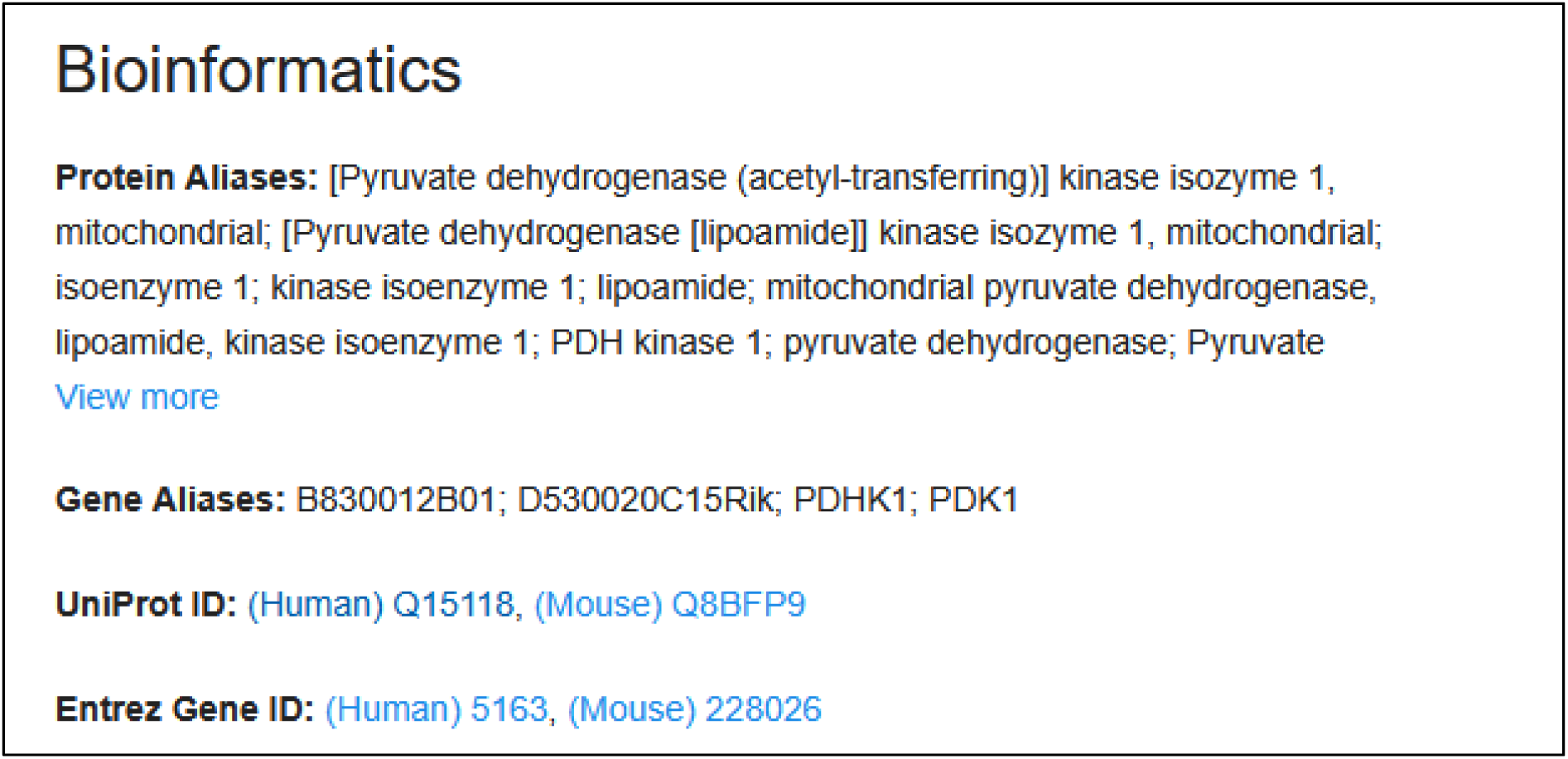
Screenshot of the webpage. Information on antibody website regarding pyruvate dehydrogenase kinase 1.

#### PDK1 Polyclonal Antibody [#PA5-82971]

Date of website access: 22.7.2025

Website link: https://www.thermofisher.com/antibody/product/PDK1-Antibody-Polyclonal/PA5-82971

Antibody website provides target information where 3-phosphoinositide-dependent protein kinase 1 is mentioned, but the immunogen sequence corresponds to pyruvate dehydrogenase kinase 1 (**Figure S17**):

“KEISLLPDNLLRTPSVQLVQSWYIQSLQELLDFKDKSAEDAKAIYDFTDTVIRIRNRHND”.

**Figure S17.**
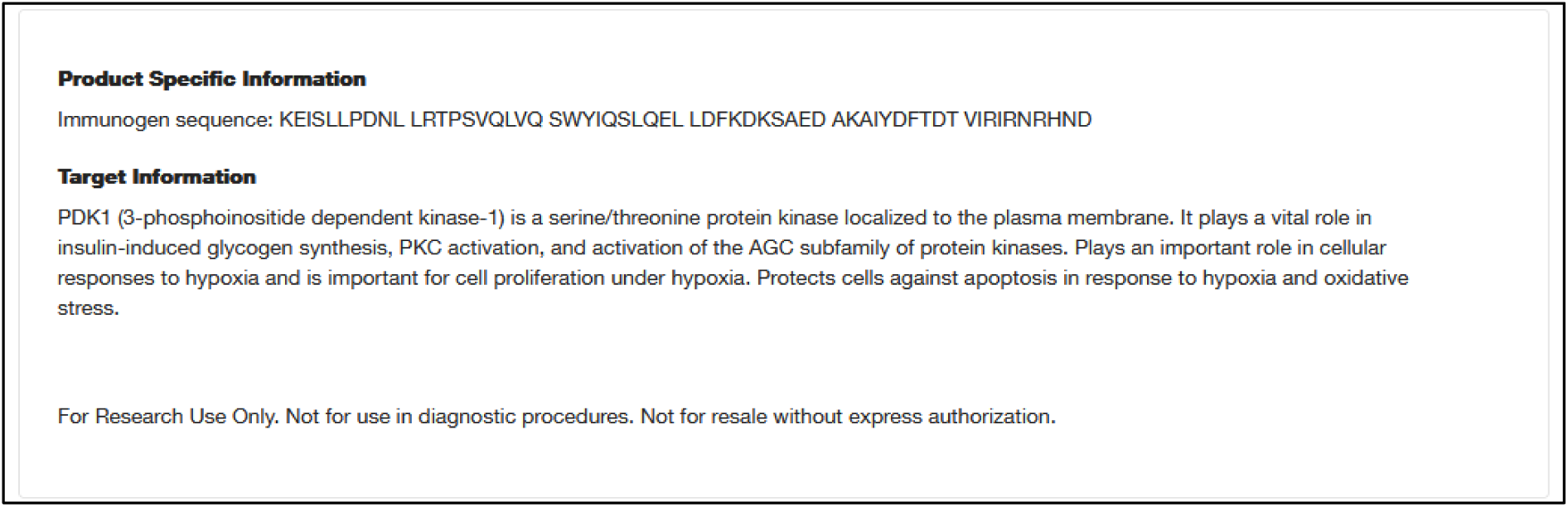
Screenshot of the webpage. Information on antibody website regarding 3-phosphoinositide-dependent protein kinase 1 and immunogen sequence.

#### PDK1 Polyclonal Antibody [#PA5-79797]

Date of website access: 22.7.2025

Website link: https://www.thermofisher.com/antibody/product/PDK1-Antibody-Polyclonal/PA5-79797

Antibody website provides target information where both 3-phosphoinositide-dependent protein kinase 1 an pyruvate dehydrogenase kinase 1 are referenced as targets (**Figure S18 and S19**).

**Figure S18.**
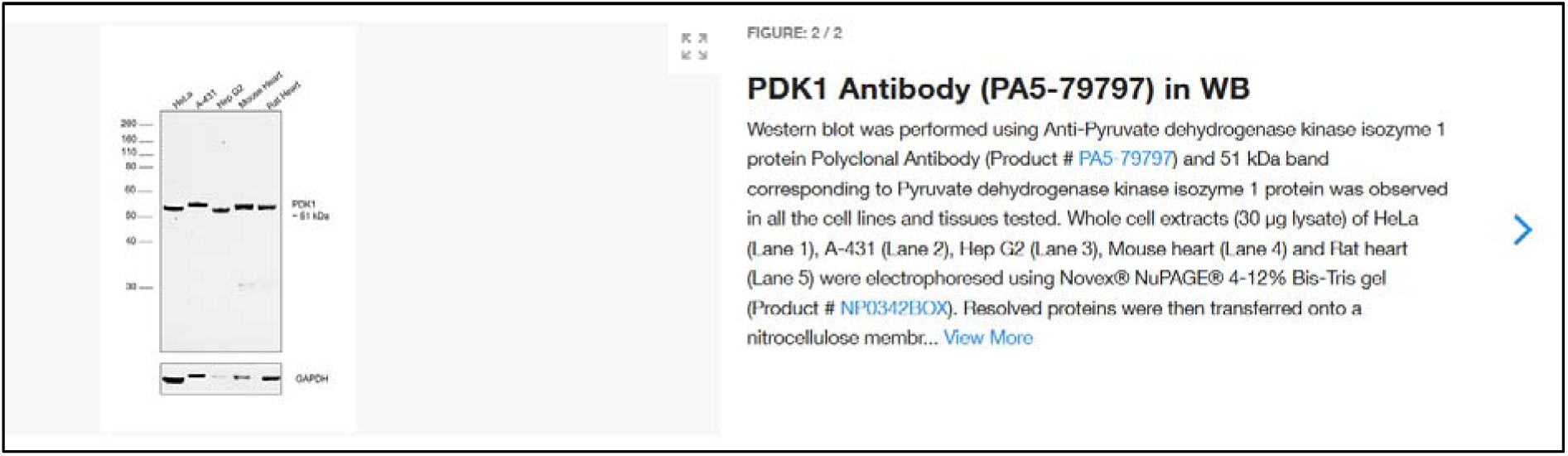
Screenshot of the webpage. Information on antibody website regarding pyruvate dehydrogenase kinase 1.

**Figure S19.**
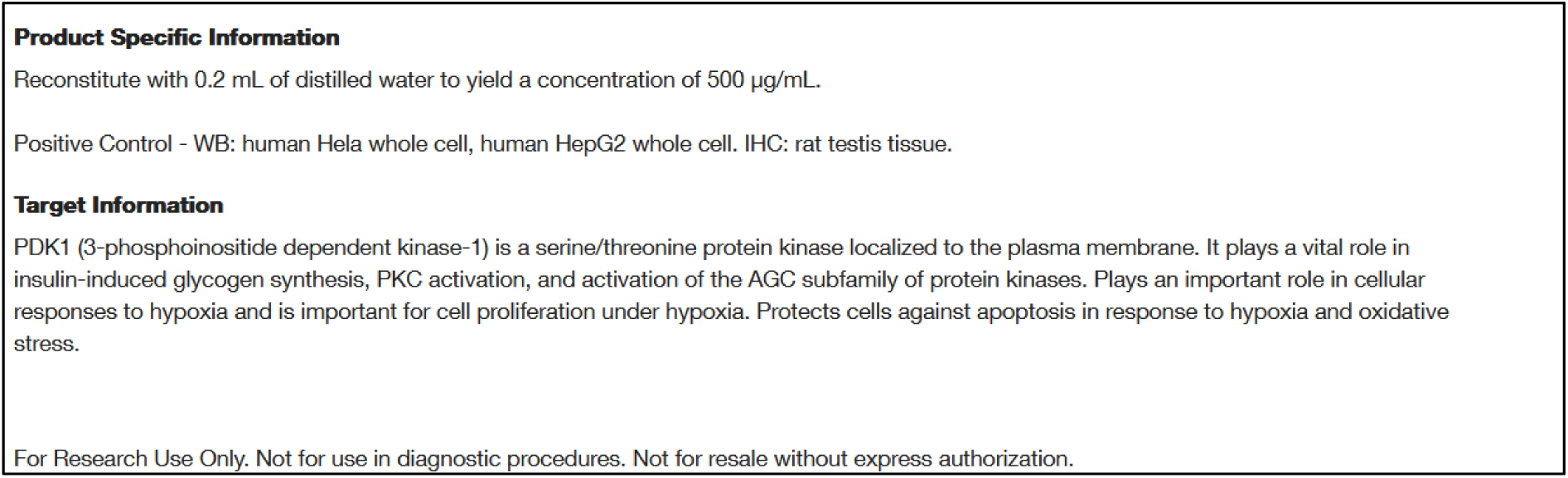
Screenshot of the webpage. Information on antibody website regarding 3-phosphoinositide-dependent protein kinase 1.

#### PDK1 Polyclonal Antibody [#PA5-79798]

Date of website access: 22.7.2025

Website link: https://www.thermofisher.com/antibody/product/PDK1-Antibody-Polyclonal/PA5-79798 Antibody website provides target information where both pyruvate dehydrogenase kinase 1, corresponding immunogen sequence (WKHYNTNHEADDWC) and 3-phosphoinositide-dependent protein kinase 1 are referenced as targets (**Figure S20 and S21**).

**Figure S20.**
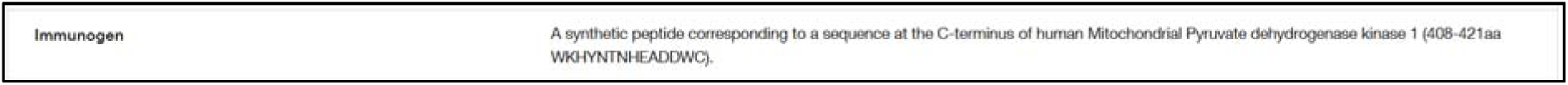
Screenshot of the webpage. Information on antibody website regarding pyruvate dehydrogenase kinase 1 and immunogen sequence.

**Figure S21.**
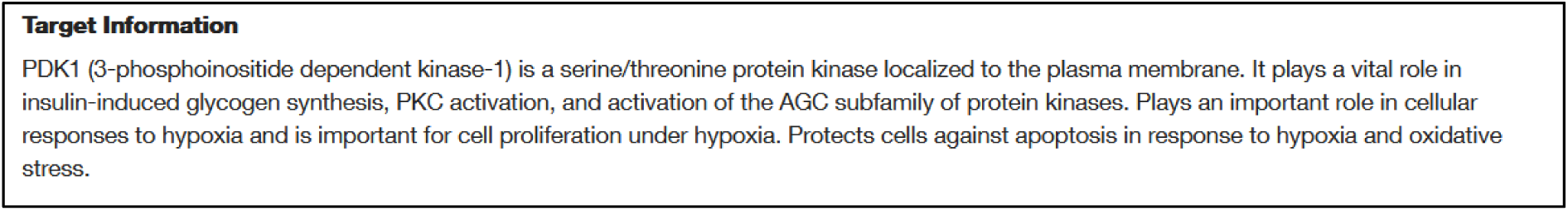
Screenshot of the webpage. Information on antibody website regarding 3-phosphoinositide-dependent protein kinase 1.

#### PDK1 Polyclonal Antibody [#PA1-16945]

Date of website access: 22.7.2025

Website link: https://www.thermofisher.com/antibody/product/PDK1-Antibody-Polyclonal/PA1-16945

Antibody website provides target information where both pyruvate dehydrogenase kinase 1 and 3-phosphoinositide-dependent protein kinase 1 are referenced as targets (**Figure S22 and S23**).

**Figure S22.**
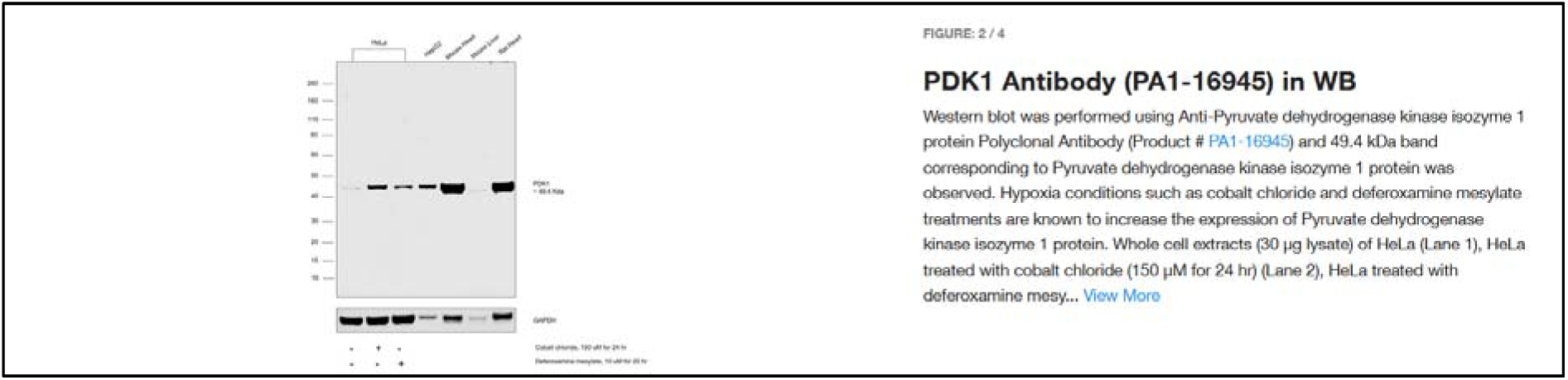
Screenshot of the webpage. Information on antibody website regarding pyruvate dehydrogenase kinase 1.

**Figure S23.**
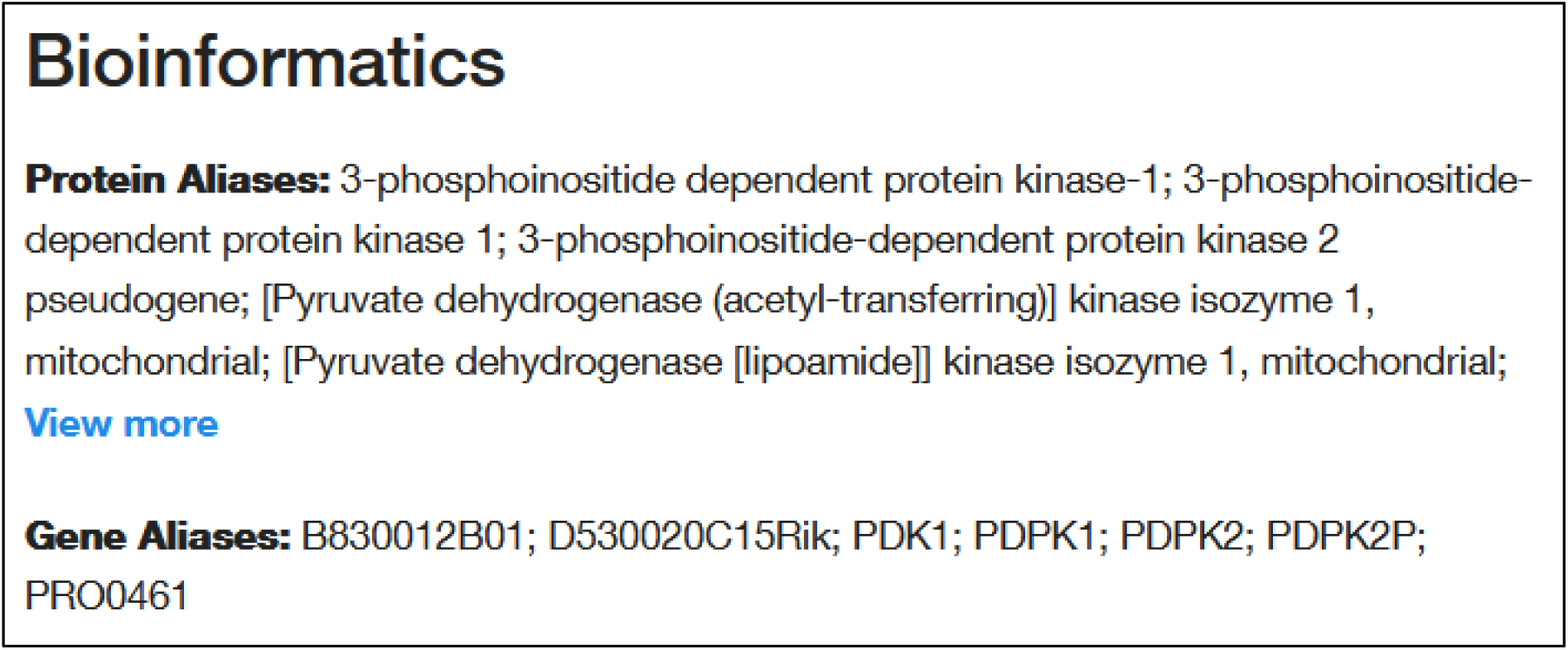
Screenshot of the webpage. Information on antibody website regarding both 3-phosphoinositide-dependent protein kinase 1 and pyruvate dehydrogenase kinase 1.

#### PDK1 Polyclonal Antibody [#PA5-115732]

Date of website access: 22.7.2025

Website link: https://www.thermofisher.com/antibody/product/PDK1-Antibody-Polyclonal/PA5-115732

Antibody website provides target information where both pyruvate dehydrogenase kinase 1 and 3-phosphoinositide-dependent protein kinase 1 are referenced as targets (**Figure S24 and S25**).

**Figure S24.**
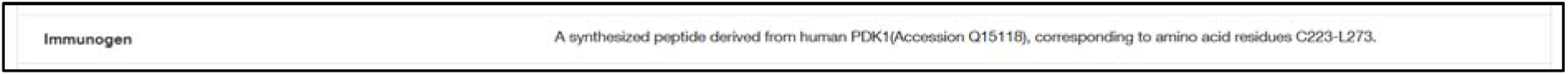
Screenshot of the webpage. Information on antibody website regarding pyruvate dehydrogenase kinase 1; Q15118 corresponds to pyruvate dehydrogenase kinase 1.

**Figure S25.**
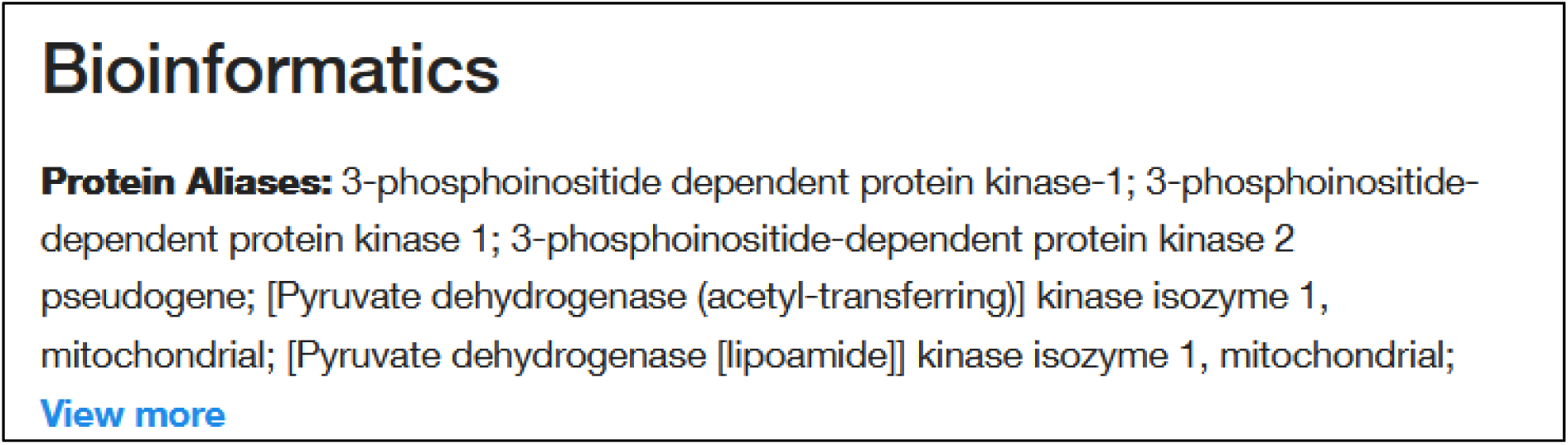
Screenshot of the webpage. Information on antibody website regarding both 3-phosphoinositide-dependent protein kinase 1 and pyruvate dehydrogenase kinase 1.

#### PDK1 Polyclonal Antibody [#PA5-106712]

Date of website access: 22.7.2025

Website link: https://www.thermofisher.com/antibody/product/PDK1-Antibody-Polyclonal/PA5-106712

Antibody website provides target information where both pyruvate dehydrogenase kinase 1 and 3-phosphoinositide-dependent protein kinase 1 are referenced as targets (**Figure S26 and S27**).

**Figure S26.**
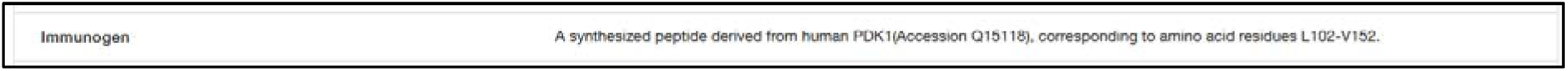
Screenshot of the webpage. Information on antibody website regarding pyruvate dehydrogenase kinase 1; Q15118 corresponds to pyruvate dehydrogenase kinase 1.

**Figure S27.**
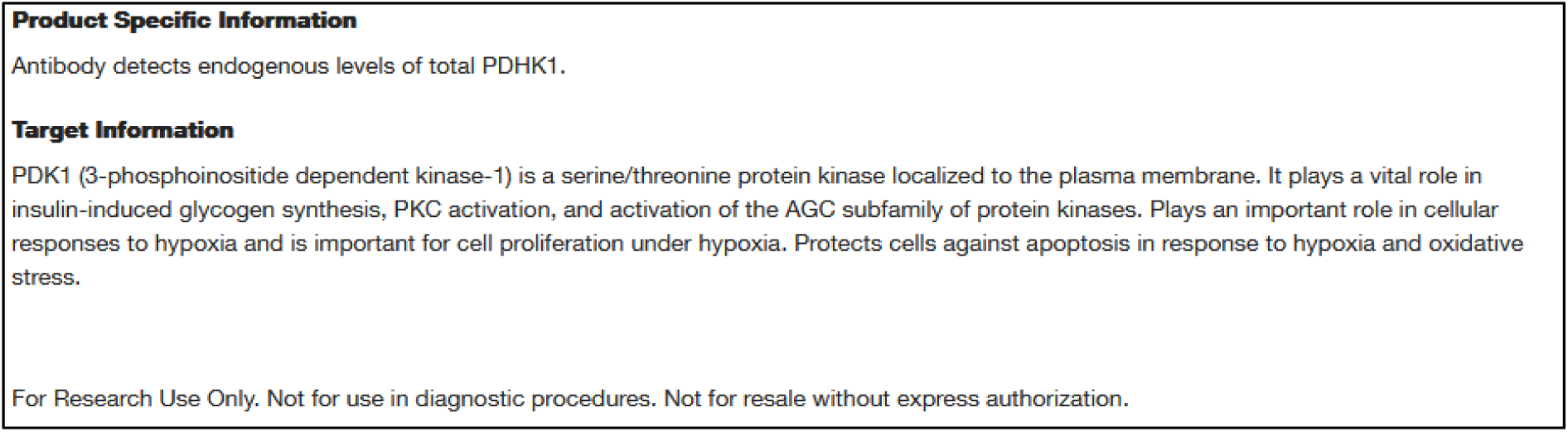
Screenshot of the webpage. Information on antibody website regarding both 3-phosphoinositide-dependent protein kinase 1 and pyruvate dehydrogenase kinase 1.

#### PDK1 Polyclonal Antibody [#PA5-28342]

Date of website access: 22.7.2025

Website link: https://www.thermofisher.com/antibody/product/PDK1-Antibody-Polyclonal/PA5-28342 This antibody was tested in our article.

Antibody website provides target information where both pyruvate dehydrogenase kinase 1 and 3-phosphoinositide-dependent protein kinase 1 are referenced as targets (**Figure S28 and S29**).

**Figure S28.**
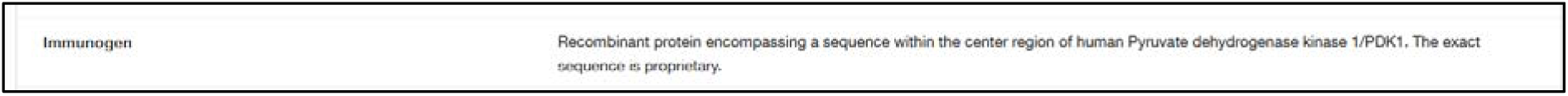
Screenshot of the webpage. Information on antibody website regarding pyruvate dehydrogenase kinase 1.

**Figure S29.**
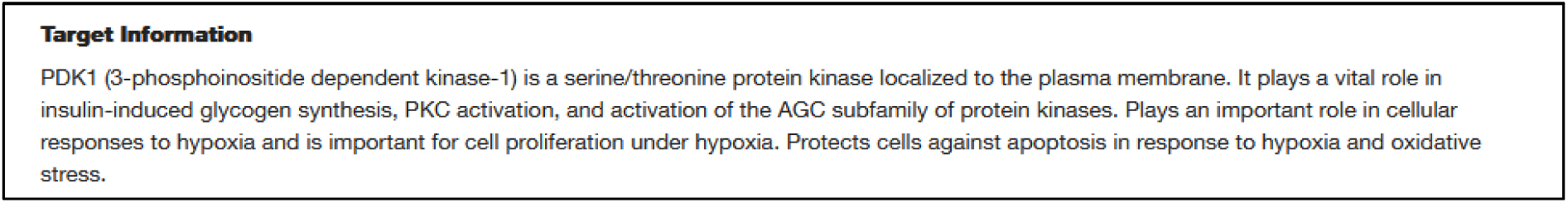
Screenshot of the webpage. Information on antibody website regarding 3-phosphoinositide-dependent protein kinase 1.

#### PDK1 Monoclonal Antibody (3B7) [#H00005163-M01]

Date of website access: 22.7.2025

Website link: https://www.thermofisher.com/antibody/product/PDK1-Antibody-clone-3B7-Monoclonal/H00005163-M01

Antibody website provides target information where pyruvate dehydrogenase kinase 1 corresponding sequence: (GGKGKGSPSHRKHIGSINPNCNVLEVIKDGYENARRLCDLYYINSPELELEELNAKSPGQPIQVVYVPSHLYHMVFELFKNAMRATMEHHANRGVYPPIQ); and 3-phosphoinositide-dependent protein kinase 1 are referenced (**Figure S30**).

**Figure S30.**
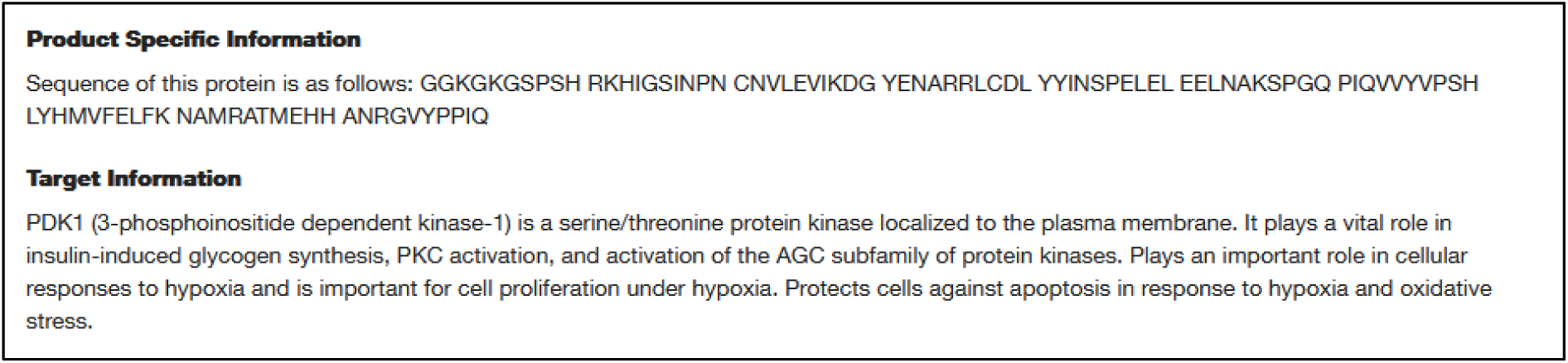
Screenshot of the webpage. Information on antibody website regarding pyruvate dehydrogenase kinase 1 corresponding sequence and 3-phosphoinositide-dependent protein kinase 1.

#### PDK1 Monoclonal Antibody (4A11) [#MA5-15797]

Date of website access: 23.7.2025

Website link: https://www.thermofisher.com/antibody/product/PDK1-Antibody-clone-4A11-Monoclonal/MA5-15797

Antibody website provides target information where both pyruvate dehydrogenase kinase 1 and 3-phosphoinositide-dependent protein kinase 1 are referenced as targets (**Figure S31**).

**Figure S31.**
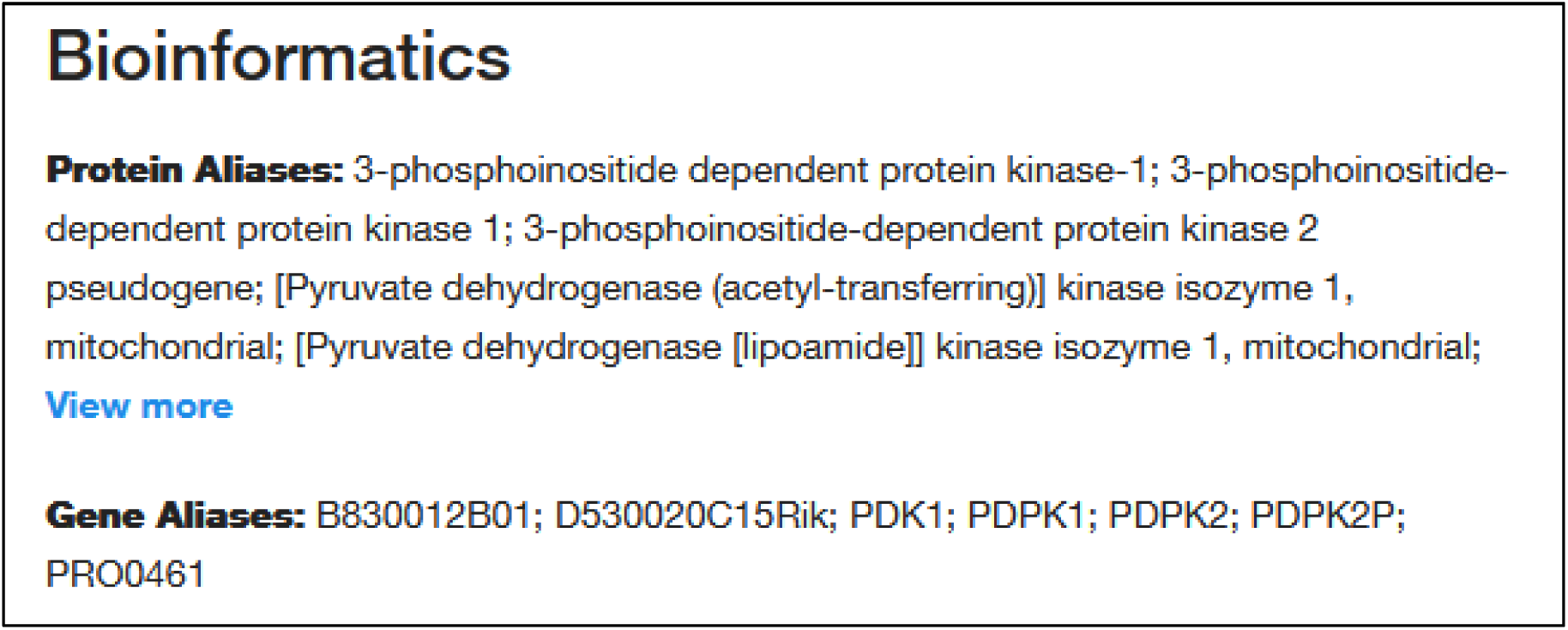
Screenshot of the webpage. Information on antibody website regarding both 3-phosphoinositide-dependent protein kinase 1 and pyruvate dehydrogenase kinase 1.

#### PDK1 Monoclonal Antibody (3E1) [#H00005163-M02]

Date of website access: 23.7.2025

Website link: https://www.thermofisher.com/antibody/product/PDK1-Antibody-clone-3E1-Monoclonal/H00005163-M02

Antibody website provides target information where pyruvate dehydrogenase kinase 1 corresponding sequence (same as stated for H00005163-M01 antibody, see 2.4.10. section):

(GGKGKGSPSHRKHIGSINPNCNVLEVIKDGYENARRLCDLYYINSPELELEELNAKSPGQPIQVVYVPSHLYHMVFELFKNAMRATMEHHANRGVYPPIQ); and 3-phosphoinositide-dependent protein kinase 1 are referenced (**Figure S32**).

**Figure S32.**
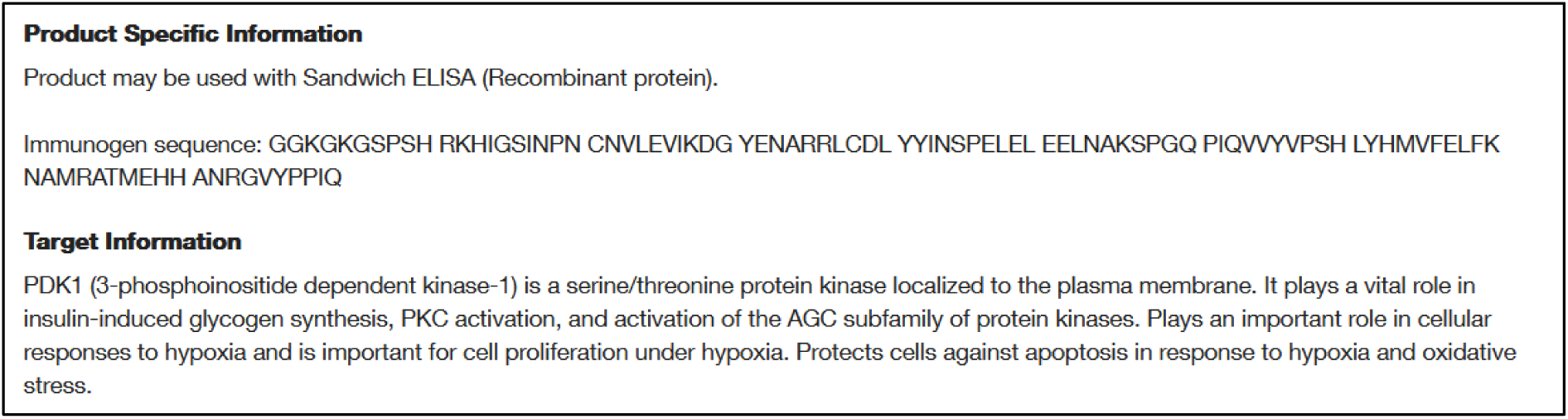
Screenshot of the webpage. Information on antibody website regarding pyruvate dehydrogenase kinase 1 corresponding sequence and 3-phosphoinositide-dependent protein kinase 1.

#### PDK1 Recombinant Rabbit Monoclonal Antibody (9A7X4) [#MA5-55304]

Date of website access: 23.7.2025

Website link: https://www.thermofisher.com/antibody/product/PDK1-Antibody-clone-9A7X4-Recombinant-Monoclonal/MA5-55304

Antibody website provides target information where both 3-phosphoinositide-dependent protein kinase 1 and pyruvate dehydrogenase kinase 1 are referenced as targets (**Figure S33**).

**Figure S33.**
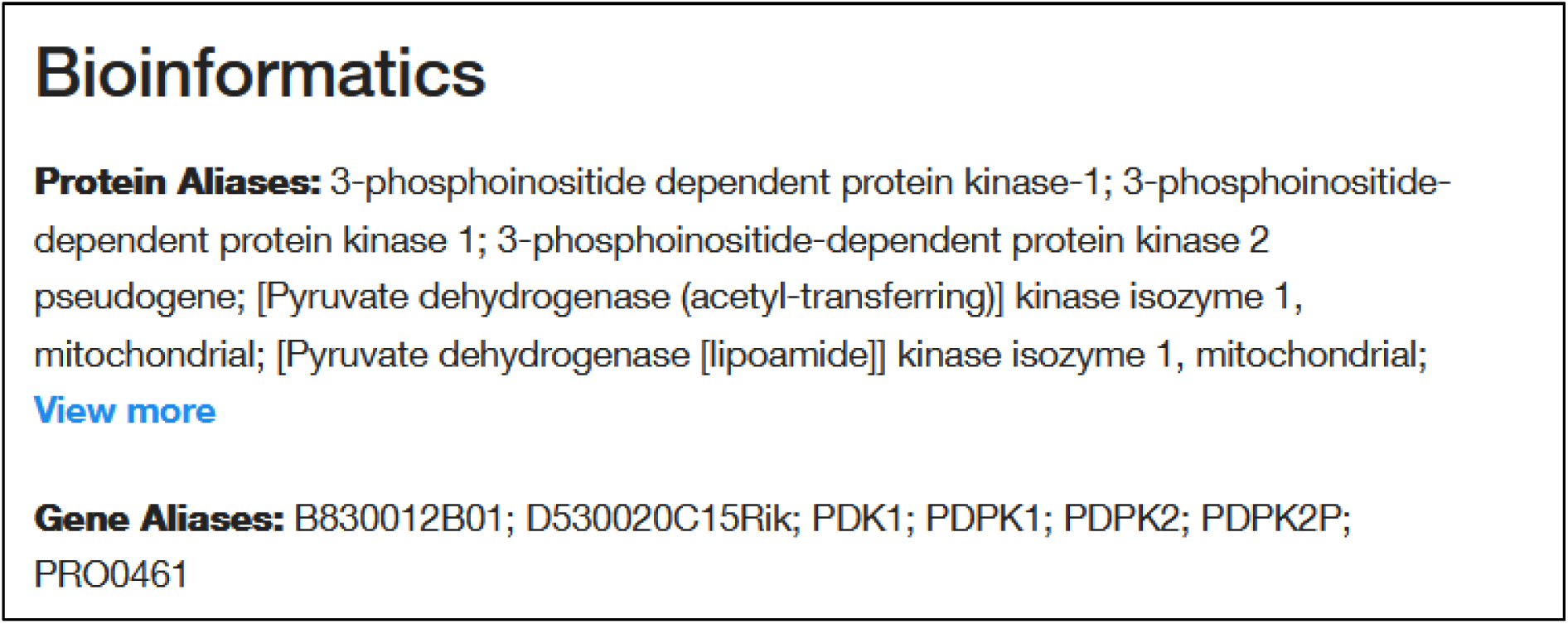
Screenshot of the webpage. Information on antibody website regarding both 3-phosphoinositide-dependent protein kinase 1 and pyruvate dehydrogenase kinase 1.

#### PDK1 Recombinant Rabbit Monoclonal Antibody (5T6S8) [#MA5-35736]

Date of website access: 23.7.2025

Website link: https://www.thermofisher.com/antibody/product/PDK1-Antibody-clone-5T6S8-Recombinant-Monoclonal/MA5-35736

Antibody website provides target information where pyruvate dehydrogenase kinase 1 corresponding sequence: (STAPRPRVETSRAVPLAGFGYGLPISRLYAQYFQGDLKLYSLEGYGTDAVIYIKALSTDSIERLPVYNKAA WKHYNTNHEADDWCVPSREPKDMTTFRSA); and 3-phosphoinositide-dependent protein kinase 1 are referenced (**Figure S34**).

**Figure S34.**
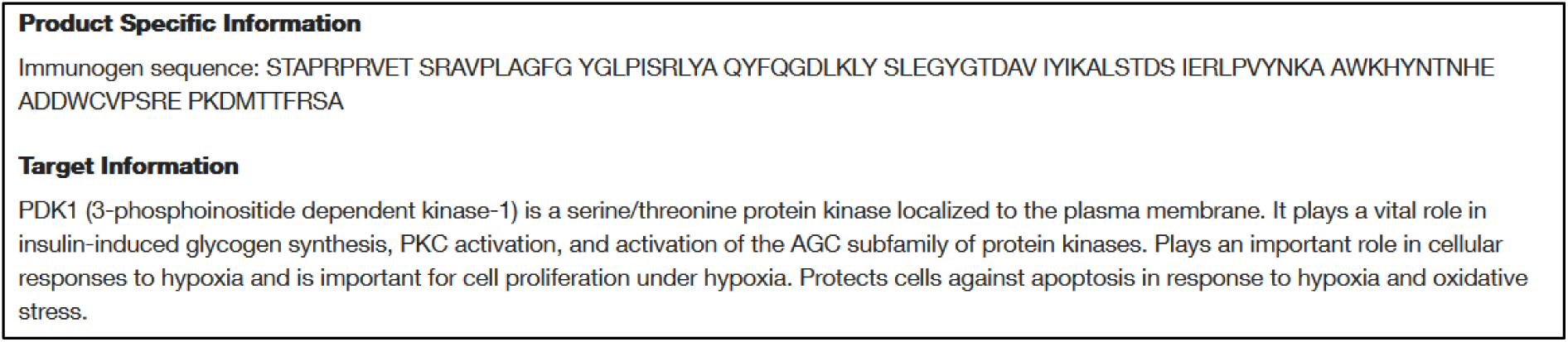
Screenshot of the webpage. Information on antibody website regarding pyruvate dehydrogenase kinase 1 corresponding sequence and 3-phosphoinositide-dependent protein kinase 1.

#### PDK1 Recombinant Rabbit Monoclonal Antibody (8J7H4) [#MA5-55303]

Date of website access: 23.7.2025

Website link: https://www.thermofisher.com/antibody/product/PDK1-Antibody-clone-8J7H4-Recombinant-Monoclonal/MA5-55303

Antibody website provides target information where pyruvate dehydrogenase kinase 1 corresponding sequence (same as stated for MA5-35736 antibody, see 2.4.14. section):

(STAPRPRVETSRAVPLAGFGYGLPISRLYAQYFQGDLKLYSLEGYGTDAVIYIKALSTDSIERLPVYNKAA WKHYNTNHEADDWCVPSREPKDMTTFRSA); and 3-phosphoinositide-dependent protein kinase 1 are referenced (Figure S35).

**Figure S35.**
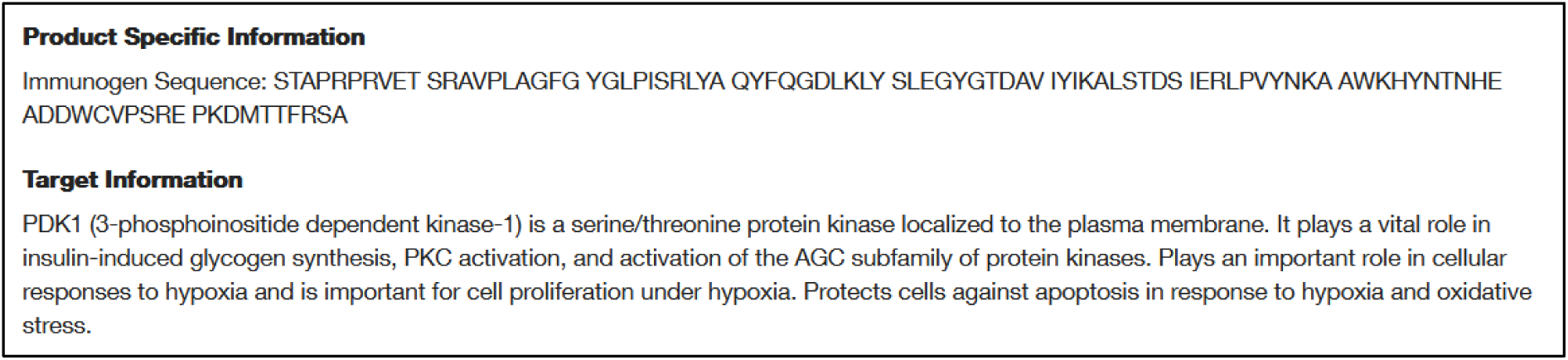
Screenshot of the webpage. Information on antibody website regarding pyruvate dehydrogenase kinase 1 corresponding sequence and 3-phosphoinositide-dependent protein kinase 1.

#### PDK1 Recombinant Rabbit Monoclonal Antibody (JA67-30) [#MA5-32702]

Date of website access: 23.7.2025

Website link: https://www.thermofisher.com/antibody/product/PDK1-Antibody-clone-JA67-30-Recombinant-Monoclonal/MA5-32702

Antibody website provides target information where both pyruvate dehydrogenase kinase 1 and 3-phosphoinositide-dependent protein kinase 1 are referenced as targets (**Figure S36 and S37**).

**Figure S36.**
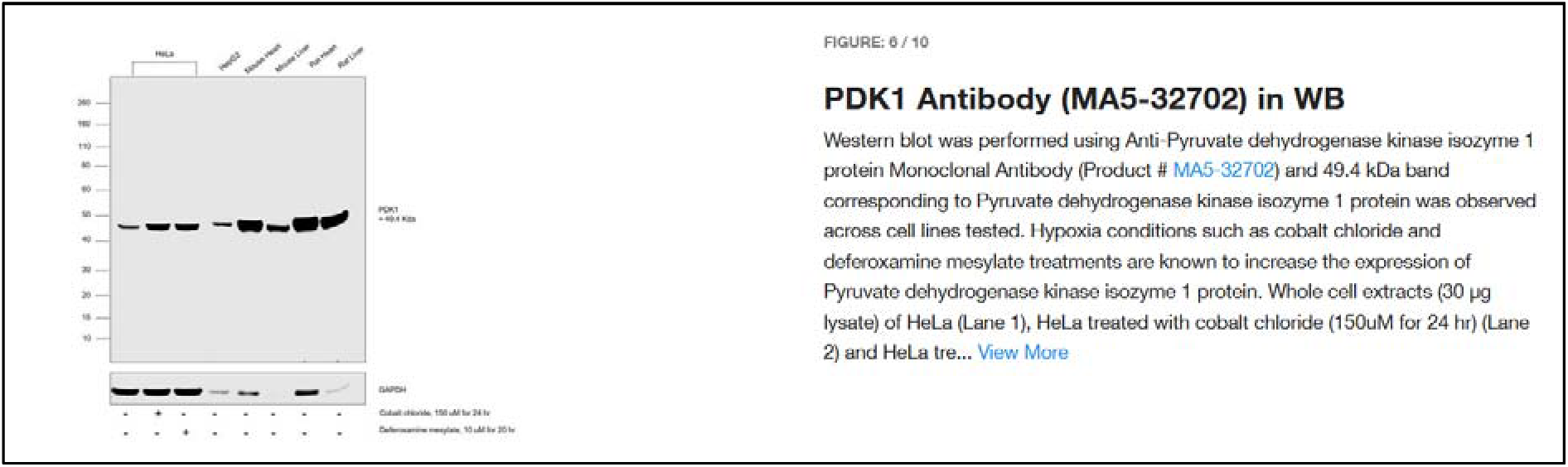
Screenshot of the webpage. Information on antibody website regarding pyruvate dehydrogenase kinase 1.

**Figure S37.**
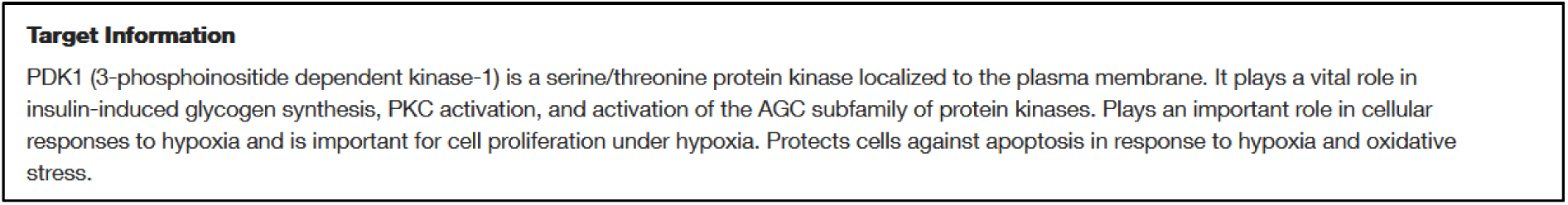
Screenshot of the webpage. Information on antibody website regarding 3-phosphoinositide-dependent protein kinase 1.

#### Human PDK1 (aa 225-254) Control Fragment Recombinant Protein [#RP-106913]

Date of website access: 23.7.2025

Website link: https://www.thermofisher.com/proteins/product/Human-PDK1-aa-225-254-Control-Fragment-Recombinant-Protein/RP-106913

Recombinant protein website provides information where pyruvate dehydrogenase kinase 1 corresponding sequence (**Figure S38**): (VLEVIKDGYENARRLCDLYYINSPELELEE); and 3-phosphoinositide-dependent protein kinase 1 are referenced (**Figure S39**).

**Figure S38.**
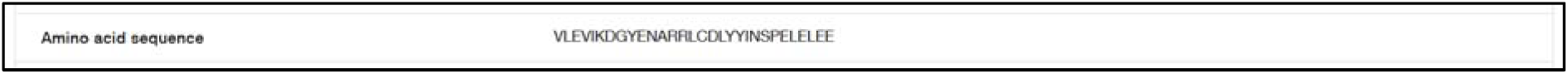
Screenshot of the webpage. Amino acid sequence corresponding with pyruvate dehydrogenase kinase 1.

**Figure S39.**
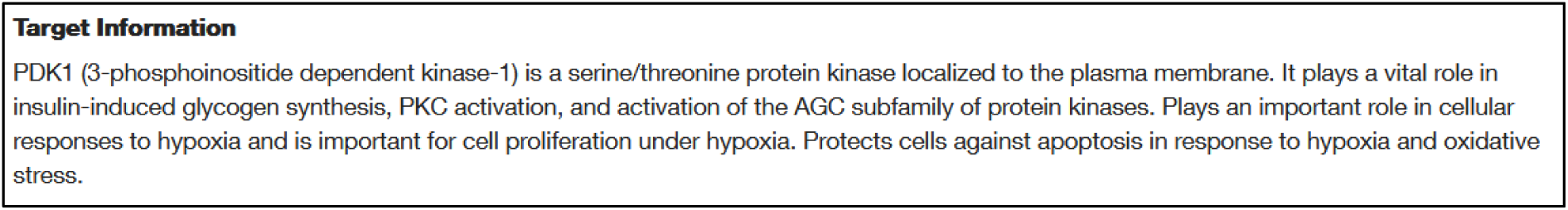
Screenshot of the webpage. Information on recombinant protein website regarding 3- phosphoinositide-dependent protein kinase 1. Note that this is stated under Target Information, even though this website is about recombinant protein and not an antibody.

#### Human PDK1 (aa 92-151) Control Fragment Recombinant Protein [#RP-95213]

Date of website access: 23.7.2025

Website link: https://www.thermofisher.com/proteins/product/Human-PDK1-aa-92-151-Control-Fragment-Recombinant-Protein/RP-95213

Recombinant protein website provides information where pyruvate dehydrogenase kinase 1 corresponding sequence (**Figure S40**):

**Figure S40.**
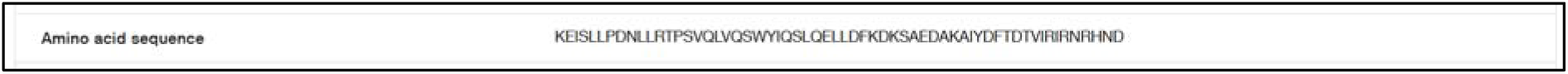
Screenshot of the webpage. Amino acid sequence corresponding with pyruvate dehydrogenase kinase 1.

(KEISLLPDNLLRTPSVQLVQSWYIQSLQELLDFKDKSAEDAKAIYDFTDTVIRIRNRHND); and 3-phosphoinositide-dependent protein kinase 1 are referenced (**Figure S41**).

**Figure S41.**
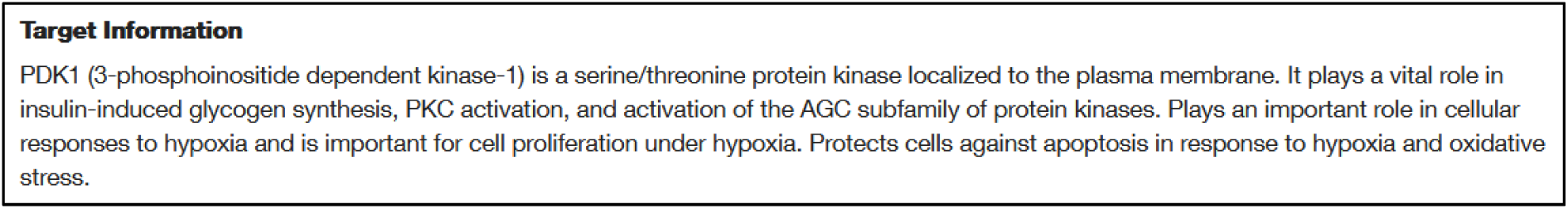
Screenshot of the webpage. Information on recombinant protein website regarding 3- phosphoinositide-dependent protein kinase 1. Note that this is stated under Target Information, even though this website is about recombinant protein and not an antibody.

#### PDK1 Polyclonal Antibody [#18262-1-AP]

Date of website access: 23.7.2025

Website link: https://www.thermofisher.com/antibody/product/PDK1-Antibody-Polyclonal/18262-1-AP Antibody website provides target information where pyruvate dehydrogenase kinase 1 corresponding sequence:

MRLARLLRGAALAGPGPGLRAAGFSRSFSSDSGSSPASERGVPGQVDFYARFSPSPLSMKQFLDFGSVNAC EKTSFMFLRQELPVRLANIMKEISLLPDNLLRTPSVQLVQSWYIQSLQELLDFKDKSAEDAKAIYDFTDTVIR IRNRHNDVIPTMAQGVIEYKESFGVDPVTSQNVQYFLDRFYMSRISIRMLLNQHSLLFGGKGKGSPSHRKHI GSINPNCNVLEVIKDGYENARRLCDLYYINSPELELEELNAKSPGQPIQVVYVPSHLYHMVFELFKNAMRA TMEHHANRGVYPPIQVHVTLGNEDLTVKMSDRGGGVPLRKIDRLFNYMYSTAPRPRVETSRAVPLAGFGY GLPISRLYAQYFQGDLKLYSLEGYGTDAVIYIKALSTDSIERLPVYNKAAWKHYNTNHEADDWCVPSREPK DMTTFRSA); and 3-phosphoinositide-dependent protein kinase 1 are referenced (**Figure S42**).

**Figure S42.**
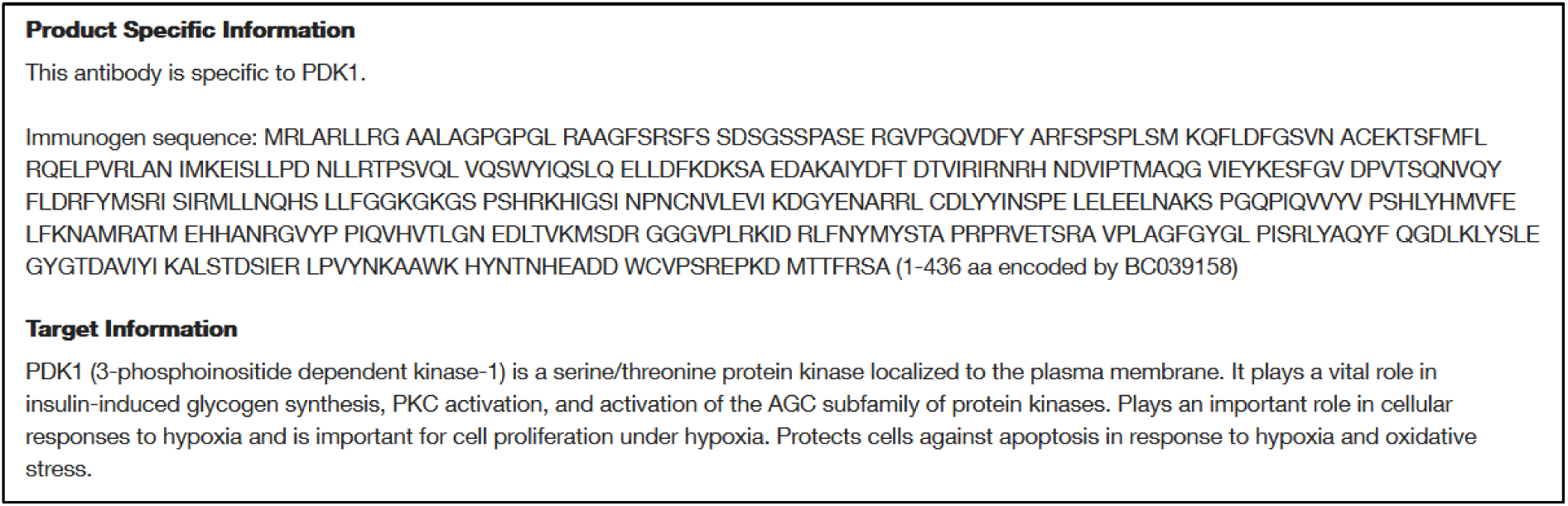
Screenshot of the webpage. Information on antibody website regarding pyruvate dehydrogenase kinase 1 corresponding sequence and 3-phosphoinositide-dependent protein kinase 1.

#### PDPK1 Polyclonal Antibody [#PA5-120471]

Date of website access: 23.7.2025

Website link: https://www.thermofisher.com/antibody/product/PDPK1-Antibody-Polyclonal/PA5-120471

Antibody website provides target information where both 3-phosphoinositide-dependent protein kinase 1 and pyruvate dehydrogenase kinase 1 (HGNC:8809 number corresponds to pyruvate dehydrogenase kinase 1, “lipoamide” term is sometimes included in the name pyruvate dehydrogenase *lipoamide* kinase 1) are referenced as targets (**Figure S43**).

**Figure S43.**
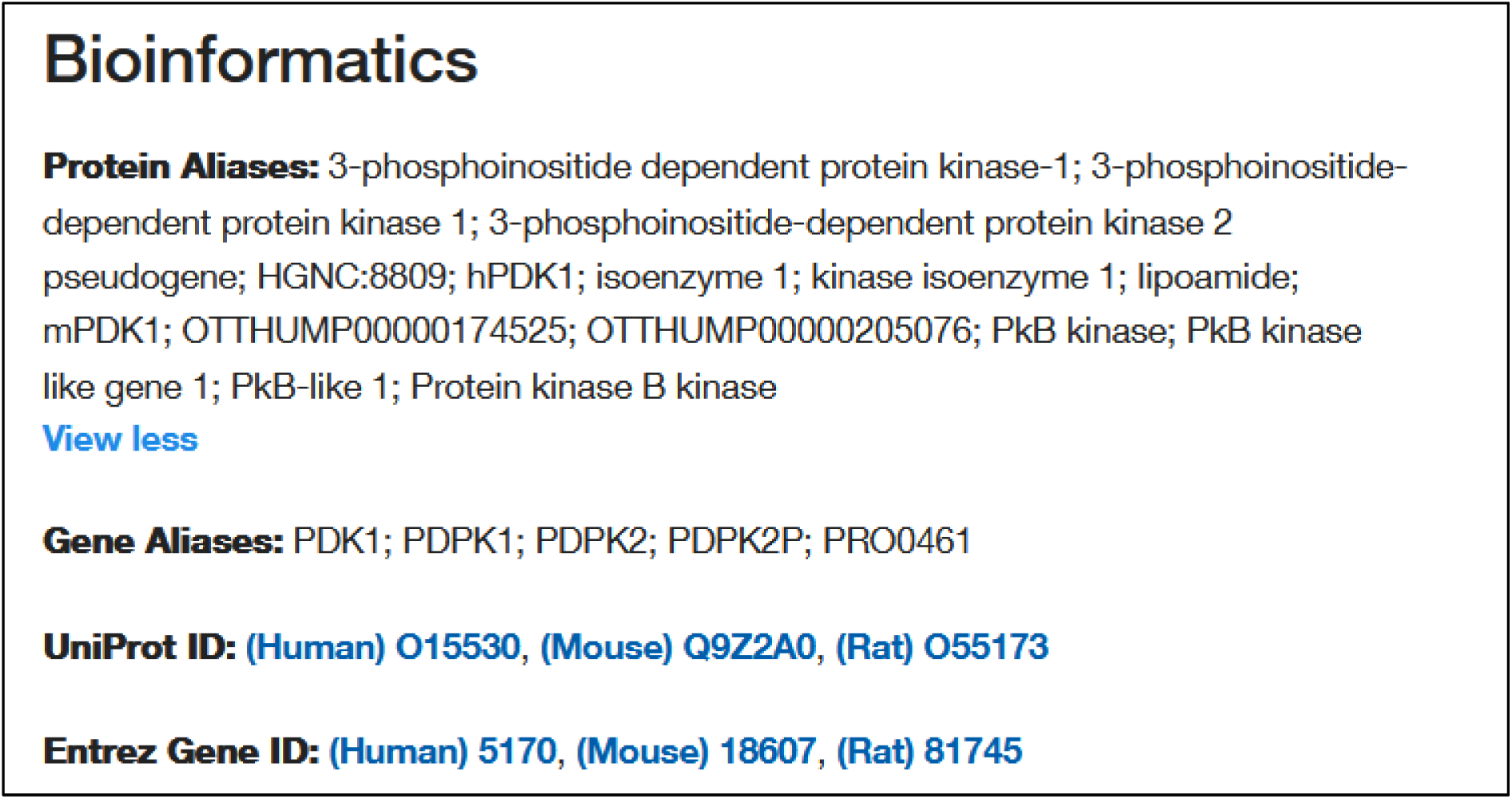
Screenshot of the webpage. Information on antibody website regarding 3-phosphoinositide-dependent protein kinase 1 and pyruvate dehydrogenase kinase 1; HGNC number 8809 and “lipoamide” term both regard the latter.

#### PDPK1 Polyclonal Antibody [#PA5-36071]

Date of website access: 23.7.2025

Website link: https://www.thermofisher.com/antibody/product/PDPK1-Antibody-Polyclonal/PA5-36071 This antibody was tested in our article.

Antibody website provides target information where both 3-phosphoinositide-dependent protein kinase 1 and pyruvate dehydrogenase kinase 1 (HGNC:8809 number corresponds to pyruvate dehydrogenase kinase 1, “lipoamide” term is sometimes included in the name pyruvate dehydrogenase *lipoamide* kinase 1) are referenced as targets (**Figure S44**).

**Figure S44.**
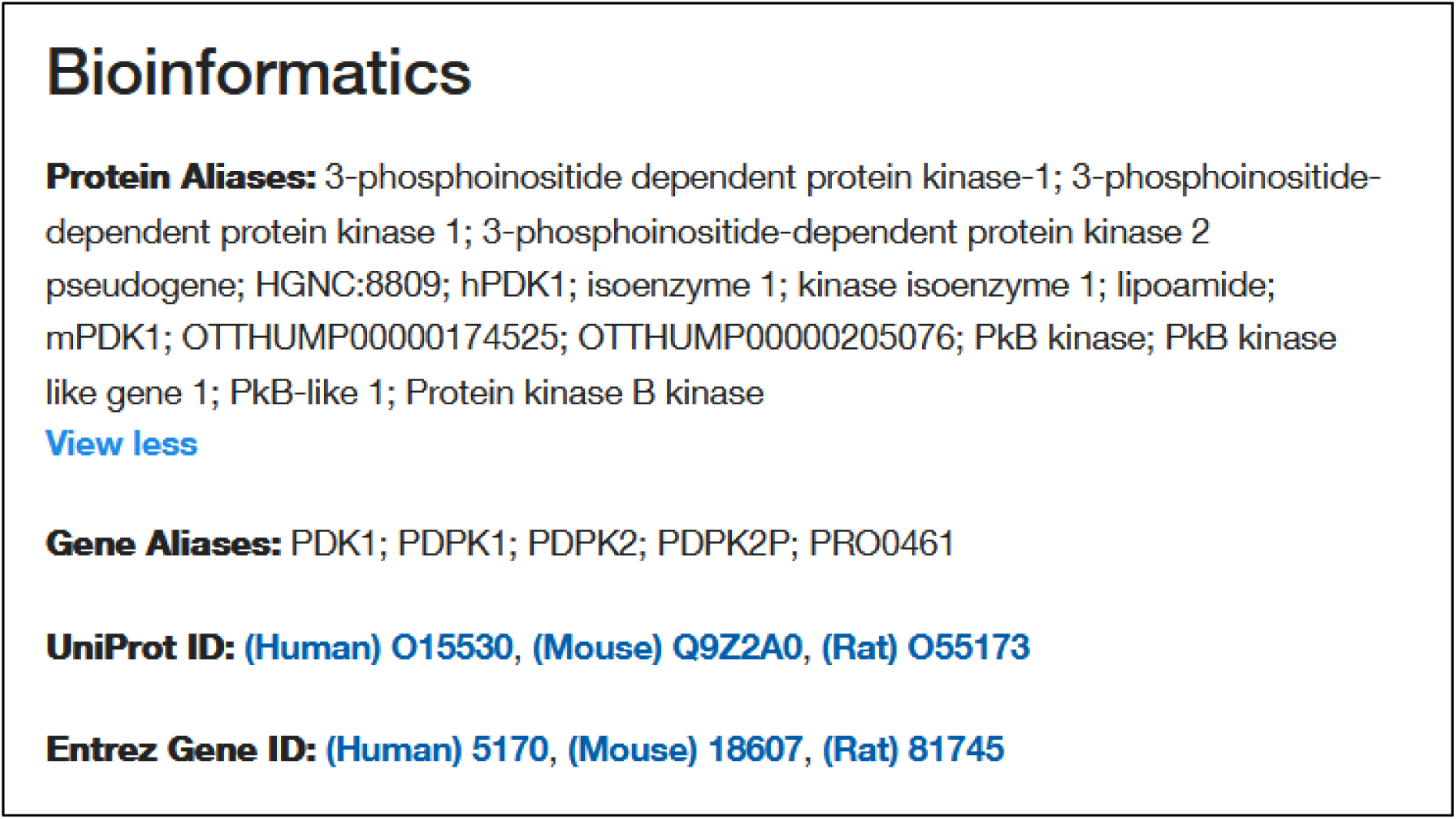
Screenshot of the webpage. Information on antibody website regarding 3-phosphoinositide-dependent protein kinase 1 and pyruvate dehydrogenase kinase 1; HGNC number 8809 and “lipoamide” term both regard the latter.

#### Phospho-PDPK1 (Ser396) Polyclonal Antibody [#PA5-12888]

Date of website access: 23.7.2025

Website link: https://www.thermofisher.com/antibody/product/Phospho-PDPK1-Ser396-Antibody-Polyclonal/PA5-12888

Antibody website provides target information where both 3-phosphoinositide-dependent protein kinase 1 and pyruvate dehydrogenase kinase 1 (HGNC:8809 number corresponds to pyruvate dehydrogenase kinase 1, “lipoamide” term is sometimes included in the name pyruvate dehydrogenase *lipoamide* kinase 1) are referenced as targets (**Figure S45**).

**Figure S45.**
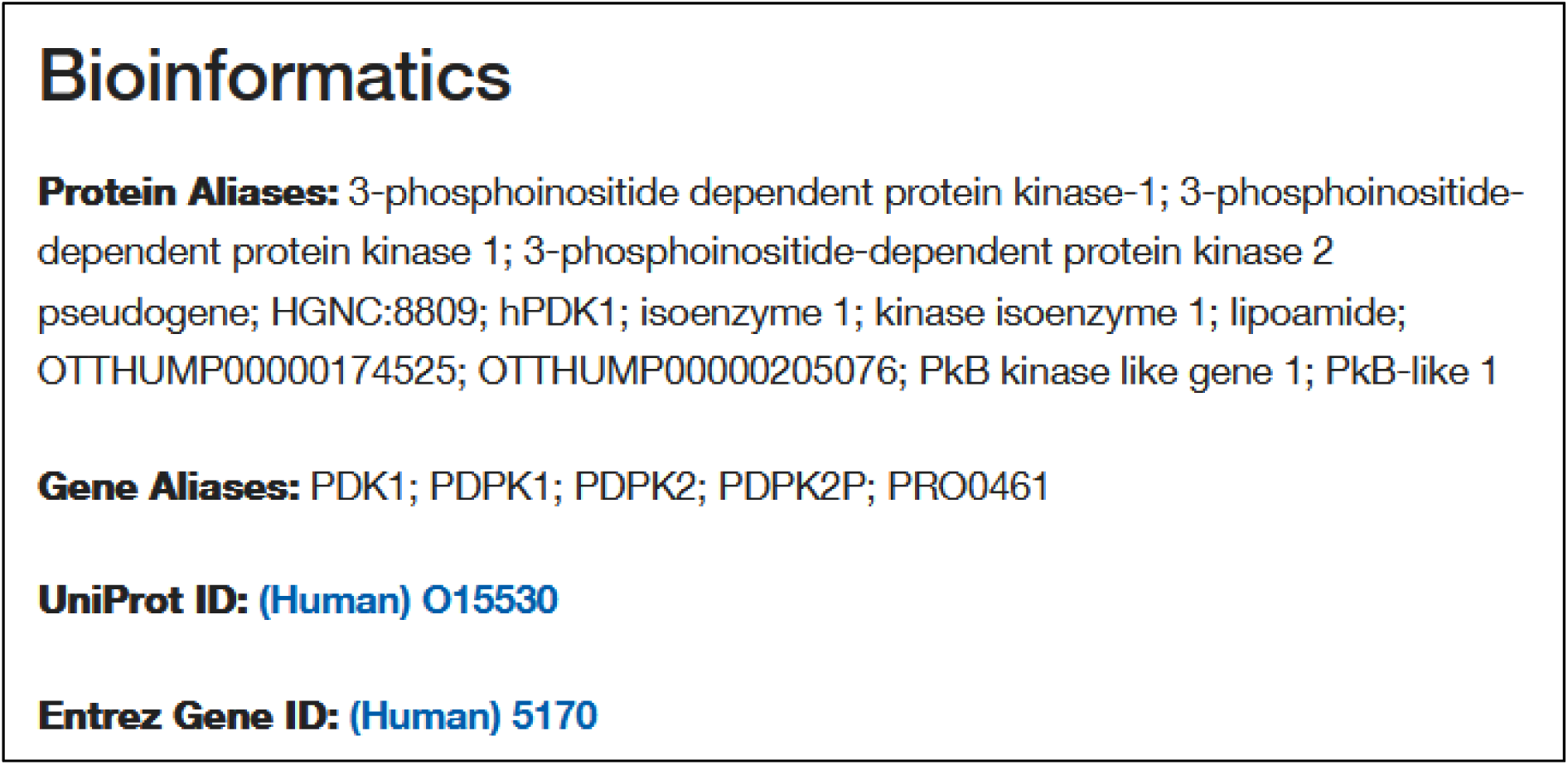
Screenshot of the webpage. Information on antibody website regarding 3-phosphoinositide-dependent protein kinase 1 and pyruvate dehydrogenase kinase 1; HGNC number 8809 and “lipoamide” term both regard the latter.

### Novus Biologicals

#### PDK-1 Antibody - BSA Free NBP2-44132

Date of website access: 19.6.2026

Website link: https://www.novusbio.com/products/pdk-1-antibody_nbp2-44132#datasheet

Antibody website provides target information where both 3-phosphoinositide-dependent protein kinase 1 (**Figure S46**) and pyruvate dehydrogenase kinase 1 (Uniprot: Q15118 corresponds to pyruvate dehydrogenase kinase 1) are referenced as targets (**Figure S47**).

**Figure S46.**
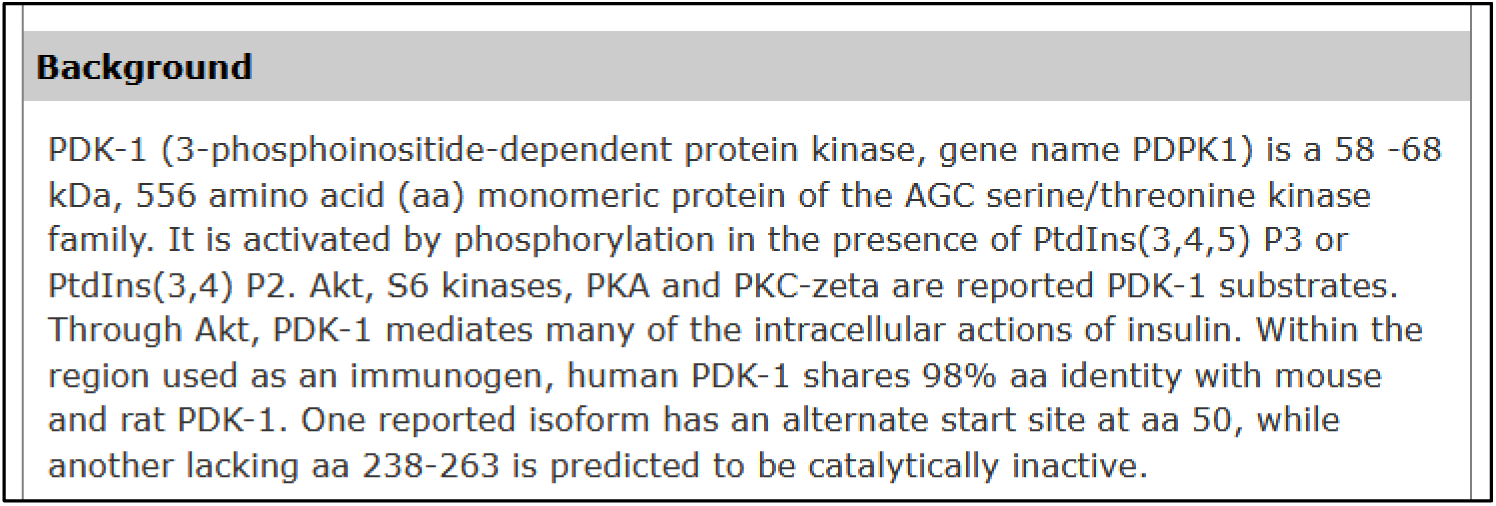
Screenshot of the webpage. Information on antibody website regarding 3-phosphoinositide-dependent protein kinase 1

**Figure S47.**
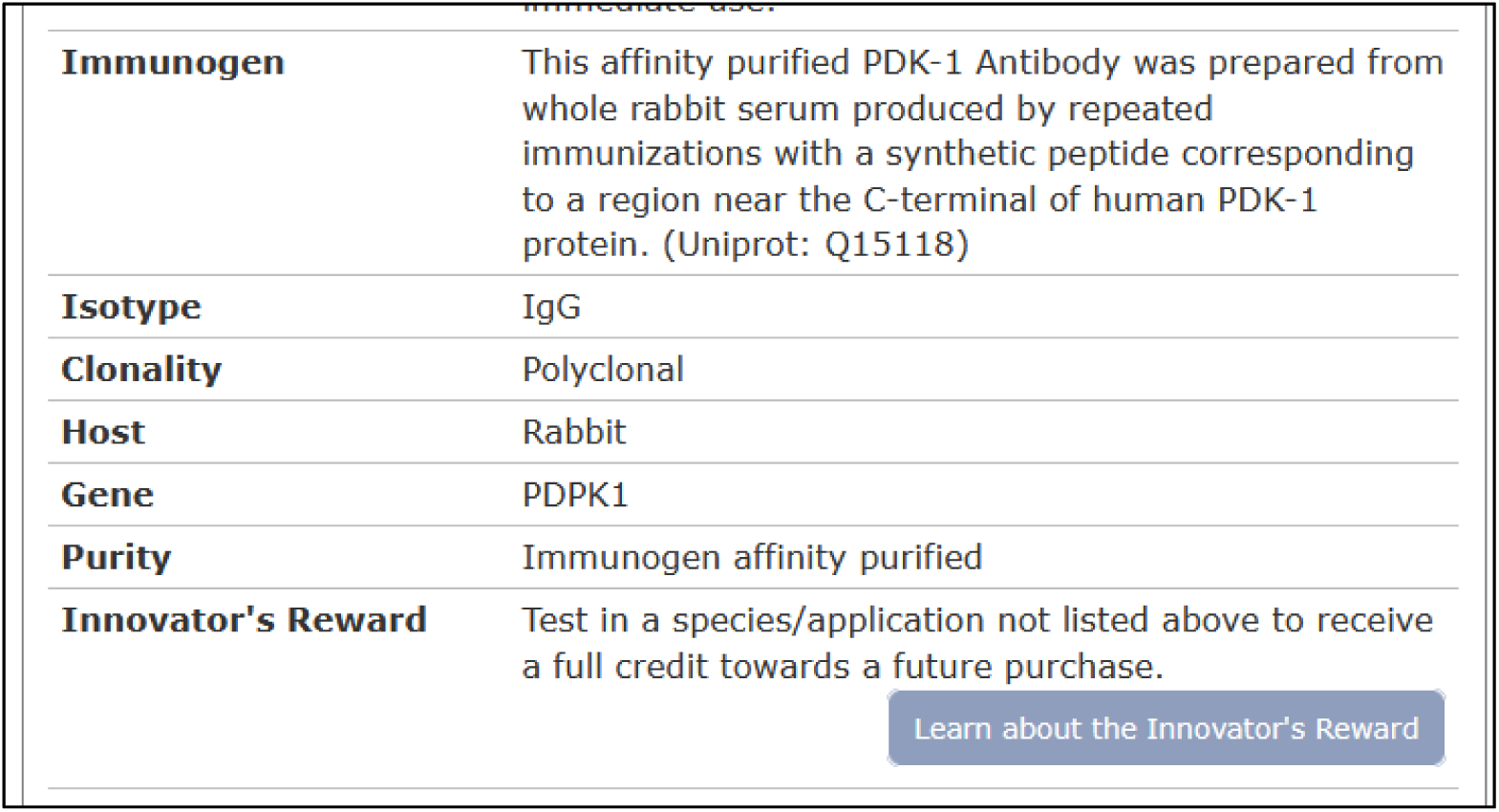
Screenshot of the webpage. Information on antibody website regarding pyruvate dehydrogenase kinase 1; Uniprot: Q15118 corresponds to pyruvate dehydrogenase kinase 1.

### Proteintech

#### PDK1 Polyclonal antibody 10026-1-AP

Date of website access: 19.6.2026

Website link: https://www.ptgcn.com/media/2671/akt-pathway.pdf https://www.ptglab.com/Products/Pictures/pdf/10026-1-AP.pdf

Poster provided by Proteintech presents AKT signaling pathway and related products. PDK1 protein is represented in the context of 3-phosphoinositide-dependent protein kinase 1 (as a kinase influenced by PI3K/PIP3 and affecting AKT) (**Figure S48**), but recommended antibody has a designated target of pyruvate dehydrogenase kinase 1 (**Figure S49**), as per 10026-1-AP Datasheet (**Figure S50**).

**Figure S48.**
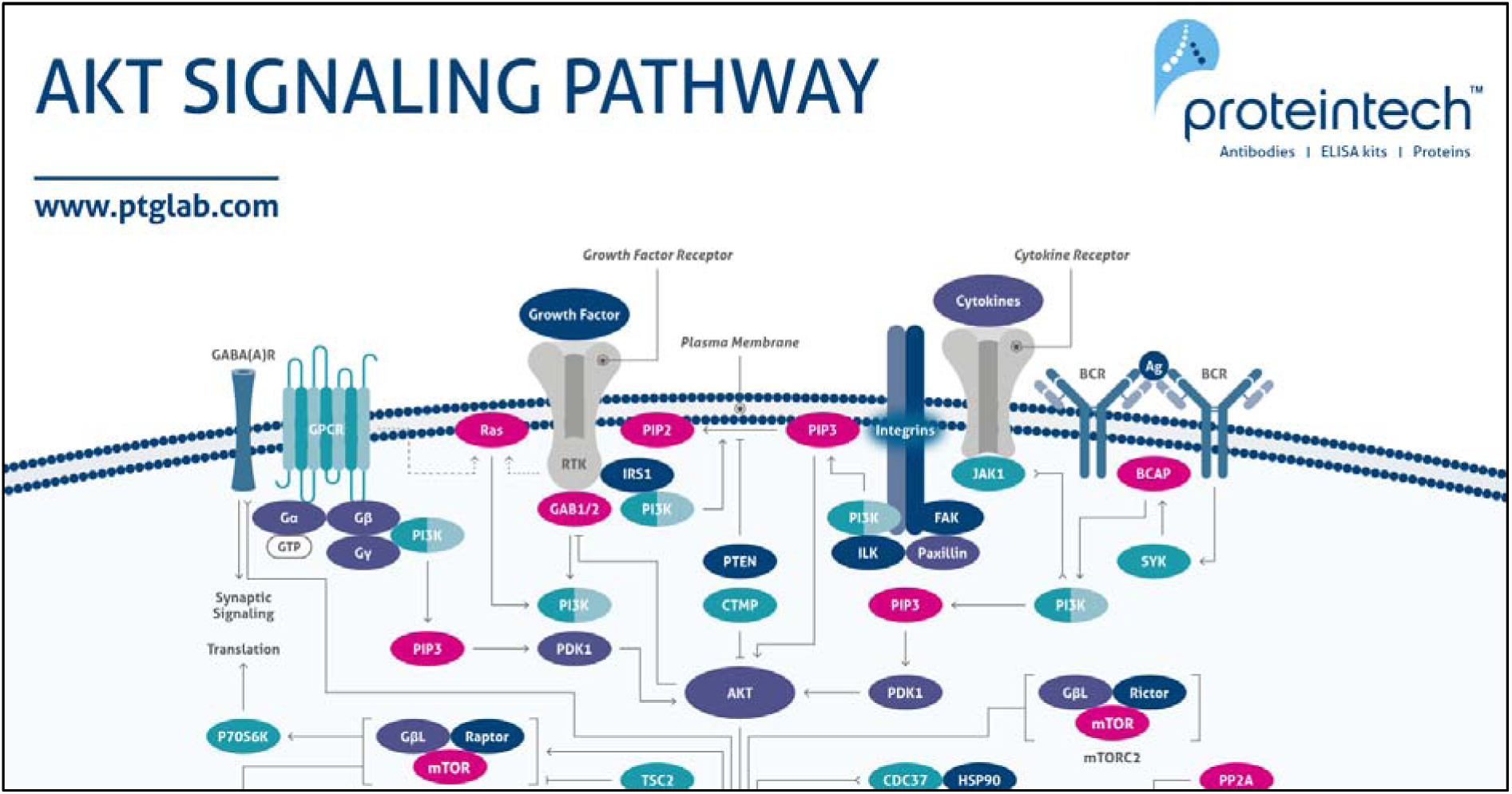
Screenshot of the poster. Information on antibody website regarding 3-phosphoinositide-dependent protein kinase 1

**Figure S49.**
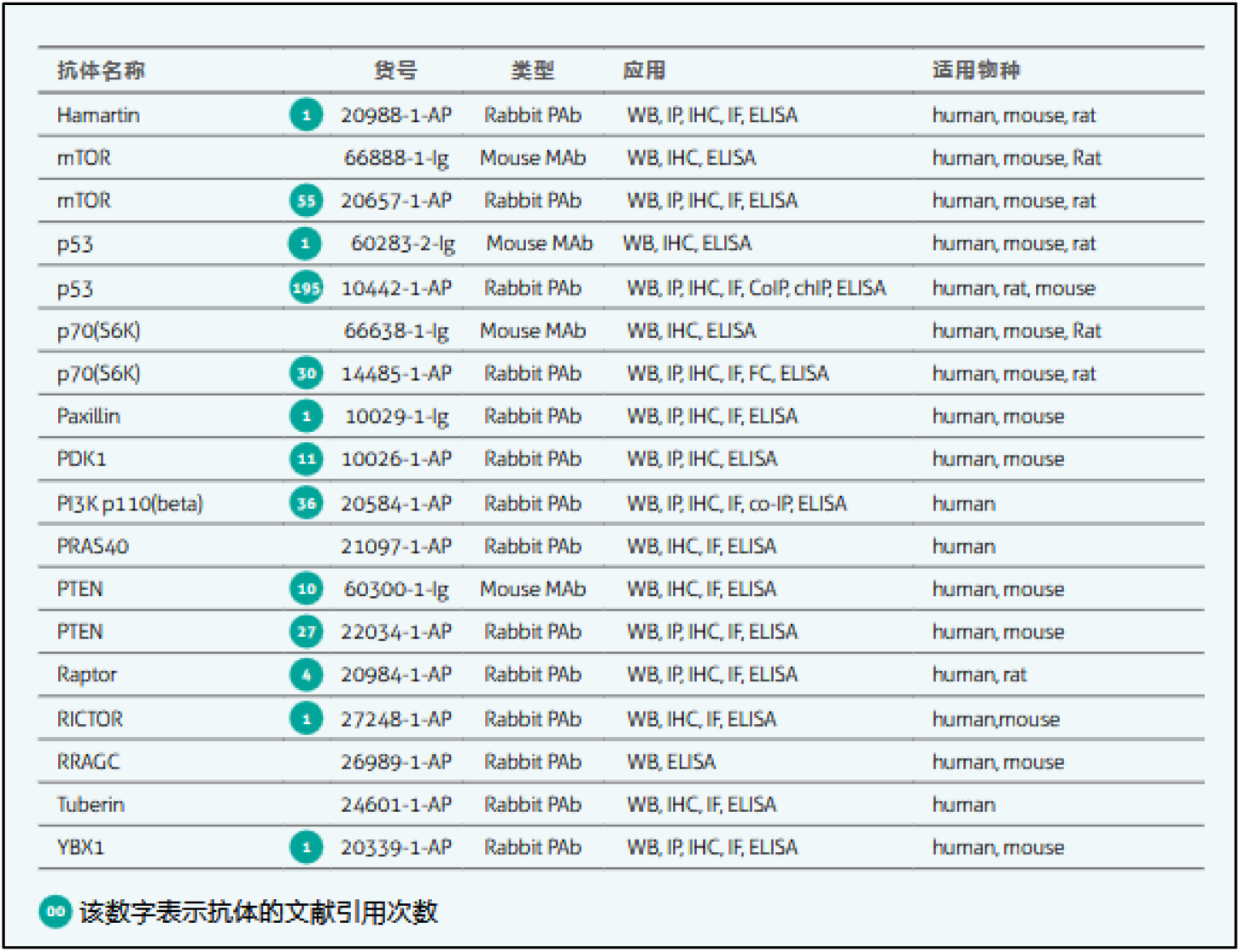
Screenshot of the poster. Information about recommended antibody.

**Figure S50.**
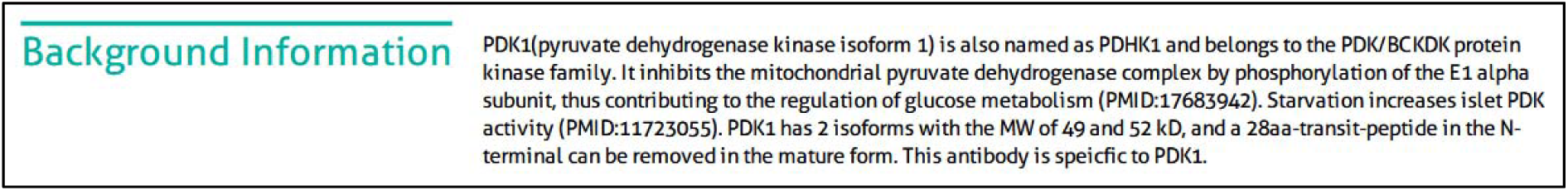
Screenshot of the poster. Information about recommended antibody.

## S7 Western blot raw data

**Figure S51.**
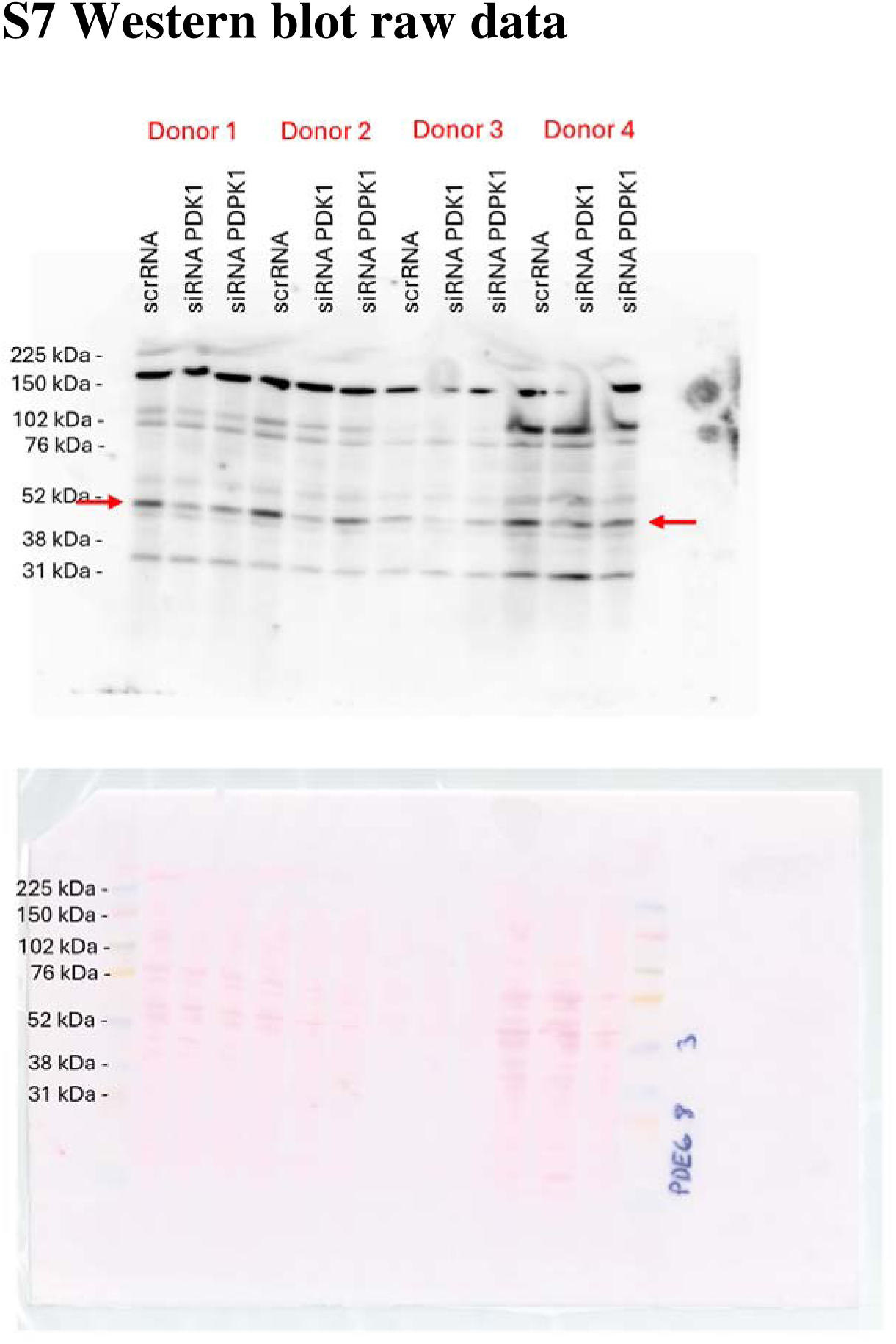
Western blot result using with PDHK1 (C47H1) Rabbit mAb #3820 and Ponceau S stained membrane for donors 1-4.

**Figure S52.**
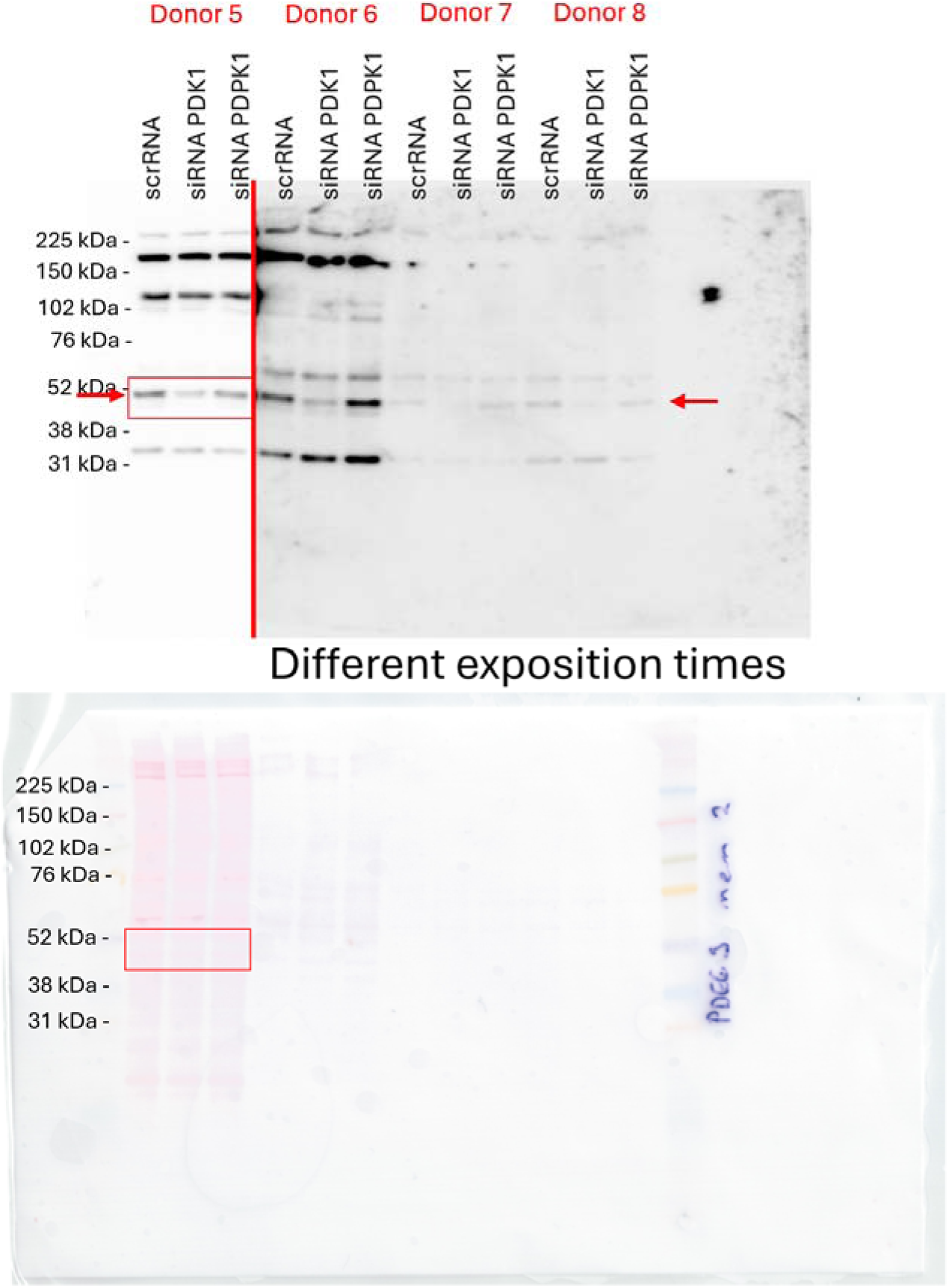
Western blot result using with PDHK1 (C47H1) Rabbit mAb #3820 and Ponceau S stained membrane for donors 5-8. Red boxes highlight which blot and membrane are shown on Figure 4C in the main article. Vertical red line divides western blot result between different exposition times.

**Figure S53.**
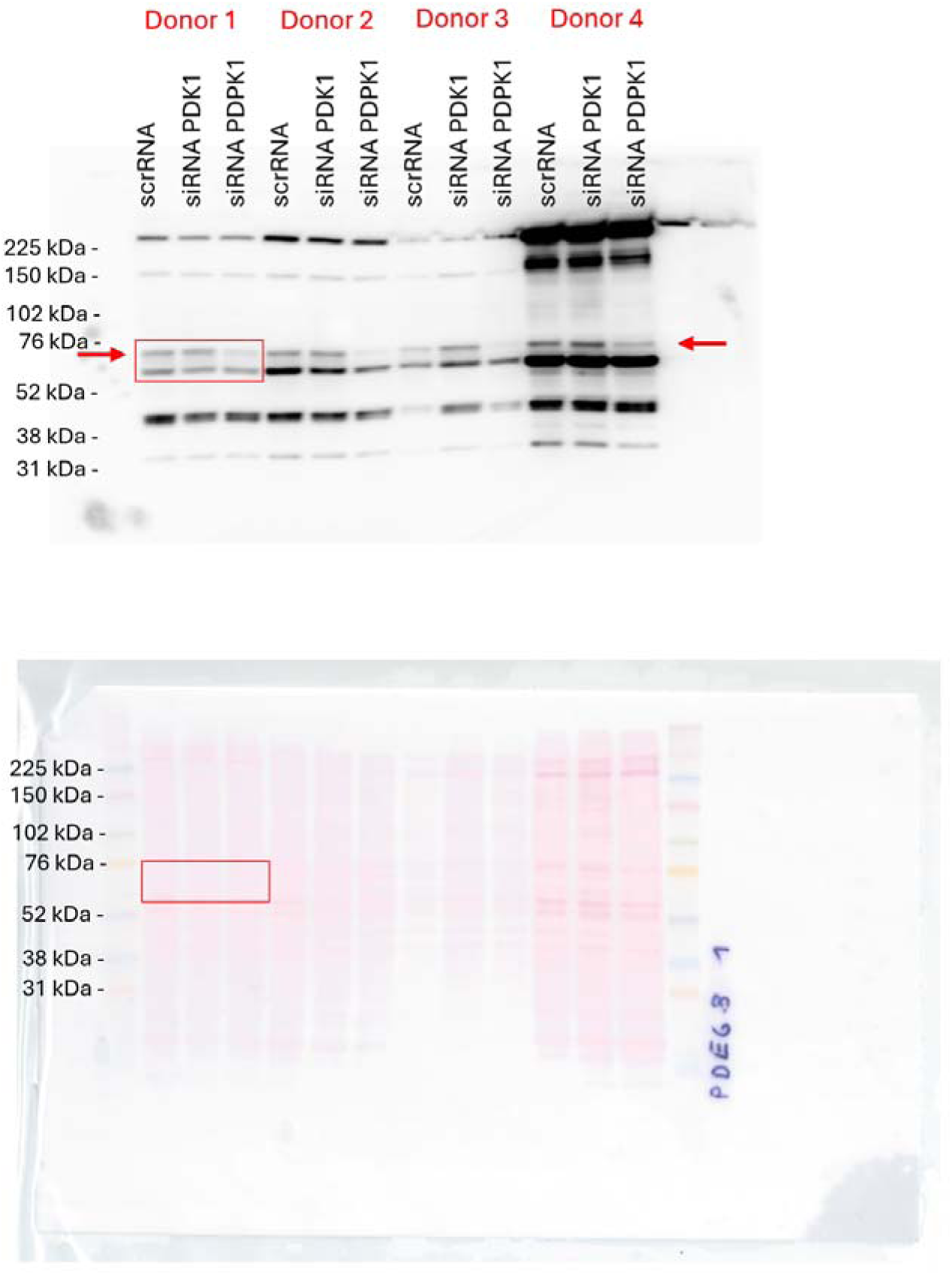
Western blot result using PDK1 Antibody #3062 and Ponceau S stained membrane for donors 1-4. Red boxes highlight which blot and membrane are shown on Figure 4C in the main article.

**Figure S54.**
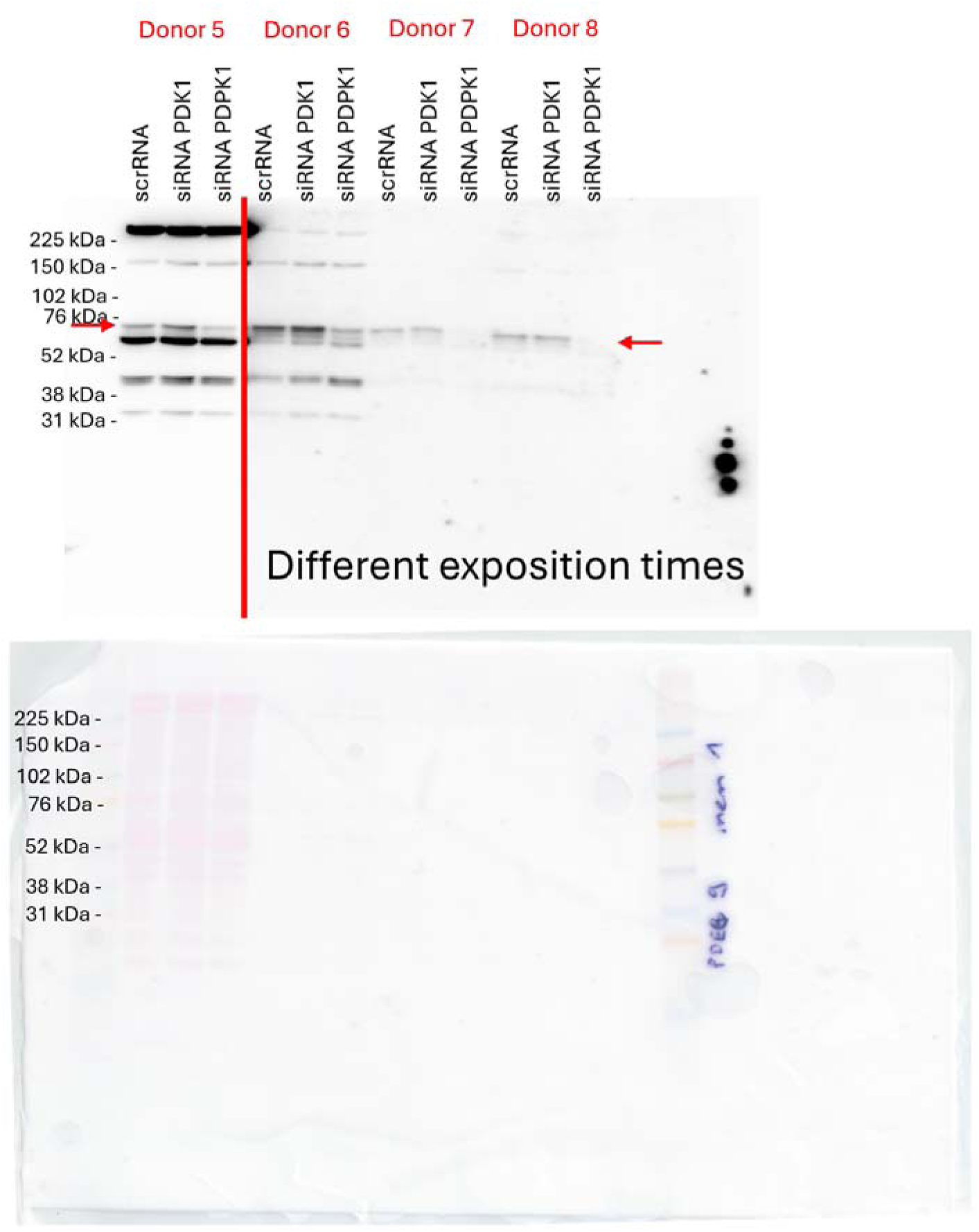
Western blot result using PDK1 Antibody #3062 and Ponceau S stained membrane for donors 5-8. Vertical red line divides western blot result between different exposition times.

**Figure S55.**
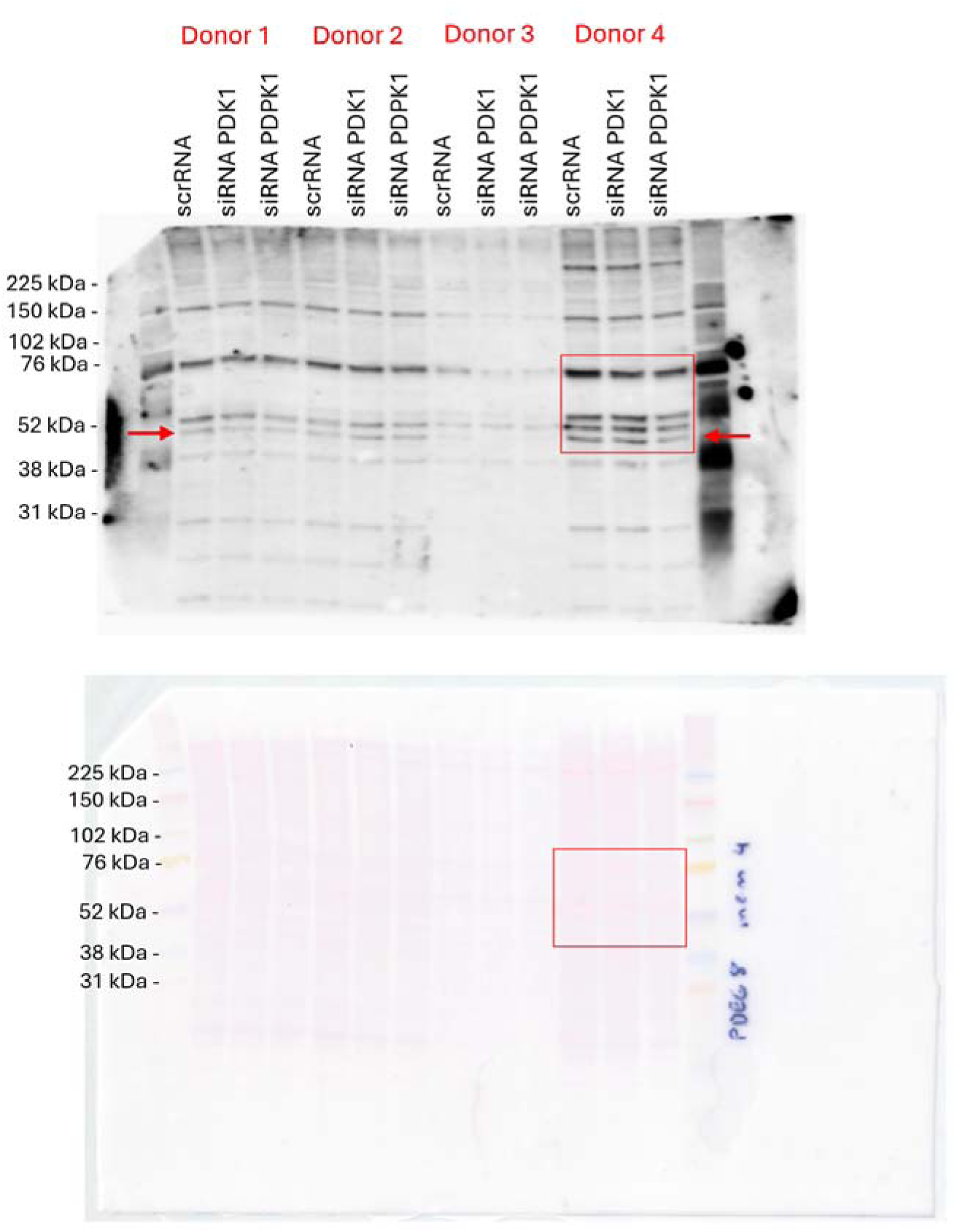
Western blot result using PDK1 Polyclonal Antibody PA5-28342 and Ponceau S stained membrane for donors 1-4. Red boxes highlight which blot and membrane are shown on Figure 4C in the main article.

**Figure S56.**
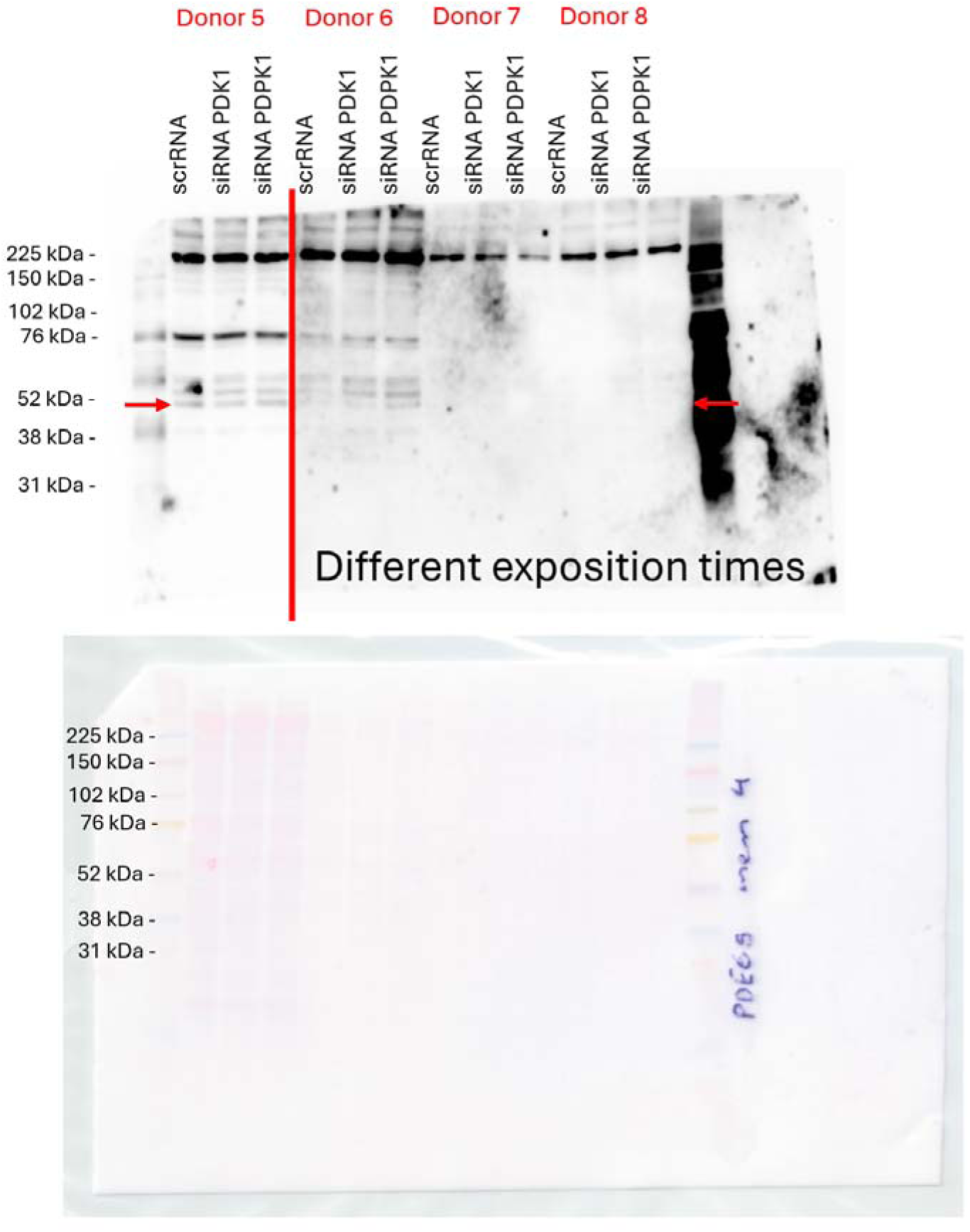
Western blot result using PDK1 Polyclonal Antibody PA5-28342 and Ponceau S stained membrane for donors 5-8. Vertical red line divides western blot result between different exposition times.

**Figure S57.**
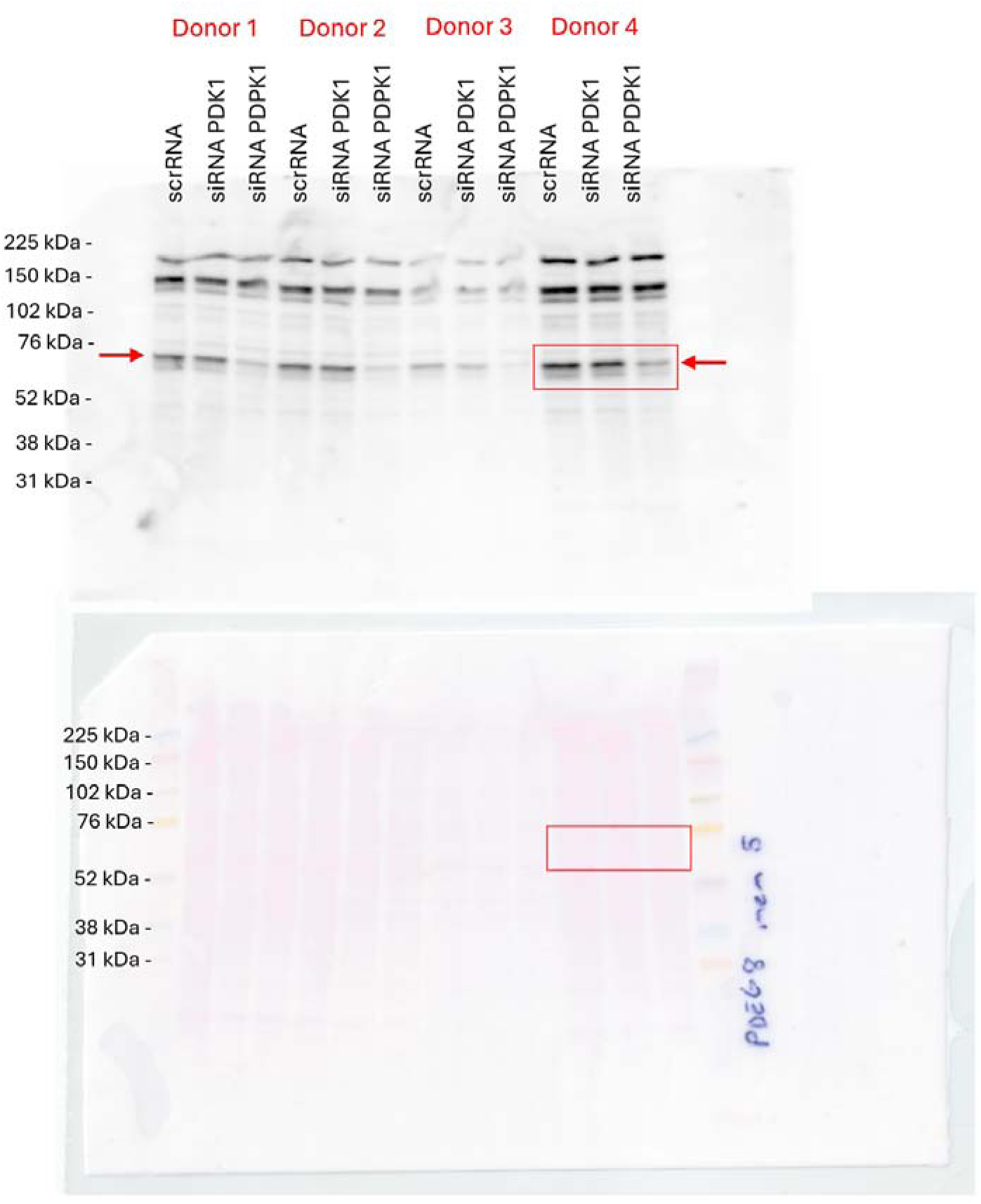
Western blot result using PDK1 Polyclonal Antibody PA5-36071 and Ponceau S stained membrane for donors 1-4. Red boxes highlight which blot and membrane are shown on Figure 4C in the main article.

**Figure S58.**
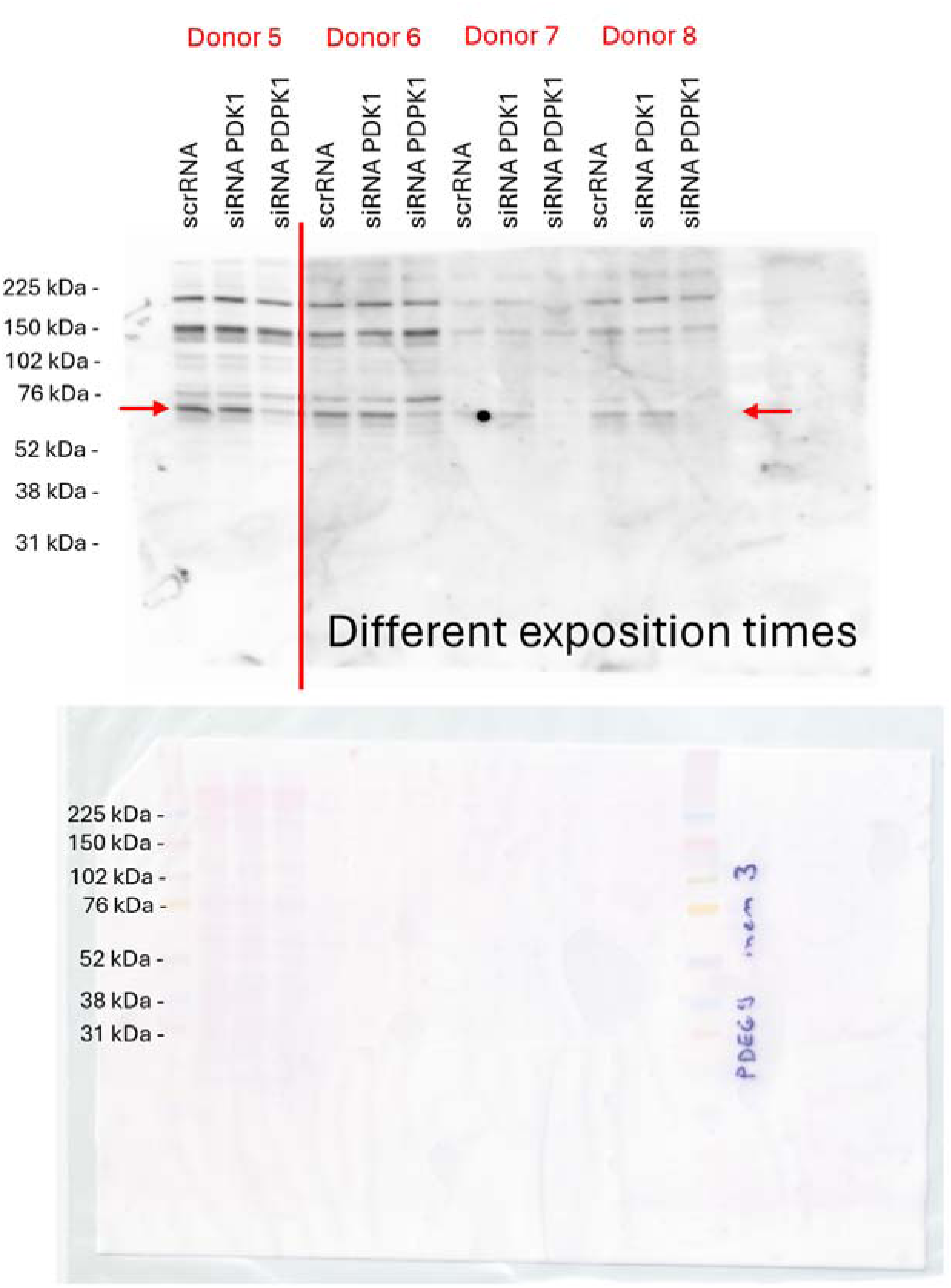
Western blot result using PDK1 Polyclonal Antibody PA5-36071 and Ponceau S stained membrane for donors 5-8. Vertical red line divides western blot result between different exposition times.

## Footnotes

1 Author discusses P16 abbreviation ambiguity.

2 One of these articles contained a PDK1-related error, which was curated, hence we treated this article as correct.

3 None of these retractions contained PDK1-related errors.

4 These include 25 retracted articles and retraction notices, 15 correction notices, 10 preprints, 2 unrelated articles, 16 articles published in Chinese, 10 articles focusing on PKD1 (Polycystic Kidney Disease 1) and not on PDK1, and 1 not accessible due to a paywall.

5 https://pdk1-tracker.ijs.si

6 Note that A~ includes error subtypes Ã 1 and Ã 2, and B̃ includes subtypes B~ 1 and B~ 2.

7 The UMAP technology used in the visualization is explained in the Supplement Methods related to citation network analysis.

8 The authors of the article (46); including an author of the present article; repeated the mRNA measurements using the proper primer.

9 Embeddings denote 2-dimensional projections of compressed vectors representing structural relationships of network nodes.

10 http://kt-nlp-demo.ijs.si:8080/pdk1/

11 Originally labelled by Kociper as B̃_C_, later reclassified as Ã_C_.

